# Resilient Living Materials Built By Printing Bacterial Spores

**DOI:** 10.1101/537571

**Authors:** Lina M. González, Christopher A. Voigt

**Affiliations:** Synthetic Biology Center, Department of Biological Engineering, Massachusetts Institute of Technology, Cambridge, MA, USA

**Keywords:** Synthetic Biology, Engineered Living Material, Genetic Circuit, Additive Manufacturing, 3D Printing

## Abstract

A route to advanced multifunctional materials is to embed them with living cells that can perform sensing, chemical production, energy scavenging, and actuation. A challenge in realizing this potential is that the conditions for keeping cells alive are not conducive to materials processing and require a continuous source of water and nutrients. Here, we present a 3D printer that can mix material and cell streams in a novel printhead and build 3D objects (up to 2.5 cm by 1 cm by 1 cm). Hydrogels are printed using 5% agarose, which has a low melting temperature (65°C) consistent with thermophilic cells, a rigid storage modulus (G’= 6.5 × 10^4^), exhibits shear thinning, and can be rapidly hardened upon cooling to preserve structural features. Spores of *B. subtilis* are printed within the material and germinate on its exterior, including spontaneously in cracks and new surfaces exposed by tears. By introducing genetically engineered bacteria, the materials can sense chemicals (IPTG, xylose, or vanillic acid). Further, we show that the spores are resilient to extreme environmental stresses, including desiccation, solvents (ethanol), high osmolarity (1.5 mM NaCl), 365 nm UV light, and γ-radiation (2.6 kGy). The construction of 3D printed materials containing spores enables the living functions to be used for applications that require long-term storage, in-field functionality, or exposure to uncertain environmental stresses.

## Introduction

Living materials, where cells are embedded within a structural scaffold, pervade the natural world, examples being wood, bone, and skin. The cells provide dynamic functions, including energy harvesting, repair, sensing, and actuation. Natural living materials can survive for years, even millennia, under fluctuating and stressful conditions. There has been an effort to design so-called engineered living materials, to artificially combine structural components with cells, including those that have been genetically engineered^1, 2^. Biomedical applications include the differentiation of human cells to grow artificial tissue or to serve in living medical devices ^3, 4, 5, 6^. The potential, however, is much greater across many applications, where cells are harnessed for their sensing^7^, chemical/material production^8^ self-healing^9^, self-powering^10^, self-cleaning^8^, and self-gluing^11, 12^ abilities. Natural microbes have been harnessed to build concrete and fix cracks^9, 13^, build packaging materials^11, 12^, and to control breathability in textiles by opening vents in response to sweat^14^. Engineered bacteria have been demonstrated to build metal wires within electronics^15, 16^, act as a pressure sensor within a device^16^, and degrade pollutants^17^. A key challenge for living materials is maintaining viable cells for long times outside of the laboratory under extreme, unpredictable conditions.

Living materials require the organization of cells within a structural scaffold. While it is possible to envision cells growing their own scaffold over macroscopic length scales, for example the growth of a tree, these processes are slow and currently difficult to control genetically. An alternative is to use additive manufacturing, also referred to as 3D printing, to organize cells within a scaffolding material at submillimeter resolution over macroscopic length scales. Additive manufacturing has revolutionized industries, from architecture to aerospace^18-23^. The range of materials that can be printed has expanded to include ceramics^24^, metals^25, 26^, nylon^27^, silk^28^, cellulose^29, 30^, and wood polymer composites^31^. The size of the objects created span from nanometers to houses^18^.

Incorporating living cells into existing 3D printingg platforms is challenging because of toxic materials and conditions, such as high temperatures to extrude plastics or UV light for curing^32-34^. Several approaches have been taken to address these issues. An object can be made using a conventional printer first, after which cells are introduced on the surface as a biofilm or diffused into a hydrogel^35, 36^. Another approach is to suspend bacteria evenly throughout a hydrogel and then use multiphoton lithography to crosslink barriers to entrap the bacteria in 3D geometries^37^. These approaches can be used to create intricate physical structures, but do not print the bacteria within the material. To this end, “bioinks” have been developed that combine bacteria and their required nutrients with the precursors that form the structural support. Zhao and co-workers developed a bioink based on *Escherichia coli* and micelles that can be UV-crosslinked after printing^38^. Using this, they constructed high-resolution 3D structures (up to 3 cm) and demonstrated that living cells can perform sensing and computing functions. Another bioink based on *E. coli* was developed that utilizes the formation of a hydrogel by alginate when it comes into contact with calcium chloride on the print surface^39^. Living materials based on *E. coli* remain functional for several days after printing. Studart and co-workers designed a bioink based on a shear-thinning hydrogel (hyaluronic acid, κ-carrageenan, and fumed silica) that was shown to be able to print *Bacillus subtilis, Pseudomonas putida* and *Acetobacter xylinum*^40^. A bioink based on *B. subtilis* has also been developed that utilizes its natural biofilm amyloids fibers to produce 2D shapes that can survive at 4°C for 5 weeks^41^. All of these approaches require that the cells be maintained in a hydrogel with ample access to water and replenished nutrients to maintain a functional living material.

Some bacteria survive under adverse conditions by forming endospores – small spherical structures that are dormant and tough. The cell membrane architecture is comprised of several layers that include an almost impermeable inner membrane, a germ cell membrane, a thick peptidoglycan cortex, the outer membrane, a basement layer, the inner coat, the outer coat and the crust^42^. The DNA is protected by its tight packing by specialized proteins^43-45^. Spores are able to survive extreme environmental insults, including high temperature, freezing, oxidizing agents, acid and alkaline solutions, genotoxic agents, solvents, high pressure, X-rays, γ-radiation, and UV-light ^46, 47^. They can also survive desiccation by replacing water with dipicolinic acid and adopting a wrinkled shape to withstand osmotic stress^44, 45^. Spores can lie dormant indefinitely and purportedly for millions of years^48^. Spore-forming bacteria have been used in biotechnology as vaccines, radiation detectors, insecticides, hygrovoltaic generators, and bio-cements^9,10,49-51^.

Here, we modify a 3D printer (MakerBot Replicator) that builds objects by extruding plastic through a high-temperature nozzle (Figure 1a). The nozzle is redesigned to mix two streams to form the bioink just prior to printing: one a polymer that is maintained at high temperature and a lower temperature mixture of cells and media. Agarose exhibits shear thinning and we show that it can be printed with various thermophilic *Bacilli* species. Using only purified spores in the bioink improves the fraction of viable cells and their distribution in the in printed structure. The spores remain distributed throughout the material and provide a constant source of germinated cells at the surface, including spontaneously in new cracks or tears. There, they can perform their programmed function, such as responding to chemicals detected by genetically-encoded chemical sensors (shown for IPTG, vanillic acid, and xylose). Further, the spore-containing material can be desiccated and stored at room temperature indefinitely. Upon rehydration, the printed shape reconstitutes, the cells germinate, and they perform their programmed function. When various extreme stresses are applied to the materials, including ethanol, high osmolarity, UV light, and radiation, the spores are able to survive and quickly germinate. This work demonstrates a route by which materials can retain the functions provided by embedded living cells long after they are created.

**Figure 1:**
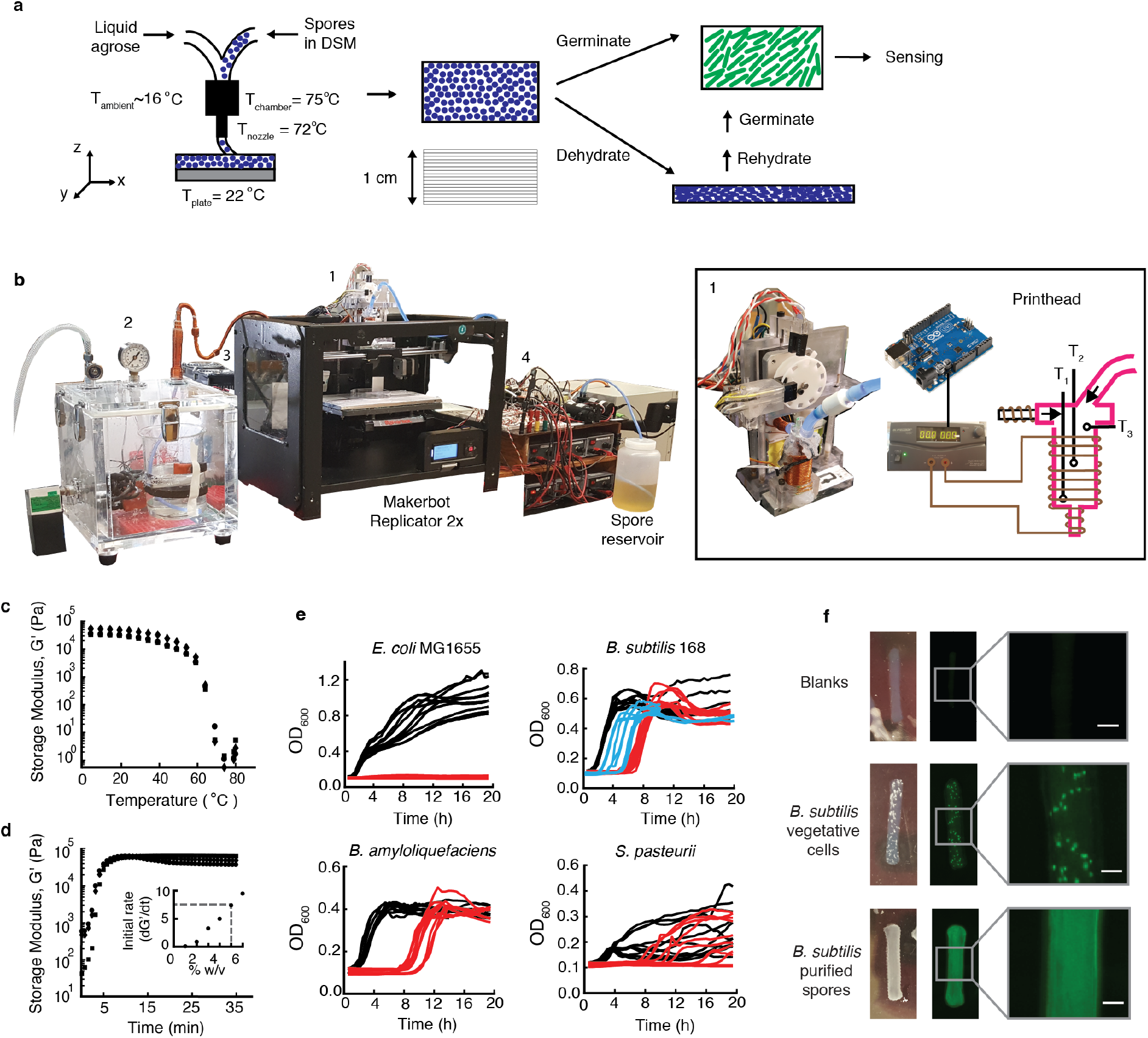
3D printer design and development of the bioink components. a, The printhead combines the material and cell streams; this schematic shows the combination of agarose and *B. subtilis* spores (blue circles). The nozzle temperature and plate temperatures are held constant and the ambient temperature of the print chamber is cooled. Once printed, the spores can be germinated (green ovals) to perform their engineered function. **b**, The modified MakerBot Replicator 2X highlighting: 1. the printhead design, 2. modified pressurized tank, 3. cooling subsystem, and 4. electronic circuit board. The locations of the temperatures measured in the printhead are shown. Details are provided in the Supplementary Information. **c**, The melting temperature of 5% agarose is shown. The points represent data from 3 independent replicates. **d**, The hardening (storage modulus) of 5% agarose when the temperature is shifted from 70°C (T_nozzle_) to 16°C (cooled print chamber). The points represent data from 3 independent replicates. The initial rate of gelation (d*G*′/dt as *t* → 0) is calculated and the inset shows its dependence on the % agarose. **e**, The response of different bacterial species to 20 minutes of heat shock at 75°C (red lines). At *t* = 0, the bacteria are resuspended in fresh media and grown at their optimal growth temperature (Methods). Each line corresponds to one growth experiment, performed in groups over three days. Blue lines show when *B. subtilis* 168 is first grown in spore-inducing media (DSM) prior to heat shock. Black lines show growth when there is no heat shock. **f**, Comparison of materials produced by printing vegetative cells or spores of *B. subtilis* 168 constitutively expressing GFP. The blank bar does not have any cells. Scale bar, 1 mm.

## Results

### Repurposing of a 3D Printer

The MakerBot Replicator was selected as a simple, inexpensive fused deposition modeling (FDM) 3D printer. The printer had to be extensively re-engineered to print living cells within a structural material (Figure 1b, Supplementary Figures 1-3). A novel printhead (#1 in Figure 1b) was designed to facilitate the blending of the structural materials and the cells from separate streams and the extrusion of the mixture through a 400 μm diameter nozzle. The streams are propelled using a modified pressurized tank for the material (#2 in Figure 1b) and a liquid DC pump for the media containing the cells (#4 in Figure 1b). The chamber where printing occurs is cooled to accelerate material hardening. Detailed photos, schematics, and electronic diagrams for these designs are provided in the Methods and Supplementary Figures 4-11.

The original MakerBot printhead has a single input that takes in a solid plastic filament and heats it to 220°C; and alternative commercial printheads were deemed inappropriate for handling cells^52, 53^. We developed a novel printhead that is able to efficiently mix the two streams while maintaining a constant temperature (*T_chamber_*) (Figure 1b, Supplementary Figure 4). The extrusion rate from the nozzle is maintained close to the rate at which the MakerBot reels in filament. When the MakerBot prints, its motor turns in the forward direction. Our printhead detects this motion with an optical sensor and it subsequently triggers the opening of solenoid valves that allows the control of both the material and cell streams. The stream is propelled by pneumatic pressure (0.75-1 psi) applied to the material storage tank that is controlled by a digital electronic relief valve (Supplementary Figures 7-8 and Table 4). This flow is discontinued when the temperature at the top of the printhead reaches ~63°C and cells have been introduced (see calibration in Supplementary Figure 9). A second optical sensor detects when the MakerBot’s motor moves via a trigger and then turns on the cell stream with a 15 ms delay (Supplementary Figures 10-11). To promote mixing in the printhead, the cell input stream enters at a 30° angle and the cell flow is set to be high (3.2 mL/s). Bubble formation during mixing (and present in the agarose) was eliminated by adding a 1 mm hole at the topmost part of the printhead for air to escape (see Supplementary Figure 4).

The MakerBot stops the filament from exiting the nozzle by turning the motor backward; in our system, this is detected by a third optical sensor, which leads to the stoppage of the material and cell streams. Early designs experienced problems with clogging (not shown). To avoid this, at the same time that a solenoid valve stops the flow of agarose near the printhead, the relief valve releases pressure in the agarose tank.

It is critical to maintain the temperature in the lines carrying agarose to and at the printhead so that it is sufficiently high to facilitate the flow of the material stream, yet is low enough to not kill the cells. For the specific material and cell combination we use (next section) this balance was achieved at 75°C (Supplementary Figures 12-13). The temperature is monitored near the nozzle output (*T*_1_), the center of the printhead (*T*_2_) and near the input streams (*T*_3_) (*T_chamber_* = <*T*_1_, *T*_2_>). *T_chamber_* is monitored using a feedback control system (a PID controller, see Supplementary Figures 14-17). *T*_3_ is used to monitor and detect when the agarose is running low and allow agarose flow via opening the solenoid valve(s). The cells are stored at *T* = 22°C and exposed to high temperature for less than 20 min. The mixture is cooled immediately after exiting the nozzle by maintaining the print chamber (#3 in Figure 1b) at 16°C using thermoelectric cooling (peltier elements) (Supplementary Figures 18 and 19). This cooling significantly improves the structural details in the printed object (Supplementary Figure 20).

To print an object, it is drawn using SolidWorks, saved in a standard format *(e.g.,* as a STL file) and loaded into the software provided by Makerbot. The gcode parameters are modified so that the nozzle temperature is 72°C, platform temperature is 22°C, and the infill is 100%. For dual printing, we merged two STL files using the merge tool in Replicator X (see picture of the system in Supplementary Figure 5). Mixing is initiated by filling the printhead with a pre-mix of material, running the temperature PID controller code to bring the temperature in the printhead to 75°C, then adding cells (Methods). After this, the objected is printed following the same protocol as the objects produced by the unmodified MakerBot.

### Material and Strain Selection

Storing the liquids containing the material and cells separately, as opposed to a single bioink, allows them to be independently varied and stored under different optimized conditions. Still, they must be compatible with each other as they are mixed in the printhead. The critical parameter in our design is temperature, where a material needs to be selected that melts at a temperature that does not kill the cells. Similarly, more thermophilic strains will be compatible with a wider range of materials that melt at higher temperatures. After testing different materials and bacterial species, we found that agarose and *B. subtilis* spores produced the most consistent structures with the highest viability of printed cells.

Agarose is a hydrogel, derived from seaweed, consisting of linear polymers of alternating D-galactose and 3,6-anhydro-L-galactopyranose subunits. It is used commonly in biology and been shown to support embedded cells^54^. Agarose was selected because it exhibits shear thinning, is transparent, has high water content, is strong after printing, and rapidly solidifies without the need for chemical/physical inputs or UV-curing. While agarose melts at 65°C (Figure 1c), we found that 72°C (T_nozzle_) is optimal for obtaining bonding between printed layers and avoid delamination.

A higher concentration of agarose leads to a more rigid structure, but is more difficult to print. We found the ideal concentration to be 4% (weight/volume). At 3%, the agarose is not able to harden into the desired geometry, even after cooling the chamber (Supplementary Figure 21). Above 6%, we started to encounter problems with line clogging. To balance these effects, the material stream was chosen to be 5% agarose so that it can be mixed with the cell stream in the printhead and remain >4% in the printed object. The storage modulus (G’) of 5% agarose is *»* 10^5^, consistent with the stiffness of previously published 3D printed hyrdrogels^17, 55^ (Figure 1d and Supplementary Figure 22). The rate at which the material hardens after exiting the nozzle is captured by the initial change in G’ upon shifting from 70°C to 16°C (Figure 1d). In our hands, a rate > 5000 Pa/s (>4% agarose) is important for accurately printing the desired structure (Figure 1d and Supplementary Figure 23). Agarose is a non-Newtonian fluid that exhibits shear thinning, which aids its extrusion through the small nozzle (Supplementary Figure 24).

Due to the temperature requirements of the modified printer *(T_chamber_* = 75°C), we needed to identify a bacterial species that can survive in the material stream. Mesophiles and thermophiles representative of different phyla were tested *(Anoxybacillus flavithermus, Bacillus amyloquifaciens, Bacillus licheniformis, Bacillus megaterium, Cupriavidus metallidurans, Escherichia coli, Geobacillus stearothermophilus, Geobacillus thermoglycosidasius, Gluconacetobacter xylinus, Gluconacetobacter hansenii, Halobacterium salinarum, Lactobacillus lactis,* and *Sporosarcina pasteurii)* (Supplementary Figures 25-27). Experiments were designed to test survival of the cells by exposing them to 75°C for 20 minutes (Methods) (Figure 1e, Supplementary Figures 26-27). Of the species tested, only spore-forming organisms were able to survive with little day-to-day variability. When *B. subtilis* PY79 (a derivative of 168)^56^ is first grown in spore-inducing media (DSM), this enhances survival and allows for cells to more rapidly recover heat shock (Figure 1e blue lines). Furthermore, we exposed the spores to higher temperature for 20 min (>75°C and up to 100°C) and found that spores can survive at 100°C, but growth is not detected until after 8h (Supplementary Figure 28).

The 5% agarose and *B. subtilis* PY79 in DSM were selected as the two components of the bioink. Note that other spore-forming organisms would also work as the cell component of the ink (Supplementary Figure 27). A simple bar was printed consisting of 7 layers (2 by 3 by 25 mm) (Figure 1f). To aid imaging, an expression cassette where green fluorescent protein (GFP) is constitutively expressed was introduced into the *B. subtilis* genome at the *amyE* locus (Methods). Figure 1f shows comparisons between the structures produced when *B. subtilis* is grown under conditions favoring vegetative cells versus spores. The former shows punctate growth distributed unevenly through the agarose structure. When spores are printed, they are evenly distributed throughout the structure.

### 3D Printing Functional Living Materials

Figure 2 shows examples of geometries that can be printed by the material stream alone and together with the cell stream. The minimum feature size is consistent with that obtained by the MakerBot Replicator when printing plastics (100 μm), which is dictated by the 400 μm nozzle diameter. The hydrogel can be printed at a scale the utilizes the maximum size in the x-and y-dimensions (10 cm by 20 cm). Simple shapes can be printed up to 1 cm in the z-dimension, requiring the printing of ~37 layers, each 300 μm (at a rate of 3 layers/min). In Figure 2i, we compare the printed objects with those obtained from a high-resolution commercially-available printer (ProJet 6000) and the shape and resolution is similar.

**Figure 2:**
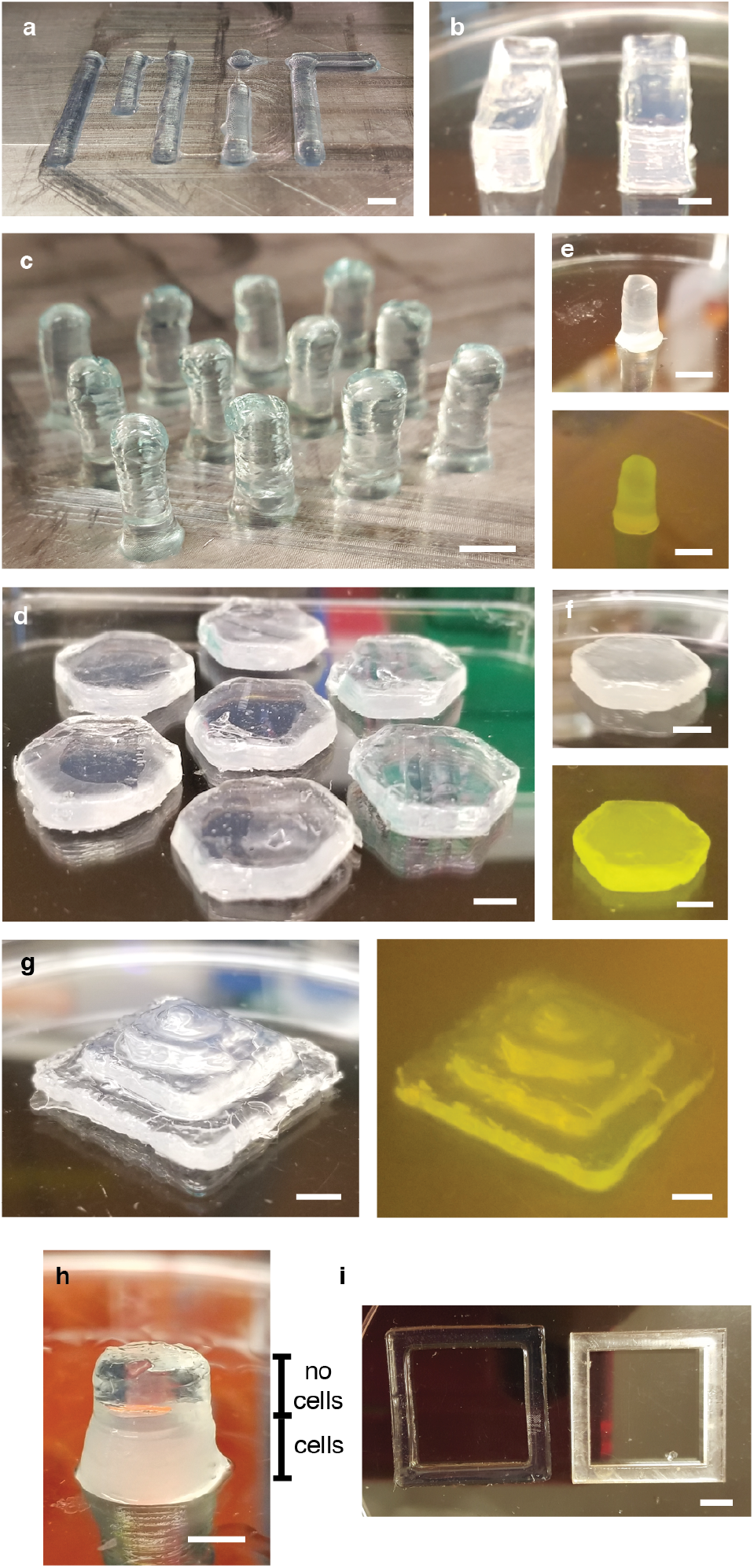
3D printing of living materials. a-d, Examples of printed objects are shown when the 5% agarose material stream is printed alone. **(e,f,g)** Objects are shown where spores of GFP-expressing *B. subtilis* 168 are printed with the agarose. GFP is imaged by illuminating the object with blue light from a Safe Imager (Invitrogen, Carlsbad, CA). **h**, An object where the bottom half is printed with cells and the top half without using two printheads (Methods). **i**, Comparison of printed parts with that of a commercially available printer (ProJet 6000). Scale bars, 5mm.

There are some applications where it is desirable to print cells at specific locations, rather than throughout the structure. To do this, a second printhead is added into which the material stream is injected via a tee connector, but for which there is no cell stream input (Supplementary Figure 5). The STL files are prepared for the portions containing and lacking cells and then merged. This generates structures with sharp boundaries between the regions containing cells (Figure 2h). It is straightforward to expand this approach to print multiple materials and cell types.

The spores are distributed evenly throughout the material in which they are printed. They germinate preferentially at the media-exposed surface of the hydrogel due to the availability of oxygen and nutrients (Figure 3 a-c and Supplementary Figure 29). Germination occurs up to a depth of 0.85 ± 0.16 mm, determined by imaging cells constitutively expressing GFP. When the structure is torn and incubated in LB media for 10 hours, this induces the germination of spores at the surface exposed to the media (Figure 3c). This demonstrates that the embedding of spores in the material provides, under the right conditions, a source of new germinated cells that can perform their function.

**Figure 3:**
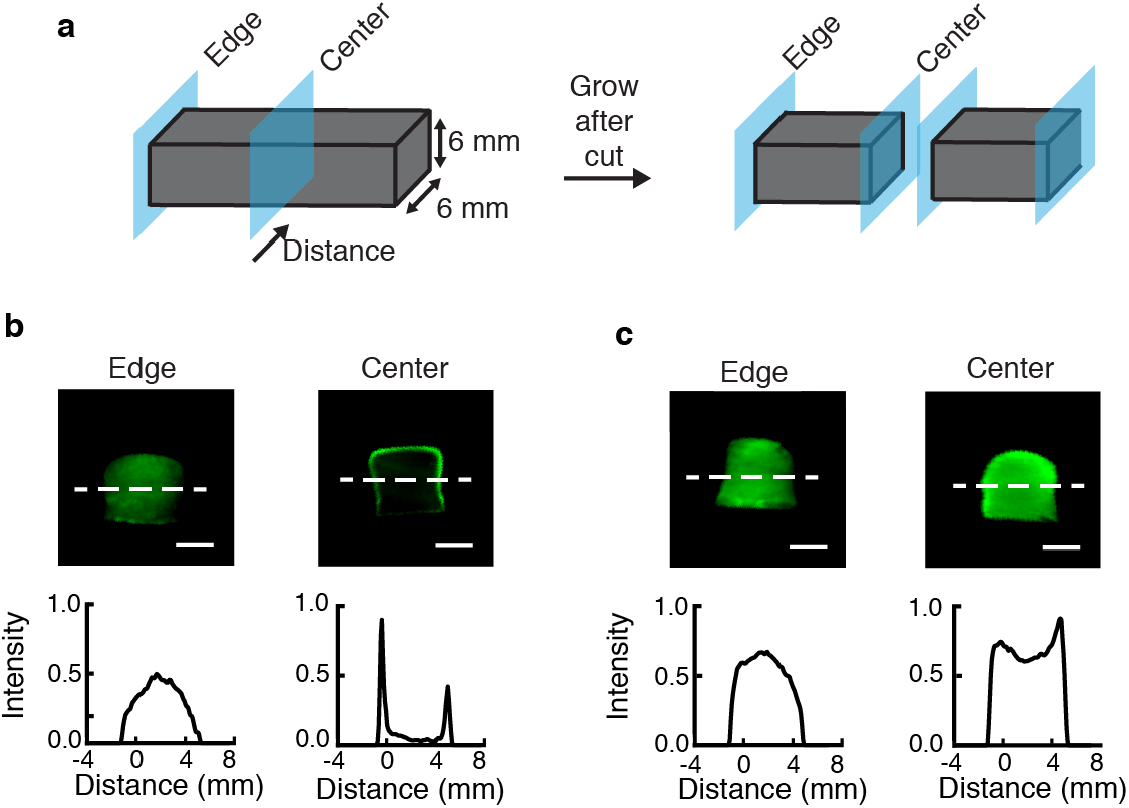
Cutting the object induces germination of spores. a, Objects were printed with spores of *B. subtilis* PY79 constitutively expressing GFP. **b**, When imaged immediately after cutting, germinated cells are only present at the edge (Methods). **c**, After incubation for 10 hours in LB media, germinated cells are present on the new surface. Images representative of three experiments performed on different days are shown. The gray scale value obtained is normalized by dividing the values by 255. Scale bars, 3mm.

The germinated cells can function as biosensors. To demonstrate this, we implemented three genetically-encoded sensors in the genome of *B. subtilis* that respond to small molecules. The sensors were either taken from the literature or built *de novo* and optimized for *B. subtilis* to maximize their dynamic range, such that they can be imaged in the material. An isopropyl B-D-1-thiogalactopyranoside (IPTG)-inducible system was used that had been previously reported to have a dynamic range of 276-fold^57, 58^ (Figure 4a and Supplementary Figure 30). A sugar sensor (xylose) was modified to function in LB media by removing the catabolite-responsive element (Figure 4c, Supplementary Figure 31, Supplementary Table 7)^59^. Finally, a sensor for a plant root exudate (vanillic acid) was designed based on an optimized VanR repressor and corresponding operator that was inserted into the -10/-35 region of P_spank(hy)_ to create P_spank(V)_ (Figure 4d, Supplementary Table 7)^7, 60^. This sensor generates a 60-fold dynamic range (Supplementary Figure 32).

**Figure 4:**
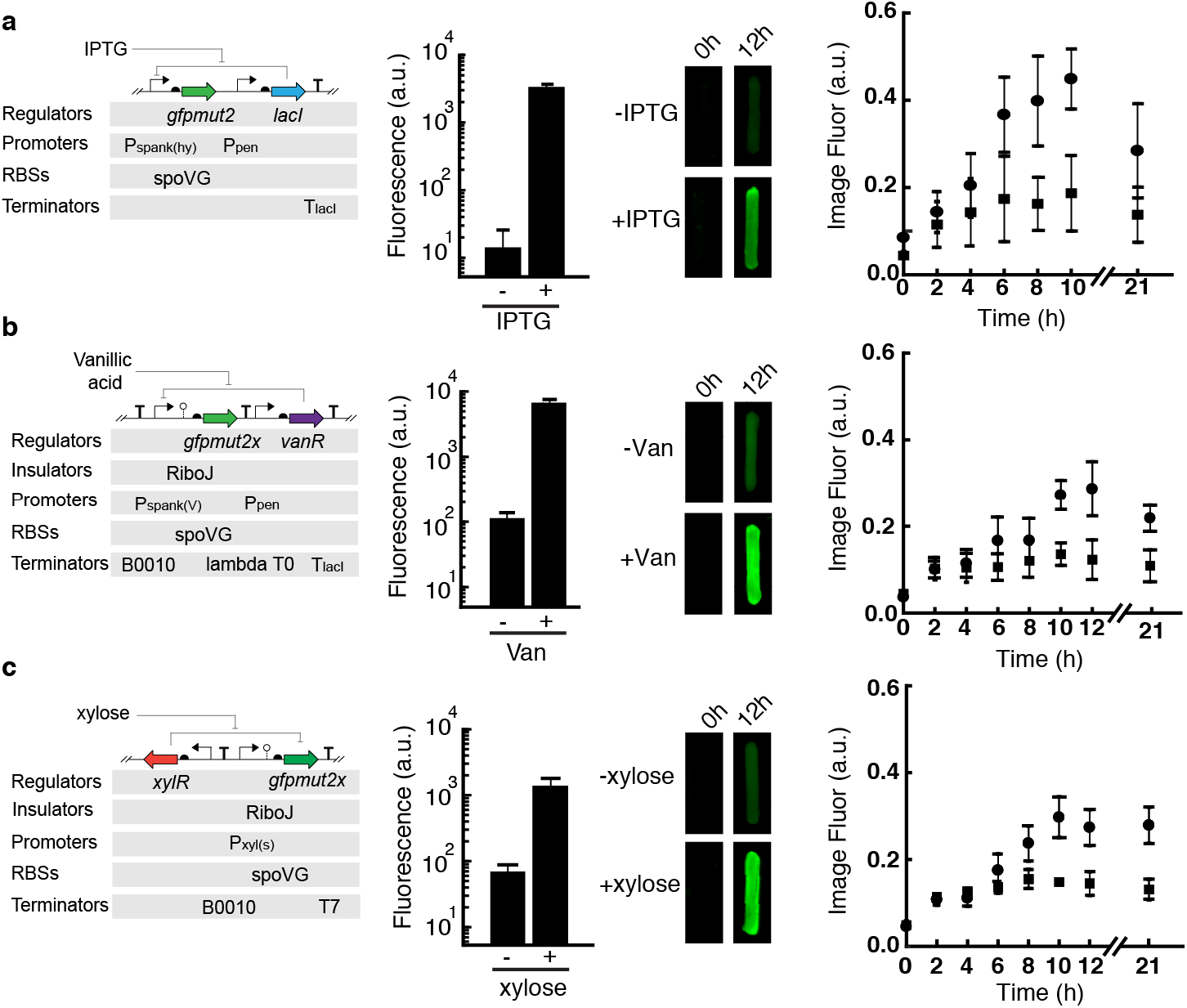
Induction of genome-encoded sensors in the printed material. Sensors were designed for *B. subtilis* that respond to: (a) IPTG, (b) Vanillic Acid and (c) Xylose (Supplementary Figures 30-32). The bar graphs (left) show the induction of cells (1 mM IPTG, 1 mM Van and 1% Xylose, respectively) for 90 minutes in LB media, measured by flow cytometry (Methods). Bars were printed (2 by 3 by 25 mm) with spores of *B. subtilis* containing the sensors. The bar fluorescence was imaged before and after induction for 12 hours in LB. The timecourses quantify the fluorescences of the bars (right) as quantified from the Images (Methods). The means and standard deviations of three experiments are shown, performed on different days.

Materials were printed with spores of bacteria containing the sensors (Figure 4). After printing, the material was incubated at 37°C in media either containing or lacking the small molecule. For all the sensors, some induction can be seen at 6 hours with full induction at 12 hours.

### Spore Survival in 3D Printed Materials

The sensitivity of living cells and their continuous requirement for nutrients and water, make it challenging to embed them within materials and function over long times. First, we evaluated the ability for the printed materials to survive desiccation. The 5% agarose material contains 92% water (Methods). After it is printed, the water can be removed by drying and stored for long times (Figure 5a). When rehydrated, it returns to its original printed shape (Figure 5b). When spores of GFP-expressing *B. subtilis* are printed, they survive for 1 month (the maximum tested) and germinate when rehydrated. GFP expression after germination and growth does not go down for longer storage times (Figure 5c, Supplementary Figure 33). Given the known longevity of spores^61^, it is expected that they would be able to regerminate after indefinite storage under these conditions.

**Figure 5:**
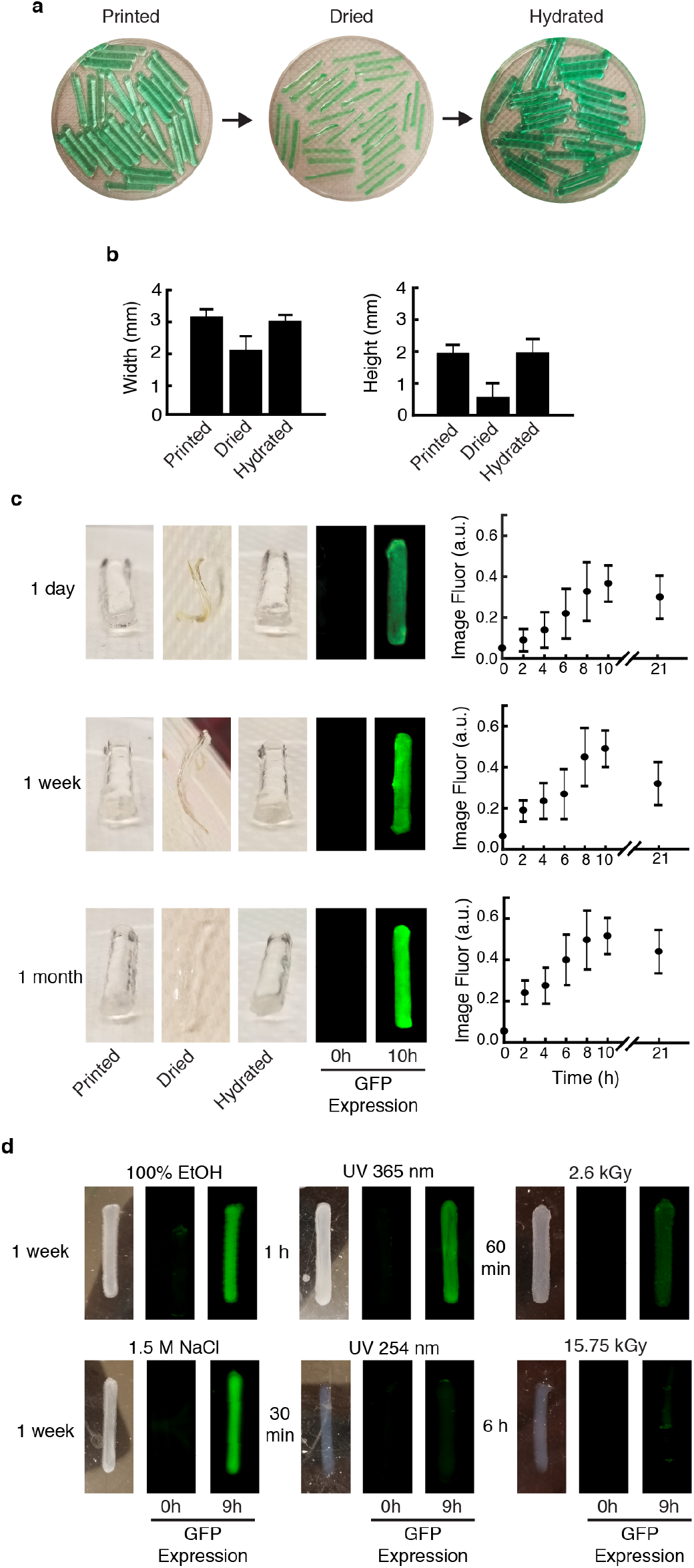
Resilience of living materials printed with *B. subtilis* spores. a, Reconstitution of the shape of the printed (2 × 3 × 25 mm) bars that have been dried at room temperature then hydrated with water (Methods). **b**, Quantification of the width of the bars. **c**, After dehydration, the bars were stored at ambient temperature on a lab bench for the indicated times and then rehydrated. The spores in the bars were germinated by placing them in LB media. **d**, Bar exposed to other challenging conditions such as 100% ethanol, high osmolarity (1.5 M NaCl), UV light (365 and 254 nm) and g-radiation (2.6 and 15.75 kGy). Images of the expression of GFP from a constitutive promoter are shown (Methods). The means and standard deviations of three experiments are shown, performed on different days. Additional time points and controls are shown in Supplementary Figure 34-37.

Spores can survive under more extreme environmental stresses. Alcohol is known to not have sporicidal activity. When printed objects containing the spores are exposed to 100% ethanol for one week, the cells survive (Figure 5d and Supplementary Figure 34). High osmolarity has been shown to have an effect on spore viability^62^. Again, after one week, the spores remain viable. Note that, for both of the stresses, some of this protection may be being conferred by the agarose material.

The materials containing spores were then subjected to UV light, X-rays, and γ-radiation (Figure 5d and Supplementary Figures 35-37). The spores survive 365 nm UV light treatment, but they do not survive when exposed to 254 nm light. This is expected as 254 nm UV light is known to be effective at exterminating spores^63^. The spores also survive when the material is exposed to X-rays for 10 min (8.469 R/min). When exposed to γ-radiation, the spores survive after 1 hour (2.6 kGy) exposure, but not 6 hour (15.7 kGy) or 12 hour (31.5 kGy) treatments. Note that γ-radiation with a dose of 25 kGy is used for sterilization^64^. In addition, after 6 hours the agarose hydrogel weakens and the bars start to deteriorate due to shaking during the growth assay (Methods).

## Conclusion

This work demonstrates that embedding spores, as opposed to vegetative cells, offers advantages when producing living materials. Using spores simplifies the preparation and storage of the cell component of the bioink and generates more reliable prints without having to carefully control the viability or growth phase of the cells in the reservoir. While we print the spores within an agarose hydrogel, they are able to survive extreme desiccation and storage under ambient conditions without a continuous nutrient supply. When they germinate, they are able to perform their programmed function, whether it be to sense a chemical or turn on a biosynthetic pathway. Further, the spores selectively germinate on the object surface. This allows the spores in the center of the material to provide a continuous source of freshly germinated cells. When the material is torn, the spores germinate and grow on the new surface. One could imagine harnessing this capability, along with the ability to produce biopolymers^65^, to create self-healing materials. While we utilize the natural signals that induce germination, methods have been developed to genetically engineer spores to germinate under defined conditions, such as the presence of chemicals, including neurotransmitters and specific DNA sequences^66^. The recalcitrant nature of spores makes this approach potentially compatible with a wide range of 3D printing technology based on harsher conditions, such as laser sintering of ceramics/metals (high T).

Many applications of synthetic biology require that engineered cells survive and perform their function in fluctuating and stressful environmental conditions^67^. UV protection will be required for bacteria that function in the open environment, for example on plant surfaces in agriculture,^68^ in open ponds for bio-production^69^, or pollution remediation^70^. The ability of spores to survive without water and nutrients and under harsh conditions can aid long term storage. Harnessing the recalcitrant nature of spores, stabilized within a 3D printed hydrogel is a step towards realizing these applications.

## Methods

### Modifications to the 3D Printer

We made extensive modifications to a MakerBot Replicator 2X, entailing five separate subsystems: a printhead, an agarose pumping subsystem, a cell pumping subsystem, and a heating subsystem and a cooling subsystem (Supplementary Figure 1). The cells and agarose are propelled to the printhead using the agarose pumping and the cell input subsystems. At the printhead, the temperature is regulated using the heating subsystem. At the other extreme, the cooling subsystem maintains the chamber at the proper temperature for the agarose to retain its filamentary shape when it exits the nozzle. Details regarding these modifications, including photos, schematics, part numbers, files for (traditional) 3D printing of novel parts, and Arduino programming diagrams are provided in the Supplemental Information.

#### Printhead and optical sensors subsystem

The original MakerBot printhead has a stepper motor that extrudes the plastic filament, drawn in as a solid filament from a single inlet. Our printhead’s barrel has two inlets, one for the material and the other for the cells. It also has three thermistor sensors to measure the temperature at the top, middle and bottom of the printhead (*T*_1_, *T*_2_, *T*_3_) (Supplementary Fig. 4a and Supplementary Figure 4b). The average of *T*_1_ and *T*_2_ is used to maintain the temperature in the printhead at 75°C to ensure mixing of the agarose and spores (see cell input and heating subsystem sections). A dual printing configuration is used by alternating between two solenoid valves to have two different printhead printing either the material alone or with the cells (Supplementary Figure 4c). Optical sensors are used to detect which motor moves and thus, which extruder is active at a given time. Using a pair of optical interrupters, we coupled the stepper motor (from the MakerBot) and the agarose pumping system. An encoder (10 spokes) monitors the stepper motor’s movement. When a movement change is detected and the temperature is ~63°C (at the very top of the printhead), the solenoid valve is open so that agarose flow freely into the printhead. A slot optical interrupter is a U-shaped sensor with an infrared emitter on one side and the corresponding detector on the opposite side. When an object breaks the infrared beam, this can be detected through a break in the circuit. Two 8 mm wide slot sensors were placed in tandem (Digi-Key Electronics part no. EE-SX1070). These are also used to detect when the motor reverse direction using a quadrature encoder signal. Briefly, if the two channels are 90° out of phase and if pin 2 sees a change before pin 3, then the motor is moving clockwise; otherwise, it is moving counterclockwise (see code). A single optical interrupter (Digi-Key Electronics part no. EE-SX1042) was used to turn on the cell stream for during the time it takes for two revolutions to occur (see cell input subsystem for more details). A hole (1 mm in diameter) was introduced at the top of the printhead for excess air to escape after priming and while printing (air from the agarose cabinet is constantly purged into the printhead). This modification relieved the problem with having bubbles inside the printed object that distorted the material and led to non-uniform prints. In Supplementary Figure 6, we show a schematic of the electronic circuit used to control the three slot sensors. Each of the two printheads contains three optical sensors.

#### Agarose pump subsystem

A hermetically sealed desiccator cabinet (Fisher Scientific, Model # 33060) was repurposed as a pneumatic pump (Supplementary Figure 7). This cabinet has a ¾” thick wall made of polymethylmethacrylate (PMMA). The vacuum gauge was replaced with a pressure gauge (McMaster-Carr, part no. 4000K721) with a range of 0-15 psi to monitor and maintain a constant pressure in the cabinet. One orifice in the cabinet was used as an inlet for house air to enter and the other orifice was used as an outlet for agarose to leave using positive pressure (Supplementary Figure 7a and b). To maintain agarose flow, a magnet wire (Applied Magnets, AWG 21) was wound around the plastic tubing such that a passing current releases heat (Joule heating). Tubing consisted of high-temperature silicone rubber. A thicker tubing (OD = 3/4”, ID = ½”, McMaster-Carr, part no. 5236K47) was used to fit the agarose outlet and a straight reducer barbed fitting (McMaster-Carr, part no. 5463K635) to connect to a thinner tubing (OD = 5/16”, ID = 3/16”, McMasterr-Carr, part no. 5236K841) that fits the solenoid valves. The length of the tube is 27” long to provide some slack for the printhead to move. The current in the power supply was set to 5.0 A and the voltage is ~ 12V (BK Precision 1901B Switching Mode Power Supply). In addition, the cabinet has an electrical outlet used to plug a heating tape for maintaining the agarose in the reservoir as a liquid. The tubing that carries agarose from the reservoir to the agarose outlet on the lid does not have any wire windings and its tubing is smaller (OD=1/4”, ID=1/8”, McMasterr-Carr, part no. 5236K83). There is an o-ring (size 114) in the inlet and outlet of the lid of the cabinet to prevent leakage. In addition, a check valve (McMaster-Carr, part no. 6079T58) prevents the flow of this viscous liquid back into the reservoir. An arrow in relief on the check valve was sanded down and the two layers of the magnet wire was wound around this valve. A 1 pole push-in signal power connector (McMaster-Carr, part no. 9193T11) was installed for maintenance as sometimes this valve gets clogged and needs to be cleaned. Epoxy was used to permanently fix the magnet wire to the check valves. This check valve adds about 1 psi of resistance to the line; more check valves led to severe clogging and cleaning difficulty. An electronic relief valve was installed in the cabinet to immediately (within a few ms) relieve the pressure when it is not needed to actuate flow to the printhead. When starting a print, the line is initially primed with molten agarose, which keeps this relief valve closed. When the mixing chamber’s temperature reaches ~63°C, this indicates it is full, and the valve relieves the pressure in the tank (Supplementary Table 4). When the molten agarose is pumbed from the cabinet, the pressure is increased and pulsed on and off so that it remains between 0.75 and 1 psi. A pair of solenoids is placed between the check valve and the printhead (Supplementary Figure 7a and Supplementary Figure 5) to exert more control over the agarose flow and to facilitate dual printing (see section on Printhead and slot sensors subsystem for more details). Supplementary Figure 8 shows a schematic of the electronic circuit used to control the pumping of the agarose.

#### Cell input subsystem

A DC liquid pump (Trossen Robotics, part no. HW-WVALVE) (Supplementary Figure 9a) is used to propel the cell stream into the printhead. The flow is modulated by a solenoid valve that is controlled by an Arduino microcontroller. To reduce the pump flow rate, 8 check valves are placed in tandem, including one close to the entry point to the printhead (each contributing ~1 psi of resistance) (Supplementary Figure 9b). We calibrated the pump with the time that the pump is energized or open and determine the amount of liquid to add for continuous pumping and to keep the agarose percentage above 4% (Supplementary Figure 9c). After the initial agarose priming of the printhead, it contains about 5 mL of agarose and we input 600 μL of cells to keep the agarose percentage above 4%. To add 600 μL of cells, we opened the DC liquid pump (and solenoid valve) for 90 ms (see DC liquid pump calibration in Supplementary Figure 9c). For continuous and subsequent input of the cells, we did a second set of calibrations to determine how much cells to add per 100 μL of agarose and chose 15 ms for the time the DC liquid pump and solenoid valves are opened (Supplementary Figure 11).

#### Heating subsystem

One power supply (Newark, part no. 78Y9404) is kept at a constant voltage/current and is used to power the heating of the materials line to keep the agarose molten prior to reaching the printhead. To mix cell and agarose streams, the temperature in the printhead needs to be maintained 75 °C (Supplementary Figure 13). To do this, a second power supply (Newark, part no. 34T4663) is controlled remotely and we implement a PID controller to maintain this constant temperature in the printhead. The PID algorithm (Supplementary Figure 14a) compares the setpoint and measured temperatures and used their difference (the error) to control the current from the power supply based on the equation

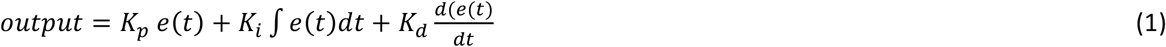

where the output is the signal controlling the current, t is the time, *K_p_, K_i_* and *K_d_* are the coefficients for the proportional, the integral and the derivative, respectively. It is implemented using an Arduino PID library and the tuning parameters are adjusted using an autotune function (SetTuning). The feedback control is implemented using the remote-controlled power supply with an 8 pin plug adapter (only pin 1 (5V), pin 2 (input) and pin3 (ground) were used), magnet wire, thermistors and an Arduino microcontroller (Supplementary Figure 14b). Supplementary Figure 15 shows a schematic of the circuit wiring. We verified that the system and code work at room temperature (Supplementary Figure 16) and while cooling the chamber (Supplementary Figure 17). The temperature of the nozzle is independent of the printhead and we controlled it through the gcode in the original MakerBot. We screened for the best temperature to keep the nozzle and we found it to be 72°C (T_nozzle_ + T_chamber_ + T_ambient_) which leads to an average temperature of ~54°C.

#### Cooling subsystem

The rapid gelation of the agarose/cell mixture after leaving the printhead nozzle required the cooling of the print chamber to 14-17°C by blowing cold air into it. A Styrofoam box with an aluminum pan was placed behind the printer (Supplementary Figure 18). Two metal bars that span across the pan were attached to peltier plates (Supplementary Figure 19). A fan blows air into the cooling system and two aluminum elbow pipes move the cold air into the printing chamber. Metal fins were attached to the bars to increase the surface area and facilitate heat transfer as air passes through these fins. Prior to using the printer, 1L of water is added to the aluminum pan, it is cooled for one hour, and then ice is added to the pan.

#### Preparation and storage of agarose

The SeaPlaque agarose (Lonza, cat no. 50100) powder is dissolved in 300 mL of water (to make a 5% w/v), autoclave and stored at room temperature. Right before adding it to the material reservoirs (a 600 mL beaker with a heating tape wrapped around it) the agarose is microwaved for 3.5 min or until it completely melts. We cover the beaker with a piece of aluminum to prevent excessive evaporation inside the desiccator cabinet.

#### Embedding and scaffolding materials characterization

Using a rheometer (AR2000, TA Instruments), we characterized the gelation time, the melting temperature, and the shear thinning properties of the agarose. All of the measurements were taken using a peltier plate (with elements for rapid cooling and heating) (it can rapidly cool and heat). To obtain adequate rheometer measurements of the hydrogels, a cross-hatched geometry (543337.001) was used. This provides a rough surface to grip on the hydrogels. To avoid slipping of the hydrogel on the metal surface of the peltier plate and provide some texture, a self-adhesive sand paper (320 grit, cat. McMaster-Carr, no. 4647A13) was used (Supplementary Figure 22a). To prevent drying of the hydrogels, we used an evaporation solvent trap (Supplementary Figure 22; inset). The % strain was set to 0.1 and the frequency at 1Hz for all measurements. The samples were prepared as in the previous section, but in a smaller volume (5 mL). To run the samples on the rheometer, about 1 mL is pipetted onto the self-adhesive sand paper, the geometry is lowered to about 1050 μm from the surface and let solidify. The excess material is then removed using a spatula so that cross-sectional area matches that of the geometry (490 mm^2^) and this geometry is then lower to 1000 μm (or specified gap in the file).

### Molecular Biology and Genetic Engineering

#### Strains and culture conditions

For a complete list of strains used in this study see Supplementary Table 5. Chemically-competent *Escherichia coli* DH10B cells were used for all routine cloning (New England Biolabs, Ipwich, MA). The *E. coli* MG1655 (in Figure 1e) and the various Bacillus strain were obtained from the lab stock originally obtained from either ATCC or Bacillus Genetic Stock Center (BGSC). All *E. coli* and *Bacillus* strains were grown at 37°C in LB-Miller media (BD, cat no. 244620) unless indicated. *Anoxybacillus flavithermus* strain y.d. (DSM 2641) cells were grown at 55°C in Nutrient Broth (5g/L of peptone (BD, cat no. 211677), 3g/L of beef extract (BD, cat no. 212303), pH adjusted to 7.0) and Anoxybacillus flavithermus strain WK1 (DSMZ, Germany) cells were grown at 55°C in Caso Broth (15g/L of peptone from casein (BD, cat no. 211921), 5g/L of peptone from soybean (Sigma-Aldrich 70178-100G), 5 g/mL of NaCl). *Geobacillus thermoglucosidasius* M5EXG (ATCC BAA-1069) were grown at 55°C in Tryptic Soy Broth (40g/L, BD cat. 211825). *Lactococcus lactis* subsp. cremoris (Mobitech, MG1363), *Lactococcus lactis* subsp. lactis (ATCC 11454) were grown in Brain Heart Infusion (BHI) Broth (BD, 237500). *Gluconacetobacter xylinus* (ATCC 700178) *and Gluconacetobacter hasenii* (ATCC 23769) were grown in Hestrin and Shramm (HS) medium at 30°C. HS contains 2% (w/v) of glucose (Fisher Scientific, C6H1206) 0.5% (w/v) of yeast extract (BD, cat no. 210929), 0.5% (w/v) peptone (BD, cat no. 211677), 0.27% (w/v) of sodium phosphase diabasic (Fisher Scientific, S375-500), 0.15% (w/v) of citric acid (Fisher Scientific, A940-500) and 0.1% (w/v) of cellulase (Sigma-Aldrich, cat no. C2730-50ML). The glucose and the cellulase were filter sterilized using a 0.2μm Acodisc syringe filter (Pall, cat no. PN 4612). *Cupriavidus metallidurans* CH34 (ATCC 43123) and a domesticated strain derived from *Sporosarcina pasteurii* (ATCC 11859) were grow in Nutrient Broth (BD cat. 234000) at 30°C. To grow the *S. pasteurii,* the pH of the Nutrient Broth was adjusted to 8.5 before autoclaving. *Halobacterium salinarum* NRC-1 (ATCC 700922) were grown in 2185 Halobacterium NRC-1 medium at 30°C. The 2185 Halobacterium NRC-1 medium is made with a Basal medium and trace metals, as described in the ATCC protocol. Minimal S750 medium contains the following ingredients in a total volume of 100 mL: 10 mL of 10X S750 salts, 1 mL of 100X metals, 2 mL of 50% (w/v) arabinose (Sigma-Aldrich, cat no. A3256-100G), 1 mL 10% glutamate (Fisher Scientific, cat no. A125-100). The 10X S750 salts consist of (filter sterilized): 0.5 M MOPS (free acid) (Fisher Scientific, cat no. ICN19483725), 100 mM of ammonium sulfate (NH_4_)_2_SO_4_ (Millipore Sigma, cat no. AX1385-1), 50 mM of potassium phosphate monobasic (KH_2_PO_4_) (Sigma Aldrich, cat no. 795488-500G), 8g of potassium hydroxide (KOH) (Sigma-Aldrich, cat no. 221473-500G) (finely adjusted to pH 7 with 1M KOH). The 100X metals consist the following metals (filter sterilized): 0.2 M of magnesium chloride (MgCl_2_) (Sigma-Aldrich, cat no. M8266-100g), 70 mM of calcium chloride (CaCl_2_) (Sigma-Aldrich, cat no. C1016-500G), 5 mM of manganese (II) chloride (MnCl_2_) (Sigma-Aldrich, cat no. 244589), 0.1 mM of zinc chloride (ZnCl_2_) (Sigma-Aldrich, cat no. Z0173), 100 mg mL^-1^ of thiamine-HCL (Sigma-Aldrich, cat no. T4625-25G), 2 mM of HCL (Millipore Sigma, cat no. HX0603-3), and 0.5 mM of iron (II) chloride (FeCl_3_) (Sigma-Aldrich, cat no. 701122-1G). For E. coli, the following antibiotic concentrations are used (both plates and in culture): 100 μg/ml ampicillin (Ap, GoldBio; CAS#69-52-3); 50 μg/ml kanamycin (Kn, GoldBio; CAS#25389-94-0), 100 μg/ml spectinomycin (Sp, GoldBio; CAS#22189-32-8) and 35 μg/ml chloramphenicol (Cm, AlfaAesar; #25-75-7). For *B. subtilis,* the following concentration were used: 5 μg/ml Cm, 100 μg/ml Sp, 5 μg/ml kn.

#### Plasmids and strain construction

All plasmids were constructed using Type IIS assembly. DNA sequences were inspected and modified to eliminate the recognition sites for the restriction enzymes (BsaI or BbsI or BsmbI) as necessary. A bsaI site was removed from the original gfpmut2 and it was named gfpmut2x. DNA was amplified using the polymerase Kapa Hi Fi DNA polymerase (Kapa Biosystems, cat no. KK2602) and gel purified using Seakem GTG agarose (for nucleic acid recovery) (Lonza, cat no. 50070), agarose dissolving buffer (ABD) (Zymo Research, cat no. D4001-1-50), Zymo-Spin I (Zymo Research, cat no. C1003-50) and collection tubes (Zymo Research, cat no. C1001-50). All genetically-modified *B. subtilis* strains are based on *Bacillus subtilis* PY79 (obtained from the lab stock). All DNA designed for *B. subtilis* were integrated into the genome at the amyE site. Genomic integration was verified by polymerase chain reaction (PCR).

#### Preparation of B. subtilis spores

To induce *B. subtilis* sporulation, we used the Difco Sporulation Medium (DSM) containing 8 g/L of Difco Nutrient Broth (BD, cat no. 234000), 1g/L of potassium chloride (Sigma-Aldrich, cat no. P5405-500g), 0.25 g/L of magnesium sulfate heptahydrate (MgSO_4_ ·7H_2_0) (Sigma-Aldrich, cat no. 230391-500). The following solutions were added to DSM after autoclaving: 1M calcium nitrate hydrate (Ca(NO_3_)_2_) (Sigma-Aldrich, cat no. 202967-10G), 0.01 M MnCl_2_(Sigma-Aldrich, cat no. 244589-500G) and 1mM iron (II) sulfate (FeSO_4_) (Sigma-Aldrich, 215422-5G) (to 1L, 1mL of each). *B. subtilis* strains were streaked on LB plates (with corresponding antibiotics) and single colonies were grown in two culture tubes (in 3 mL of DSM) for 3h prior to transferring this inoculum to 500 mL in 2800-L Erlenmeyer flask. These cells were grown for 2 days, shaking at 250 rpm, at 37°C, and in a New Brunswick Scientific Innova 44 incubator. The OD_600_ was measured to be ~1. We verified the presence of spores in the culture by microscopy. The cells were store at 4°C prior to use in the printer. This media was transferred to a 1L Nalgene centrifuge bottles (Fisher Scientific, 05562-25) for loading into the printer. The lid of these bottles were modified by drilling a hole (0.25” in diameter) to fit a tubing (McMaster-Carr, cat no 5236K83). The other end of the tubing was tightly fit into the inlet of the DC liquid pump.

#### Heat shock and outgrowth experiments

A thermo cycler (Bio-Rad C1000 Touch) was use to heat shock the cells for 20 min at 75°C. Cells were streaked in plates then single colonies were picked and grown in their strain-specific media and temperature prior to dilution. The cells were heat shocked in 50 μL (OD_600_ = 0.04) and transferred to plates (Nunc 96-Well Optical-Bottom Plate) containing 150 μL of their respective medium (final of OD_600_ = 0.01). The cells were grown in a multi-mode microplate reader (Biotek, Synergy H1) for 20 hours with continuous orbital shaking and at the respective strain temperature.

#### Measurement of GFP in 3D Printed Parts

The printed objects were grown in 5 mL of LB and incubated with shaking (250 rpm) at 37°C. The parts were washed with 12 mL of 1X phosphate-buffered saline (PBS) to reduce the auto fluorescence of the LB media. The excess PBS was gently removed using Kimwipes (Kimberly-Clark™ Professional 34120). The objects were then imaged with a ChemiDoc MP Imaging System (BioRad, Hercules, CA) using the Alexa 488 filter and a 0.5 sec of exposure time. The plasmid pLG166 was used to make the +GFP *B. subtilis* strain LMG09 (Supplementary Figure 33a). The *gfpmut2x* gene is driven constitutively with the promoter *P_pen_* and a spectinomycin cassette was used for selection. The plasmid pLG173 was used to make the -GFP *B. subtilis* strain LMG16 (Supplementary Figure 33b). To quantify and to do the image analysis of the data shown in Figure 4 and 5, we loaded the raw images into a routine macro for ImageJ (available upon request). The reported mean fluorescence values are the +Gfp cells.

#### Drying and rehydration experiments

The bare agarose materials were printed and dried overnight at room temperature. A green dye was used to color the agarose (Figure 5a). The dried object was hydrated with deionize autoclaved water. The width and the height were measured using a digital caliper (Global, item# WG534250). For the longevity experiments, we dried the materials overnight in petri dishes, then wrapped them with 2” parafilm (VWR, cat no. 52858) and stored at room temperature for 1 day, 1 week and 1 month. For the hydration part of the experiment, we completely submerged the 3D printed parts in 1X phosphate-buffered saline (PBS) for 2 hours prior to transferring the 3D objects to the routine culturing conditions as denoted in the previous section *(Measurement of GFP in 3D Printed Parts).* To calculate the water content (72%), the absolute value of the difference in the wet weight and the dry weight (after drying for 24 hours at room temperature) of the hydrogel is divided by the wet weight multiplied by 100%.

#### Inducible systems characterization in culture

Chemical inducers were used in the following concentrations unless indicated: 1mM of isopropyl-β-D-thiogalactoside (IPTG) (Gold Bio; CAS 67-93-1), 1mM of vanillic acid (Sigma CAT 94770) and 1% (w/v) xylose. Prior to using the instrument, single colonies were inoculated in 2 mL of LB (with the respective antibiotic) and grown at 37°C with 250 rpm shaking for approximately 3h. The cells were back diluted to OD_600_ of 0.001 and grown for 1 hour (or until early log phase) in plates (Costar Assay 96-well plate clear round bottom) using an Elmi shaker at 1000 rpm at 37°C. After this time elapsed the corresponding inducers were added and then the cells were allowed to grow for 1.5 h.

#### Inducible systems characterization in material

The same culture conditions as described in the Measurement of GFP in 3D Printed Parts section was used. The only difference is that the inducers were added at t = 0 hr. The concentration of inducers added for each sensor was the same as in culture (see Inducible systems characterization in culture). The strains used in this set of experiments were the *B. subtilis* LMG04 (IPTG inducible system), *B. subtilis* LMG117 (the vanillic acid inducible system), and *B. subtilis* LMG125 (the xylose inducible system).

#### Spores in the core of the materials

Blocks with dimension of 6 × 6 × 25 mm were printed using the spore of the *B. subtilis* strainLMG09 (gfp constitutively active) and the *B. subtilis* strain LMG16. The cells were grown as indicated in the section *Measurement of GFP in 3D Printed Parts,* except, that the media was replaced every 3h and the cells were incubated for a total of 10 hours. The blocks were cut in half using disposable scalpel (Electron Microscopy Science, cat no. 72042-21) prior to incubation. The control group were the intact blocks (no cut in half). At the 10 h the blocks were washed with 1X PBS. A 1 mm thin slide was cut from either the center or the edge of the blocks. For the experimental group, a 1 mm thin slide from the edge was cut and for the control group a 1 mm slide was cut from the center of the block. For the image analysis, we used the straight line and plot profile tools in ImageJ (using command K). The gray scale value obtained is normalized by dividing the values by 255 (denoted as ‘Intensity’ in Figure 3).

#### Flow cytometry

Cytometry was performed with a LSRII Fortessa (BD Biosciences, San Jose, CA). 20 μL of *B. subtilis* cells were transferred to 180 μL 1X PBS with 34 μg/mL of chloramphenicol. For each sample, at least 20,000 events were collected. The data were analyzed using FlowJo (TreeStar, Inc., Ashland, OR). The populations were gated based on forward and side scatter and the geometric mean was recorded. White cells (cells not producing Gfp, but containing a Sp cassette) were run during each experiment to subtract the cellular autofluorescence.

#### Ethanol and NaCl exposure experiments

The printed bars containing spores were exposed them to 100% ethanol and 1.5 M NaCl. We added 25 mL (or until submerged) of either to petri dishes containing the printed parts and covered them for the duration indicated (Supplementary Figure 34). After which, we washed the objects with 12 mL of 1X PBS prior to culturing the cells as described in *Measurement of GFP in 3D Printed Parts*.

#### Experiments with the Ultra violet (UV) light, X-ray and the Gamma Irradiators

(Supplementary Figure 34-37). An Ultraviolet (UV) lamp (Model: ENF-240C; Spectroline™, WestBury, NY) with dual wavelength (365nm and 254nm) was used for the UV experiments. The lamp was held with a stand (Spectroline™ SE140) at 60 mm from the plate with the parts. The X-ray machine used to do this experiment is a repurposed home-built system (MIT RPP X-ray machine). The operational range is from 75-250 kV and the irradiated diameter is ~6 inches. The filter used to perform these experiments was a 0.2 mm Cu filter and we measured the dose rate to be 8.282 R/min using an ion beam (Cat.10X6, RadCal, Monrovia, California). To do the γ-radiation experiments, two gamma irradiators were used: GC-40E (dose rate was 93.16 rad/min) and GC-220E (dose rate was 4375 rad/min in the month of December 2018). The time course was initiated immediately following the exposure period and then the experiments described in *Measurement of GFP in 3D Printed Parts* section were performed (except the LB media was replaced and image taken every 3 h).

## Acknowledgments

LMG and CAV were supported by the Vannevar Bush Faculty Fellowship program sponsored by the Basic Research Office of the Assistant Secretary of Defense for Research and Engineering and funded by the Office of Naval Research through grant number N00014-16-1-2509. This research was also funded by the Institute for Collaborative Biotechnologies (ICB) through contract number W911NF-09-0001 with the U.S. Army Research Office.

## Print files

The name of the STL files of the print designs are provided in the Supplemental Files.

## Data availability

The data that supports the findings of this study are available from the corresponding author upon request.

