## Supplementary Information for "Resilient Living Materials Built By Printing Bacterial Spores"

#### Hacking of a 3D printer system to print cells

##### Overview of the 3D printer system

The MakerBot Replicator 2X (referred to hereafter as "MakerBot") was repurposed to mix and print material and cell streams. Our redesign consists of five subsystems: an agarose pumping subsystem, a cell pumping subsystem, a cooling subsystem, and a heating subsystem (Figure S1). The electronic circuits used to control these subsystems are shown in Figure S2 and S3. These subsystems were designed to be modular. The original pinching and feeding mechanism to reel in polylactic acid (PLA) and acrylonitrile butadiene styrene (ABS) plastic in the original MarkertBot was replaced by a printhead where agarose and cells blend prior to being extruded out of the nozzle. The cells and agarose are propelled to the printhead from two separate reservoirs. This design allows the cells to be exposed to elevated temperature for a short period of time (< 20 min). This design permits us to use an established skeinforge gcode generator with minor modifications in the temperature and infill settings. In practice, the biggest challenge in building the 3D Printer was pumping the 5% agarose and balancing of the temperature in the heating and cooling subsystems. The pumping rate varies as a function of temperature and pressure. Pumping the heavy agarose required high pressure to prime the lines and reach the printhead and for this reason we used compressed air. The flow rate of the agarose was controlled using a relief electronic valve and by adding a check valve that adds ~1 psi of resistance. This maintains the pressure at ~0.75 psi. In the printhead, the temperature was kept constant at 75°C using a PID controller. Peltier plates were used to extend the time the print chamber could be kept cool. The chamber needs to be cold for the agarose to solidify at the instant it extrudes from the nozzle. Cooling the chamber allowed the agarose to retain its filamentary shape while it exits the nozzle. A set of 3 Arduino boards were used to drive the printer. One Arduino board (Arduino Mega) was used to control the agarose and cell pumping subsystems, another Arduino microcontroller (Arduino Uno) was used to control the scaffolding subsystem and an RFIArduino was used for controlling the temperature in the mixing chamber (Supplementary Table 1). In Supplementary Table 2 and Table 3, we list all of the parts used to build this 3D printer system.

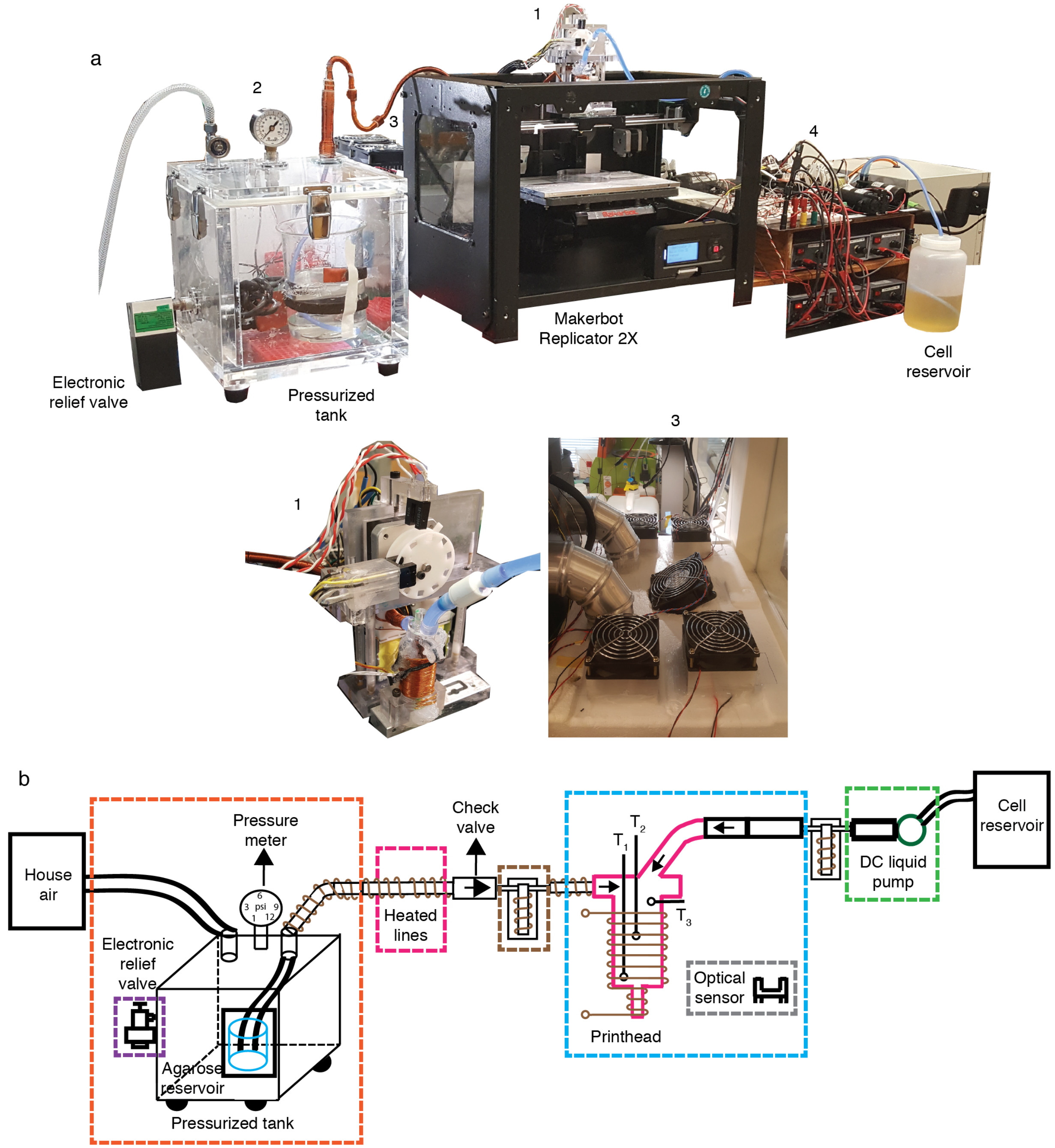

**<u>Figure S1.</u> 3D printer design**. **a,** Picture of the 3D printer and its subsystems: 1. printhead, 2. agarose pumping subsystem, 3. cooling subsystem, 4. electronic boards (Figure S2 and S3). **b,** Schematic showing the flow of liquids from the agarose and cell reservoirs. Purple box: an electronic relief valve to keep the air pressure in the pressurized tank constant. Pink Box: the heating device to maintain the agarose in the molten state while reaching the printhead from the reservoir. Orange box: a pressurized tank with one inlet for the incoming air and one outlet for the agarose to reach the printhead. Brown box: a solenoid valve (normally closed) that when energized opens to allow the flow of agarose from the lines into the printhead. Cyan box: the printhead where the cells and agarose mix and carries the optical sensors. Grey box: the optical sensor to detect which extruder is functioning (for dual printing) and when to input cells. Green box: the DC liquid pump used to actuate the cells to the printhead.

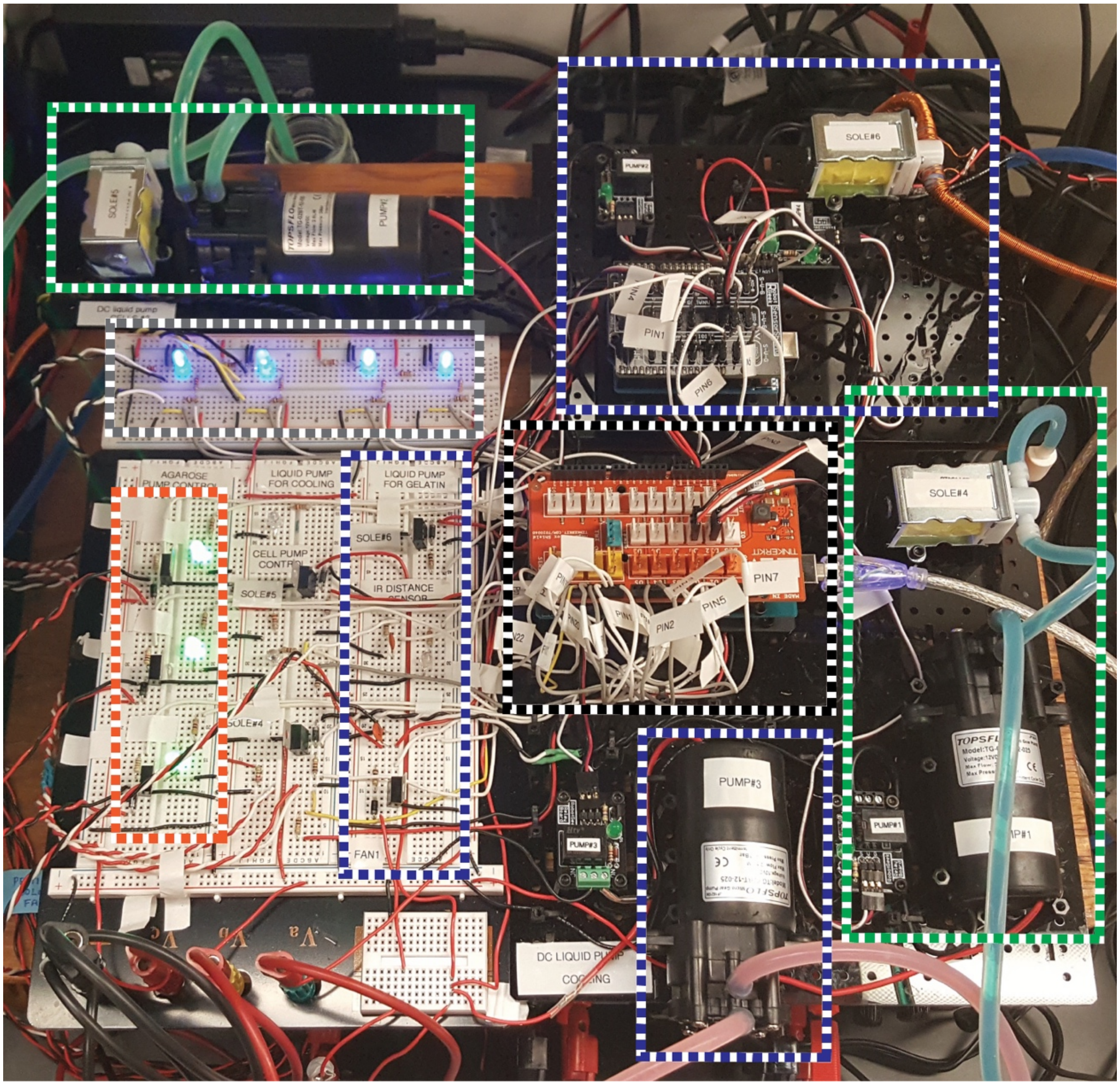

**<u>Figure S2.</u> Electronic circuit boards and DC liquid pumps**. The color scheme corresponds with Figure 1S. The green dotted squares show the cell liquid pumps. The dotted orange square shows the solenoids controlling the agarose input (the circuit controlling electronic relief valve is not shown here). The gray dotted box shows the circuit that controls the optical sensors. The black dotted box in the middle shows the main Arduino board wiring. Figures 6S, 8S and 10S include detailed wiring diagram of all of the circuits.

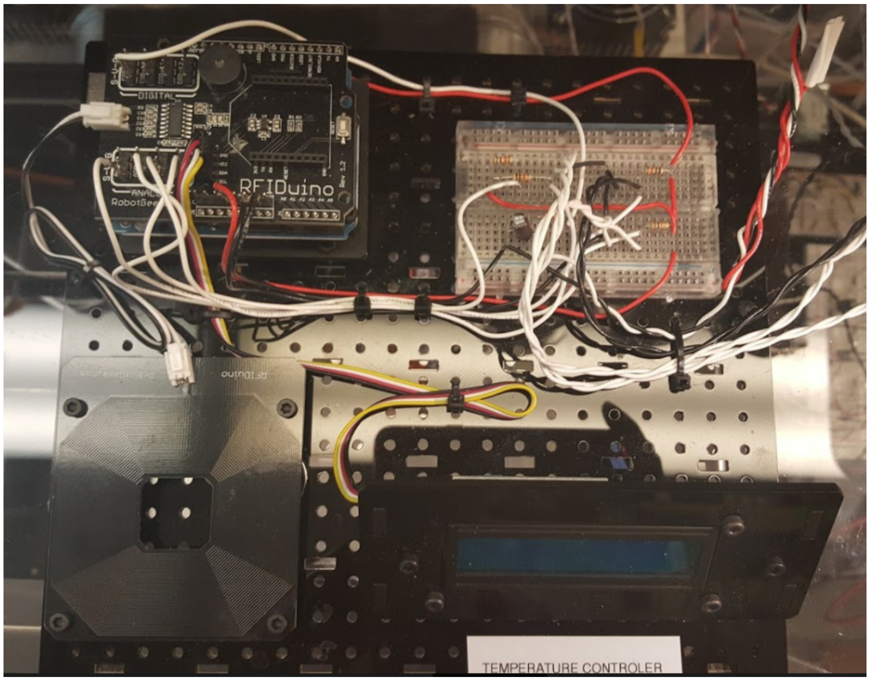

**<u>Figure S3</u>. Electronic circuit board for controlling the temperature.** This shows the LCD that displays the temperature in the chamber. An RFIDuino module was used to set-up this circuit. Figure S13 shows the detailed wiring diagram corresponding to the circuit board.

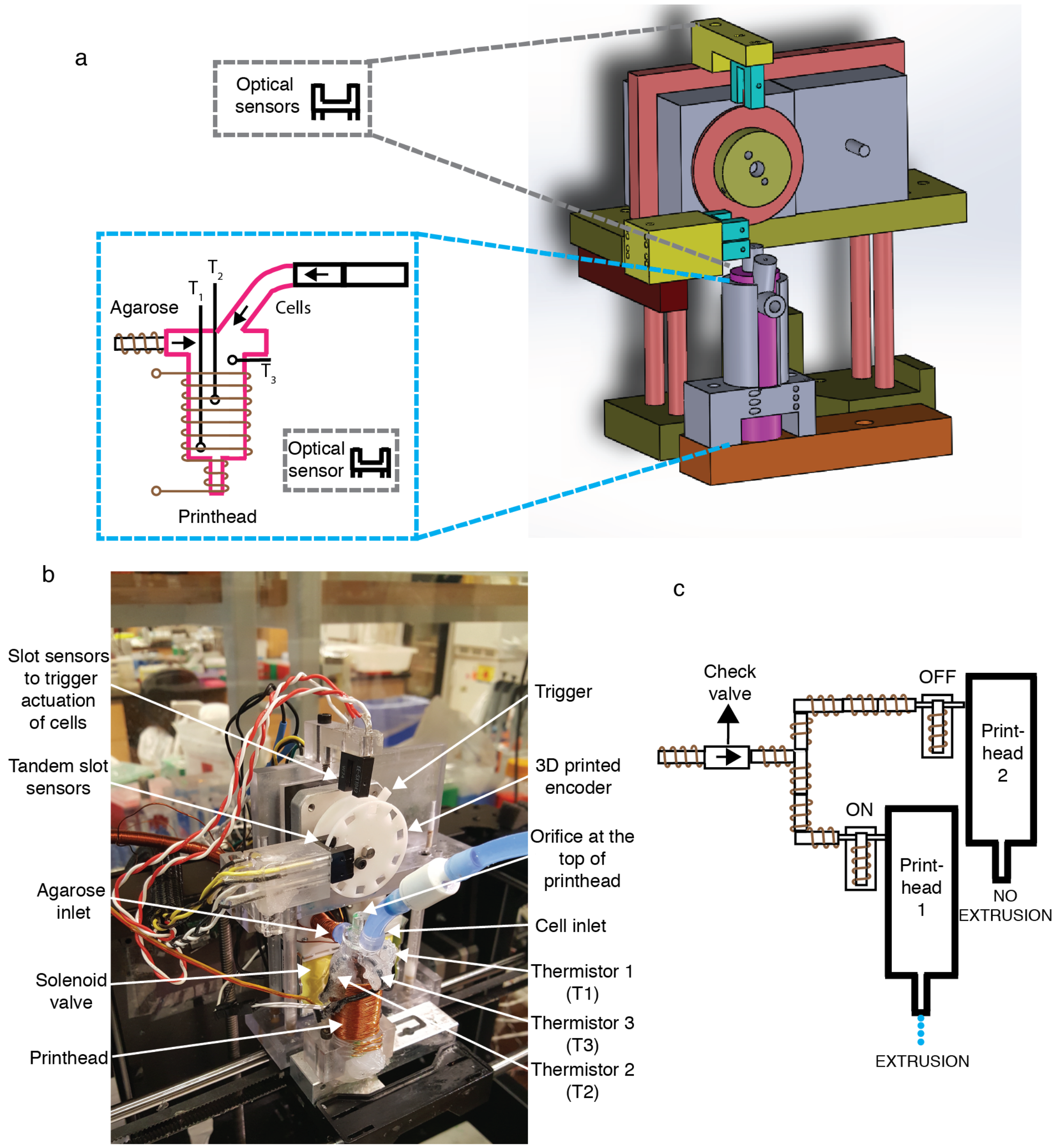

**<u>Figure S4.</u> Printhead and optical sensors subsystem**. The original MakerBot printhead was replaced with a new one; designed to be able to handle a different polymer, agarose. **a**, Schematic of the printhead and optical sensors. The printhead's barrel has two inlets, one for the agarose and for the cells. The insect shows the thermistors' positions (a thermistor is a resistor that changes resistance with temperature). The three thermistor sensors are used to measure the temperature at the top, middle and bottom of the printhead, labelled as $T_1$, $T_2$, $T_3$, respectively. The average of $T_1$ and $T_2$ is used is compared to a setpoint to maintain the temperature in the printhead at 75°C to ensure mixing of the agarose and spores. The temperature $T_3$ is used to monitor the temperature at the top of the printhead to indicate when to input more agarose. **b**, Picture of the printhead showing the different components. A pair of optical interrupters (labelled tandem slot sensors). A single optical interrupter was used to input the cells about every 60 seconds or the time it takes for two revolutions to occur. An orifice (1mm in diameter) was introduced at the top of the printhead for excess air to escape after priming and while printing as house air at the agarose cabinet (see the agarose pump system) is constantly purged into it. This modification is a key feature that relieved the problem with having bubbles inside the print that distorted the parts and disrupted the uniformity in the printing. **c**, Optical interrupts were coupled to stepper motors (of the MakerBot) to switch between the solenoid valves of the pumping system. This pair of optical sensors are also used to detect when the motor reverse direction.

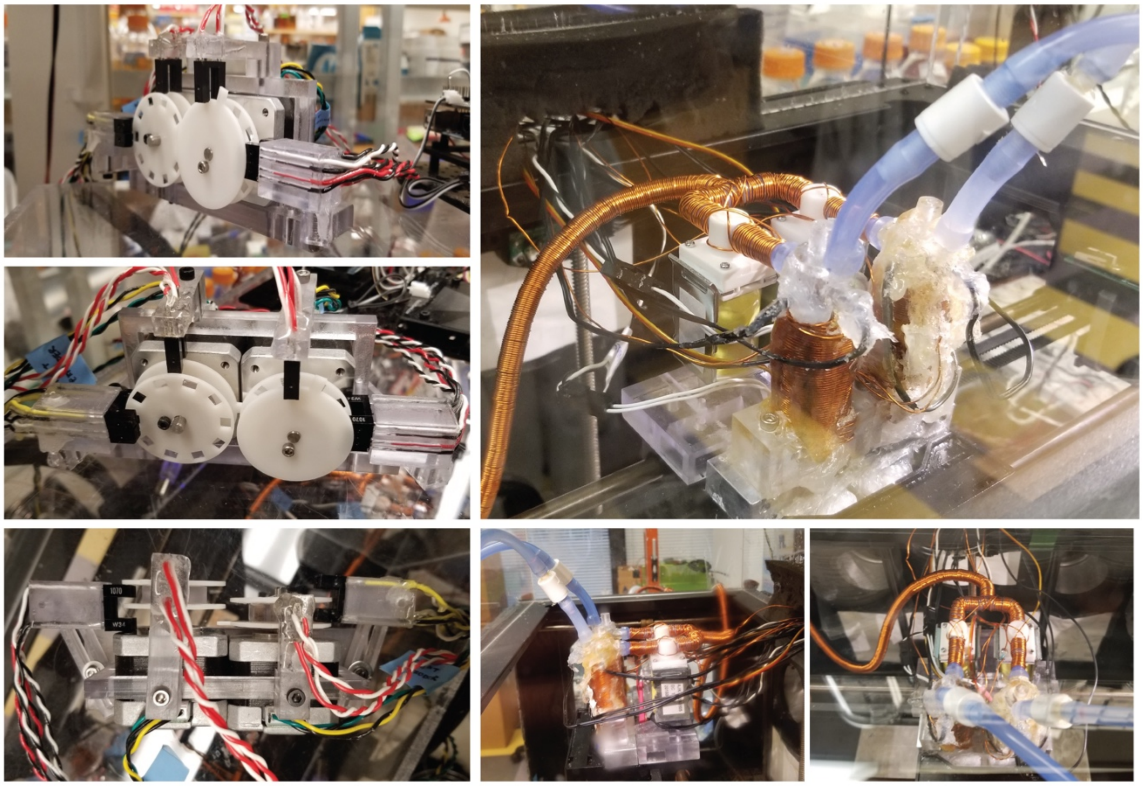

**<u>Figure S5</u>. Dual printing configuration.** These pictures were taken of the same system at different angles. The agarose line was split to enter two separate solenoid valves using a barbed tube fitting tee connector. These solenoid valves are couple to the original MarkerBot's stepper motors via the tandem optical sensors. This set-up was used to print a hybrid material with cells on one set of layers and only agarose in a second set of layers. Individual gcodes are merged using the Merge tool in ReplicatorG.

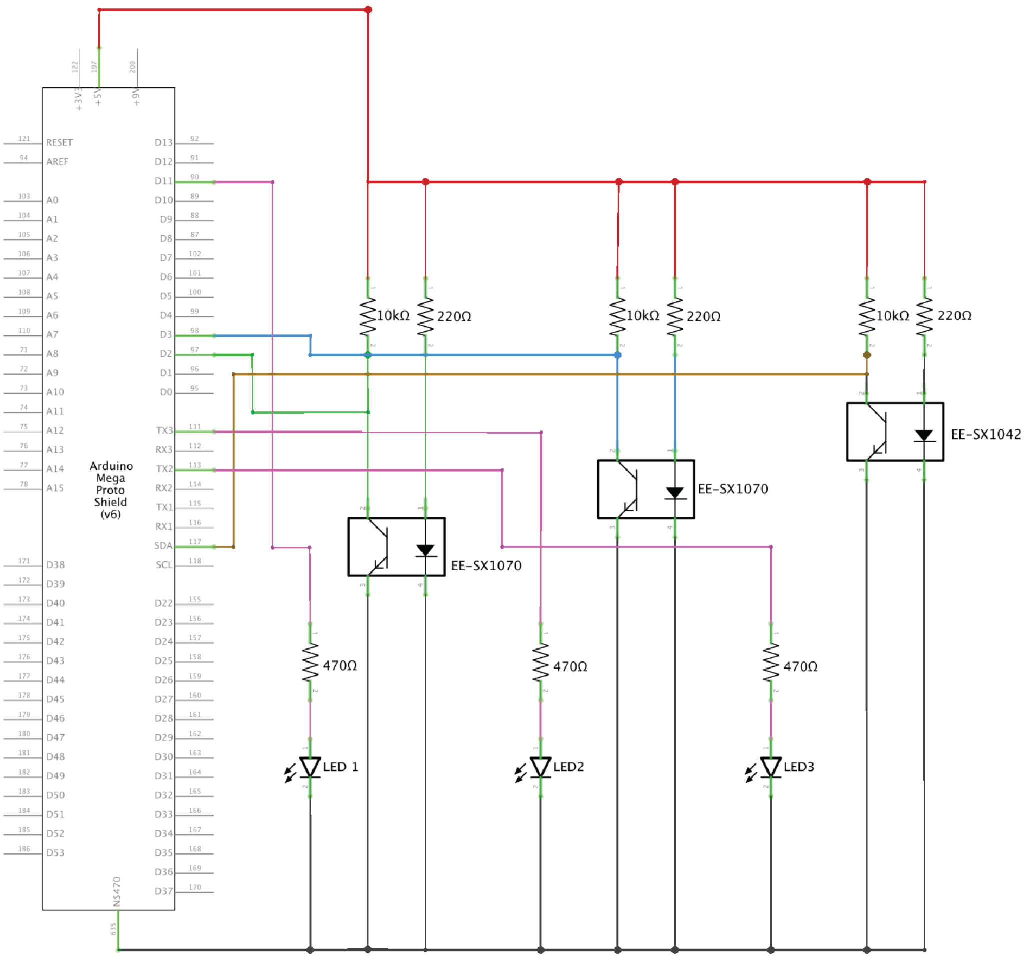

**Figure S6. Electronic circuit diagram of the optical sensors**. This shows the wiring for each optical sensor. A pair of tandem optical sensors (part no. EE-SX1070) connected to digital pins 2 and 3 is used to detect reversal in direction of the motor (to close the valves) and to indicate which stepper motor is working at a given time (for dual printing; Fig. S4 and S5). The single optical sensor (part no. EE-SX1042) is used for the timed input of cells and it is couple to the MakerBot's stepper motor. All of the optical sensors use interrupt functions as shown in the code. This wiring also shows the LED indicators for each optical sensor to indicate which sensor is active at a given time. This diagram only shows sensors for one of the printheads. See Supplementary Table 1 for the other pin assignments and Supplementary Table 2 for more details on the parts used.

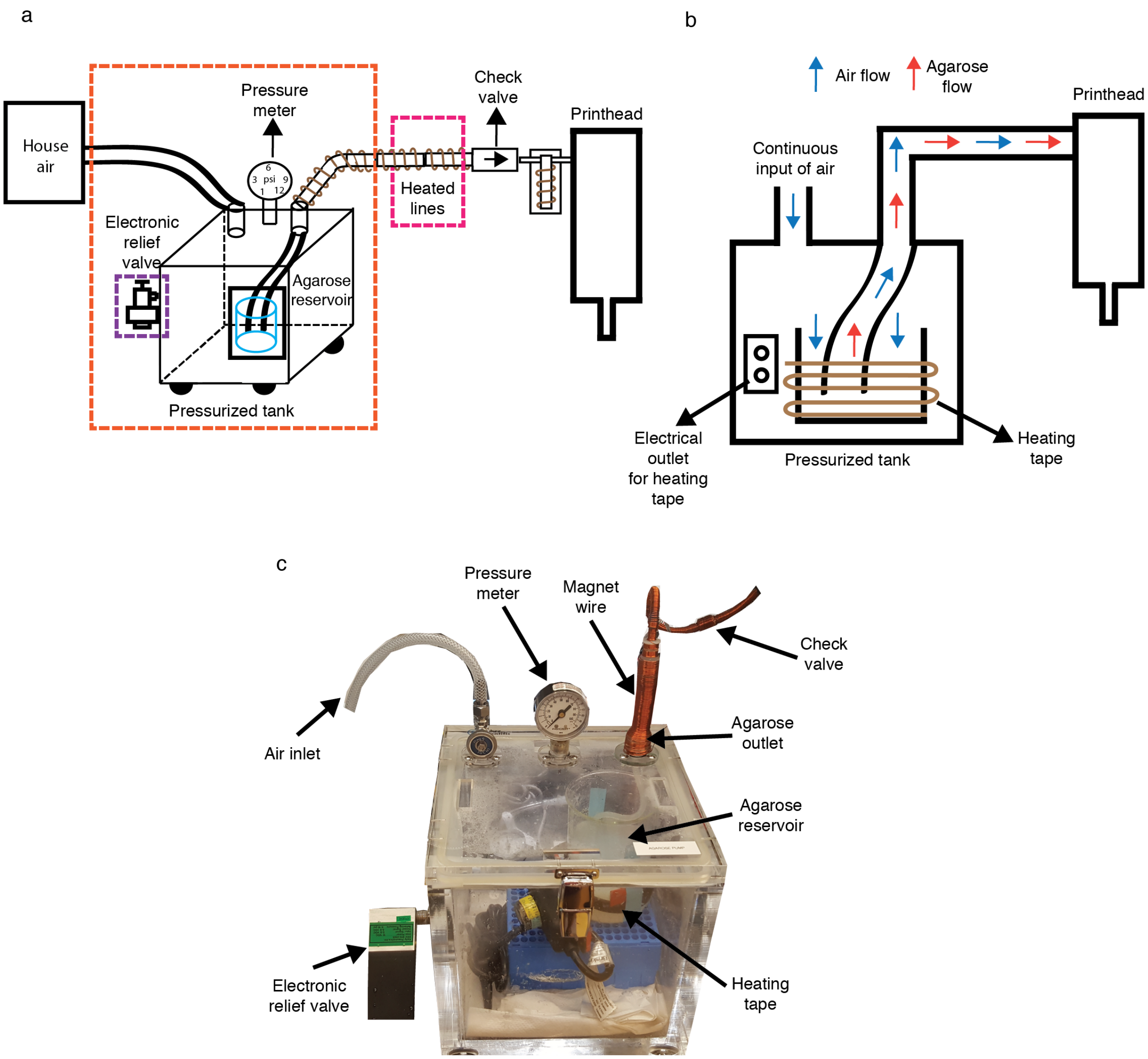

**Figure S7. Mechanical configuration of the agarose pumping subsystem. a**, Schematic of the agarose pump subsystem. The agarose is pumped to the printhead and there it mixes with the cells. **b,** Air and agarose flow within the pressurized tank for applying positive pressure to actuate the agarose to the printhead. **c,** Picture of agarose tank showing relief valve, heating tape, agarose reservoir, check valves, magnet wire and pressure meter. For this agarose printing system, we used a hermetically sealed desiccator cabinet (originally used to create a vacuum) that we turned into a pump; with a ¾ in thick wall made of polymethylmethacrylate (PMMA). The vacuum gauge was replaced with a pressure gauge, with a range of 0-15 psi to monitor and maintain a constant pressure in the chamber. To maintain the agarose flowing a magnet wire (AWG 21) is wound around the plastic tubing as passing current through a conductor releases heat (Joule heating). For the tubing, we used a high temperature silicone rubber. In addition, the cabinet/chamber has an electrical outlet used to plug a heating tape for maintaining the agarose melted in the reservoir. There is an o-ring (size 114) in the inlet and outlet of the lid of the cabinet to prevent leakage. In addition, a check valve prevents the flow of this viscous liquid back into the reservoir. This check valve adds about 1 psi of resistance to the line. Adding more check valves causes problem with severe clogging of the line and is hard to clean. An electronic relief valve was installed at the chamber to immediately, within a few milliseconds, relieve the pressure when it is not needed. Essentially, we initially primed the line and chamber with molten agarose keeping the relief valve closed. The pressure used is adjusted and pulsed on and off so that the pressure is about 0.75 psi. A pair of solenoid valves is placed between the check valve and the printhead. Fig. S8 show a schematic of the electronic circuit to control the pumping of the agarose. Supplementary Table 2 contains the details of all of the parts used.

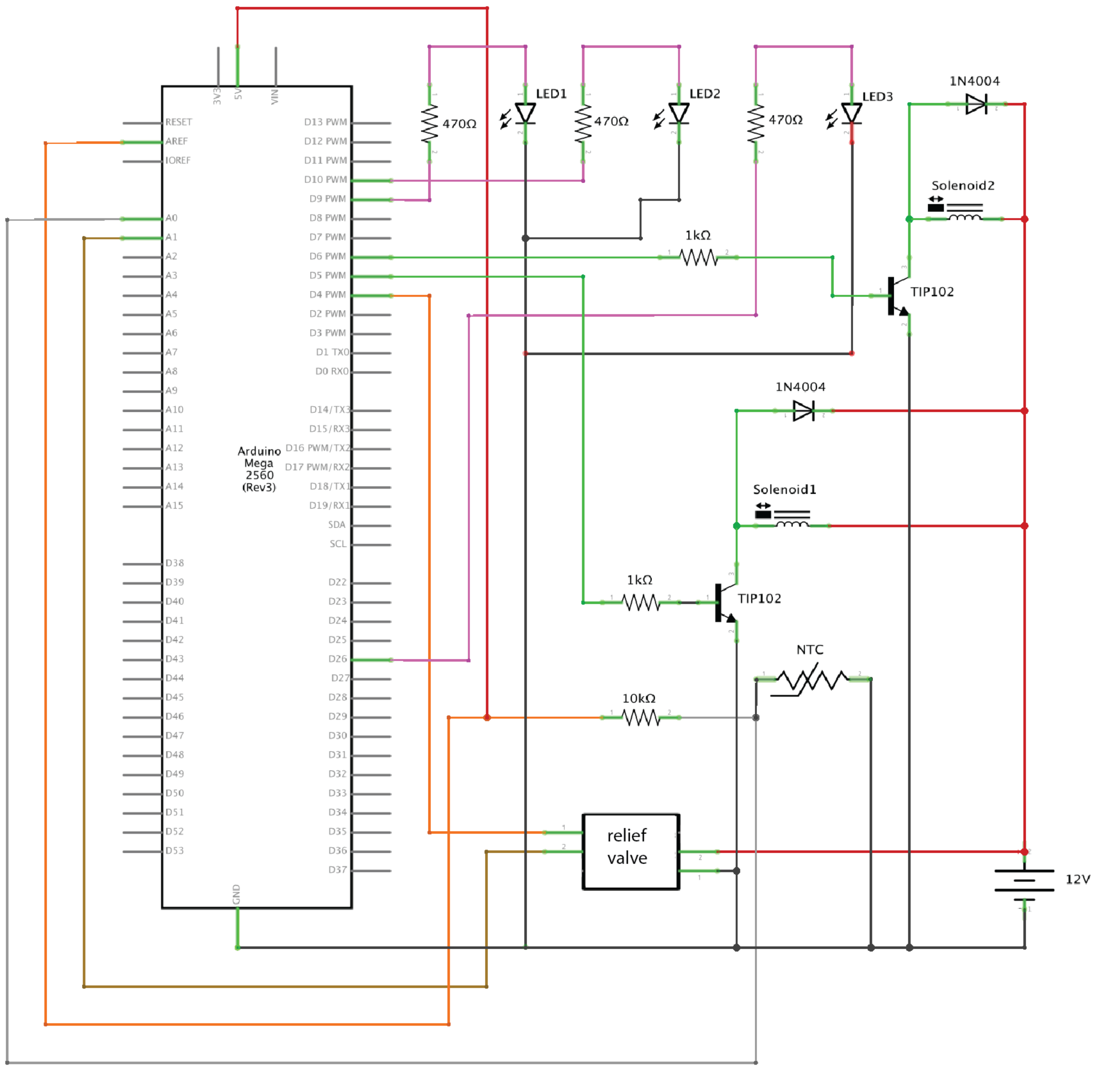

**Figure S8. Electronic circuit diagram to control the agarose pumping subsystem**. This wiring shows the LED indicators for each solenoid valve. It highlights the relief valve's input, output, power and ground wires. It shows the thermistor (3950 NTC) and solenoid valves circuitry. Power transistor (TIP102) and diode (1N4004) are additional elements used to control the solenoid valves. See Supplementary Table 1 for the pin assignment and Supplementary Table 2 for details of all of the parts used.

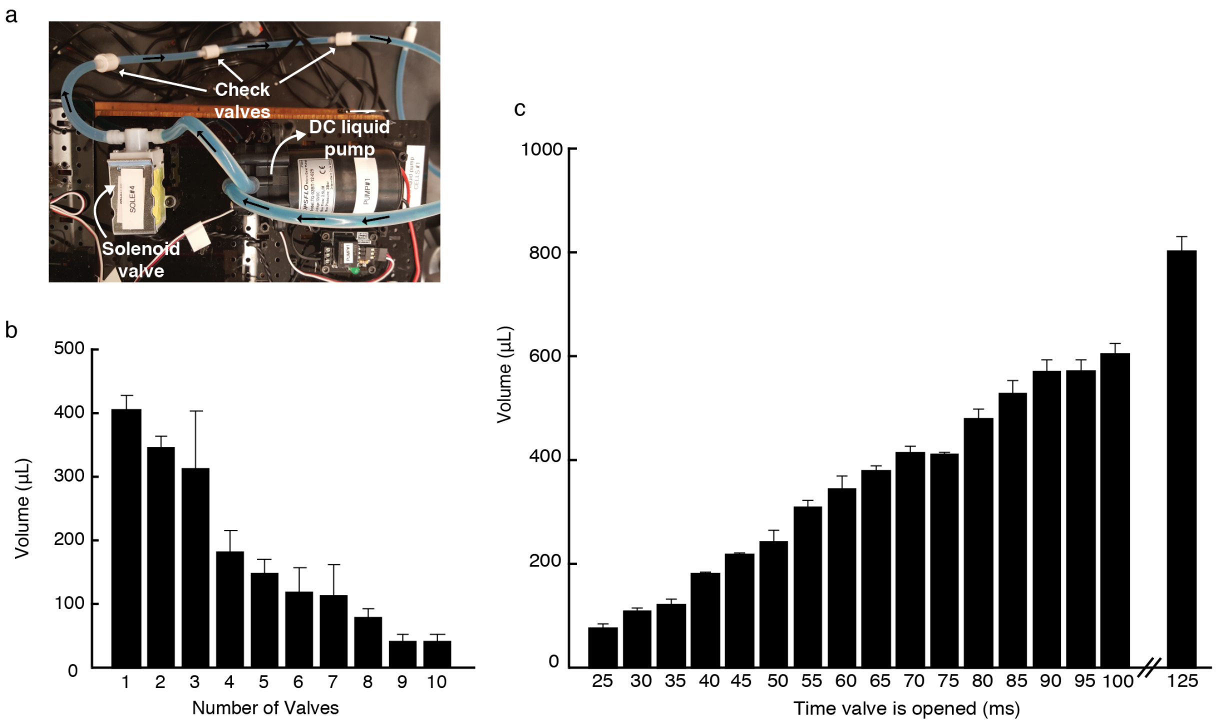

**<u>Figure S9.</u> Cell input subsystem for initial priming. a**, Picture of a solenoid valve, a DC liquid pump and check valves (only 3 out of 8 are shown in this picture) used to control the cell input subsystem. The flowrate of the DC liquid pump without any check valves is 2 L/min or 33.3 mL/sec. Check valves were added to decrease the flow rate and determine how much cells to add so that the agarose percentage is greater than 4%. The power supply (power supplies 3-12V, 2A) used was fixed at 9V and 2A for all of these calibrations. **b,** Calibration of the flowrate to pump cells by adding check valves using a fixed 25 ms delay (the time the solenoid is energized; or opened). **c,** Eight check valves were used for this calibration and the time the solenoid valve was energized was varied. After priming the lines with agarose and cells, the printhead contains ~5 mL of agarose and we input ~600 uL of cells (90 ms delay). See Supplementary Table 2 for the details of all of the parts used. The means of three replicates are shown and the error bars are the standard deviation.

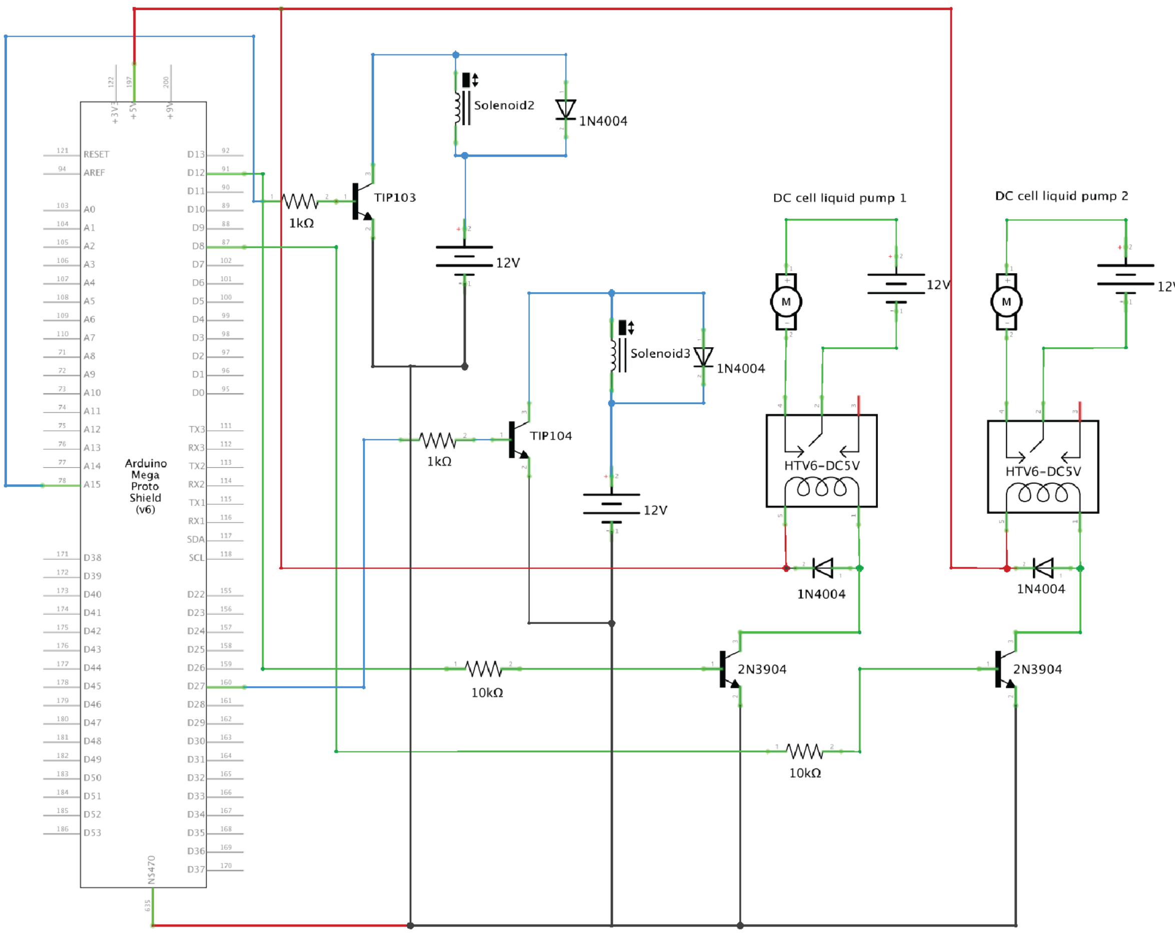

**<u>Figure S10</u>. Electronic circuit diagram to control DC liquid pumps and solenoid valves.** This circuit controls the quantity of cells added to the printhead to maintain a constant agarose percentage greater than 4%. The solenoid valves are opened at the same time the DC liquid pumps are opened. The solenoid valves are used as a safety to prevent any liquid from entering as we are using diaphragm pumps. This wiring diagram shows the connection of the RobotGeek Relay board (labelled HTV6-DC5V) with diode (1N4004) and NPN bipolar junction transistor, 2N3904. The corresponding solenoid valves circuitry is also shown. Supplementary Table 1 has details on all of the pin assignments and Supplementary Table 2 contains details of all of the parts used.

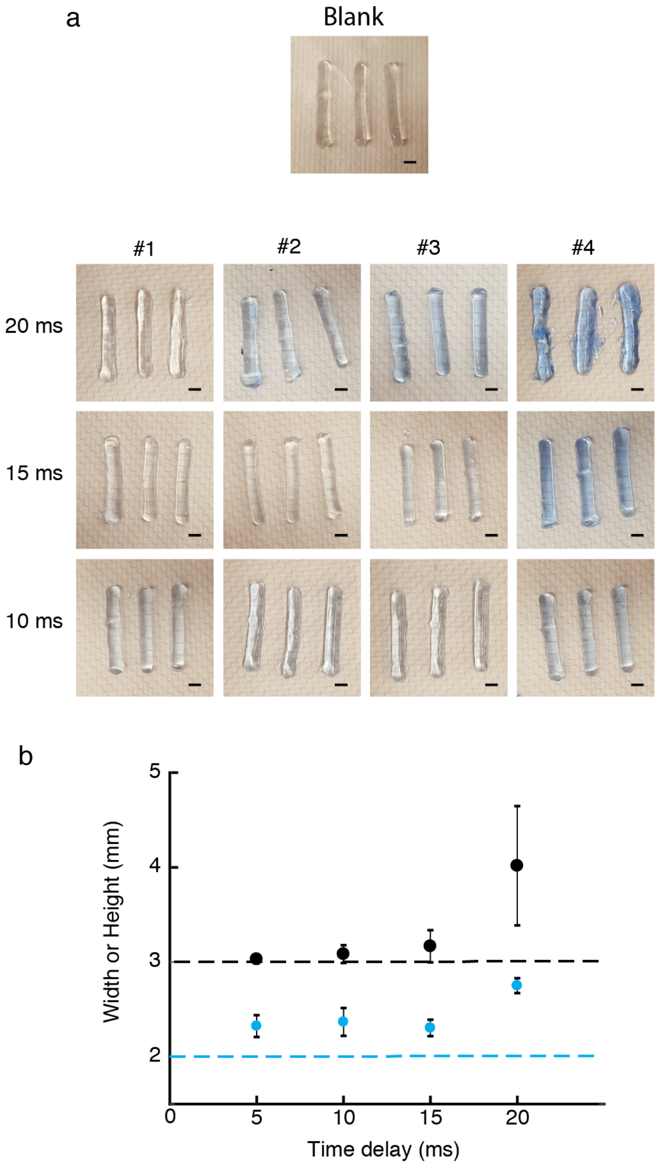

**Figure S11. Cell input subsystem for continuous printing.** After the initially filling the printhead with agarose and cells (**Fig. S9**) the cells added for ~100 μL (amount of agarose entering the printihead when one solenoid valves is opened) of agarose needed to be calculated. Using the 8 check valves and the power supply set at a 9V and 2A, we determined the time to keep the pump opened. **a**, Instead of cells, we used water with a blue indicator dye. We printed 3 bars (with dimensions 2 x 3 x 25 mm) 4 times in consecutive runs and the number on the top row indicates the order in which they were printed (#1 to #4). The column number is the time the pump and solenoid valves are opened. **b**, Plot of the measured width and the height of the printed bars. The black (blue) dashed lines highlight the target width (height) of the printed bars. Fifteen or ten ms would work. We choose a 15 ms delay for continuous printing. The means of three replicates are shown and the error bars are the standard deviation. Scale bars, 3 mm.

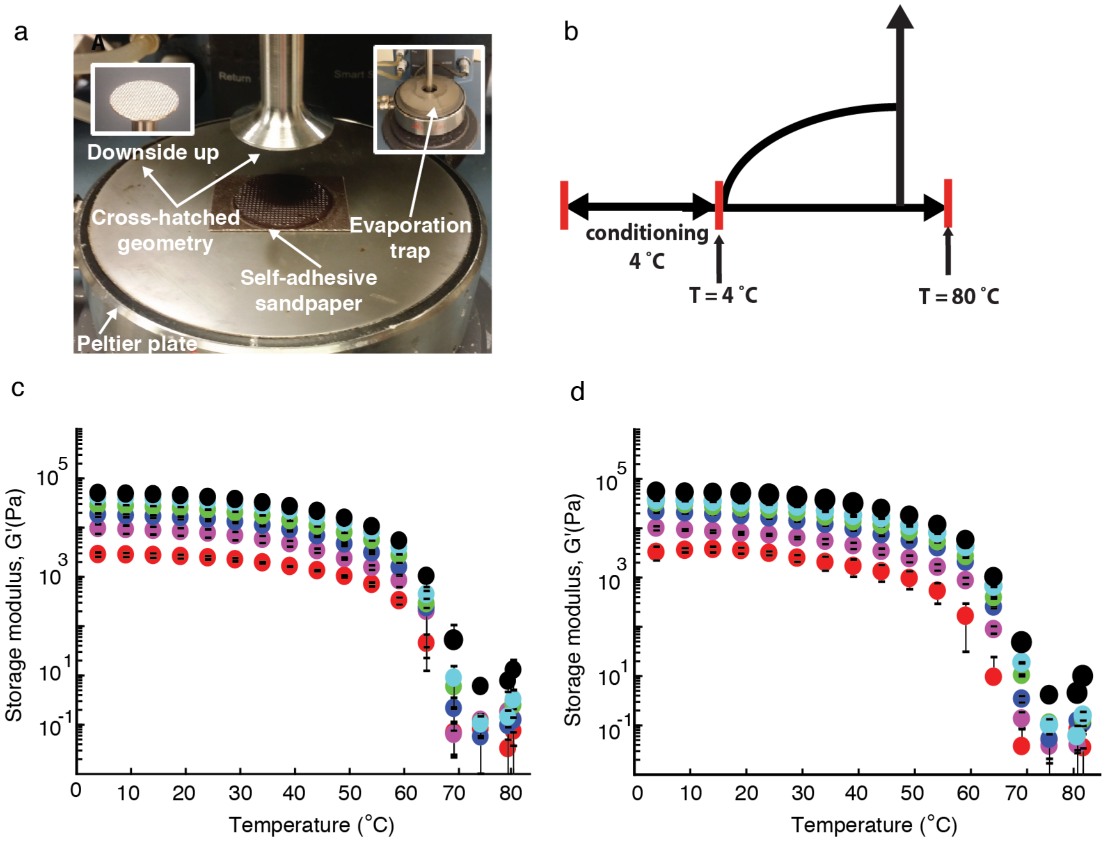

**Figure S12. Melting temperature of the SeaPlaque agarose**. **a**, Rheometer set-up to do this experiment using a Peltier plate, an evaporation trap, a cross-hatch geometry and self-adhesive sandpaper. **b**, A schematic showing the conditioning step (temperature at 4°C prior to the start of data acquisition) and increasing the temperature from 4°C to 80°C. Plots showing the storage modulus, G', of various agarose percentages (1%-red, 2%-magenta, 3%-blue, 4%-green, 5%-cyan, 6%-black) for the agarose **(c)** and the UltraPure agarose **(d)**. There is no difference in melting temperature between the SeaPlaque and the UltraPure agarose. There is no difference in melting temperature between the SeaPlaque and the UltraPure agarose. The datapoints show the means from three replicates and the error bars are the standard deviations. See Methods for more experimental details.

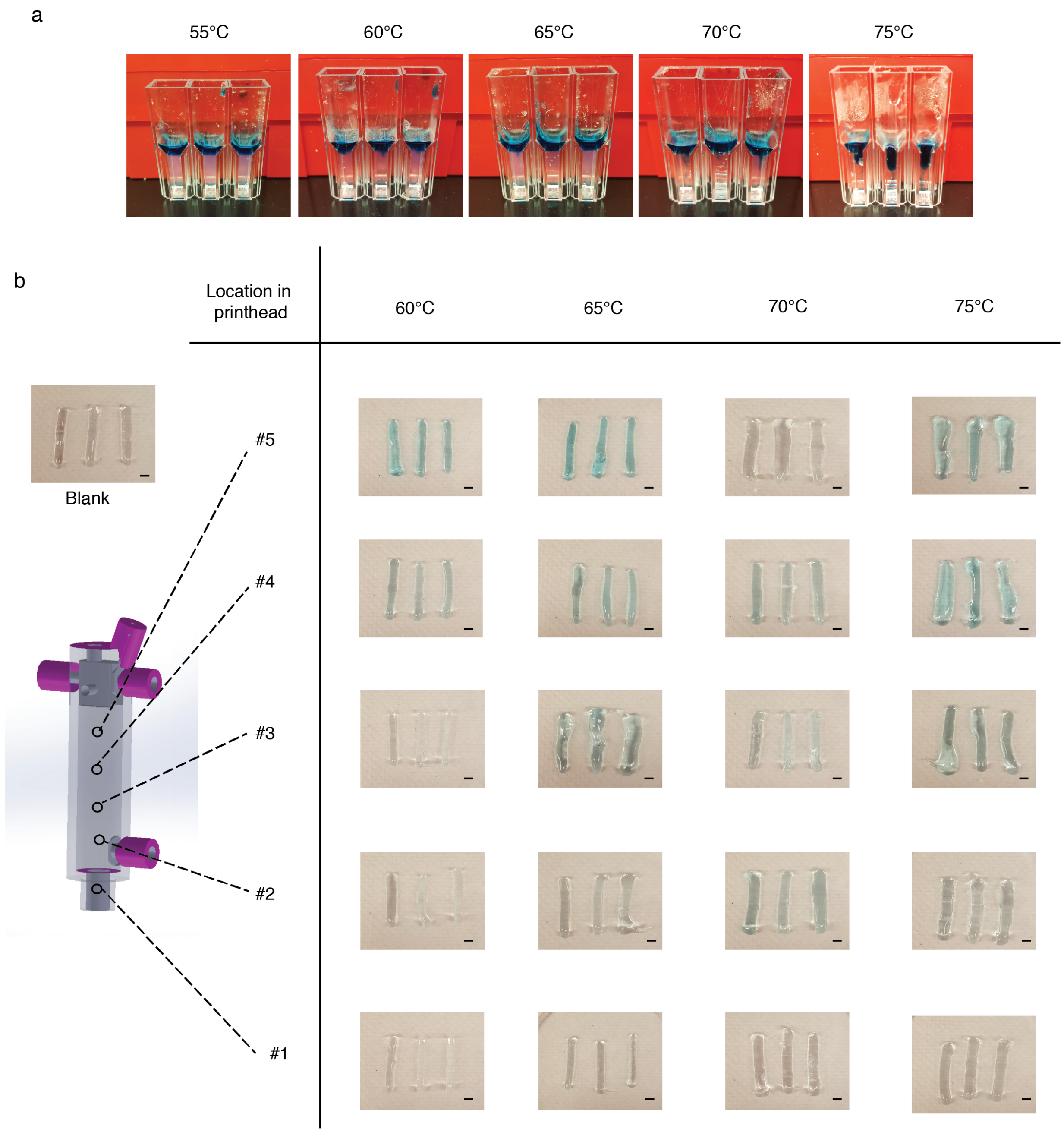

**Figure S13. Temperature dependence for mixing agarose and cell streams**. **a**, Heating the 5% agarose outside the printhead and injecting water with a blue dye to see how much the water penetrates at each temperature. **b**, Heating agarose inside printhead and accessing how well the agarose and blue water are mixing. This experiment was done while the printing chamber was at room temperature ($T_{ambient}$ = 22°C). Based on the agarose blue color present on the printed bar on the second row (#2) from the bottom, we concluded that either 70 or 75°C would work. The temperature at the printhead must be about 75°C to allow sufficient mixing based on the two independent experiments depicted in **a** and **b**. Scale bar, 3 mm.

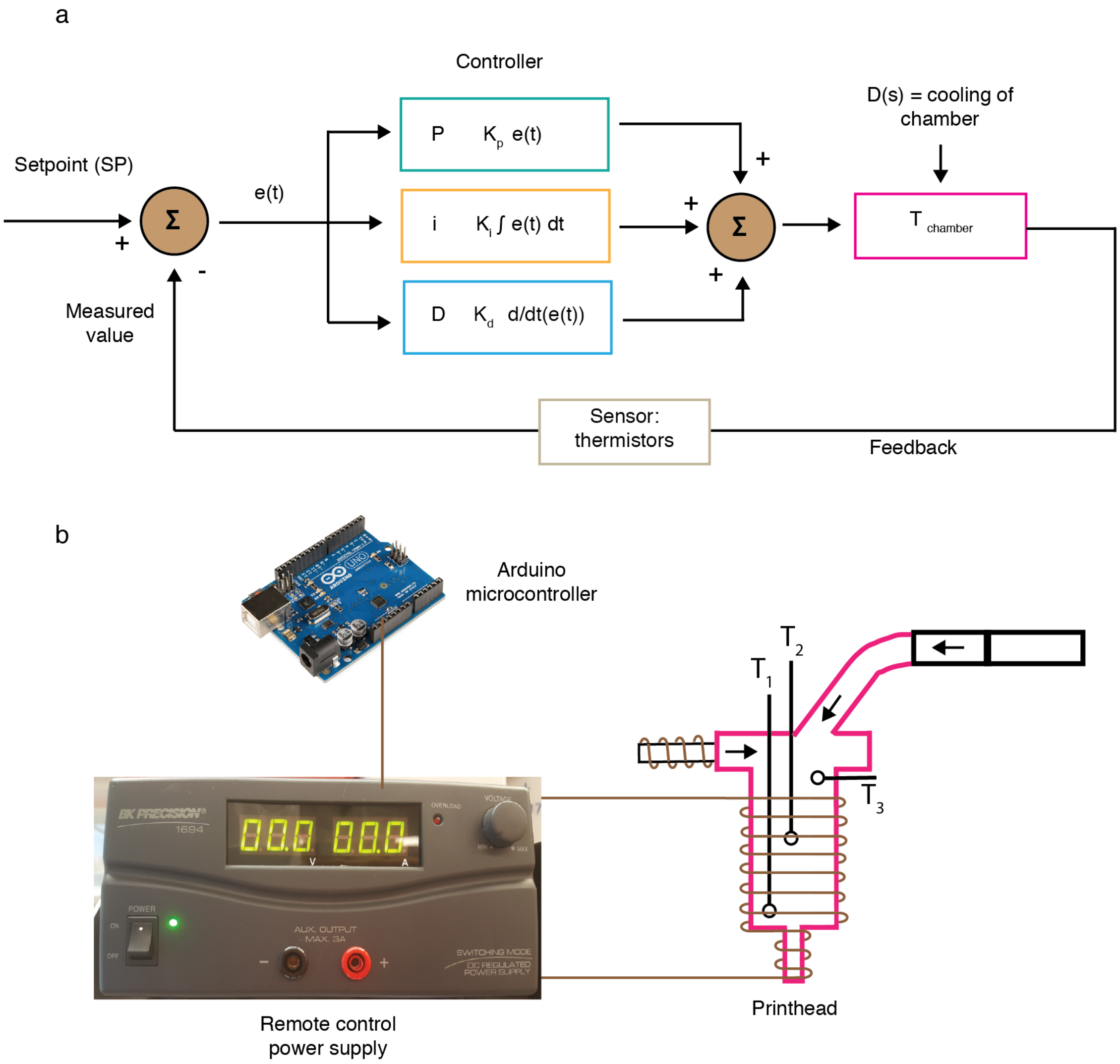

**Figure S14. Implementation of a PID controller using a remote-controlled power supply.** The temperature in the mixing chamber where the cells and agarose come together is maintained at a constant temperature of 75°C to ensure mixing the cells with the agarose. **a**, Diagram depicting the PID controller, sensors, disturbances and feedback. In a PID algorithm, the measured temperature (the average of $T_1$ and $T_2$ measured via two thermistors) is compared to a setpoint and the difference between them is the error. The PID controller ameliorates the problem of having disturbances that affects the output, one of the being the cooling of the chamber. The parameters and equation for the PID controller are provided in the Methods. **b**, Connection of the printhead, a remote-controlled power supply and an Arduino microcontroller. See Figure S13 for the circuit board corresponding to this heating subsystem and Supplementary Table 2 for the parts used.

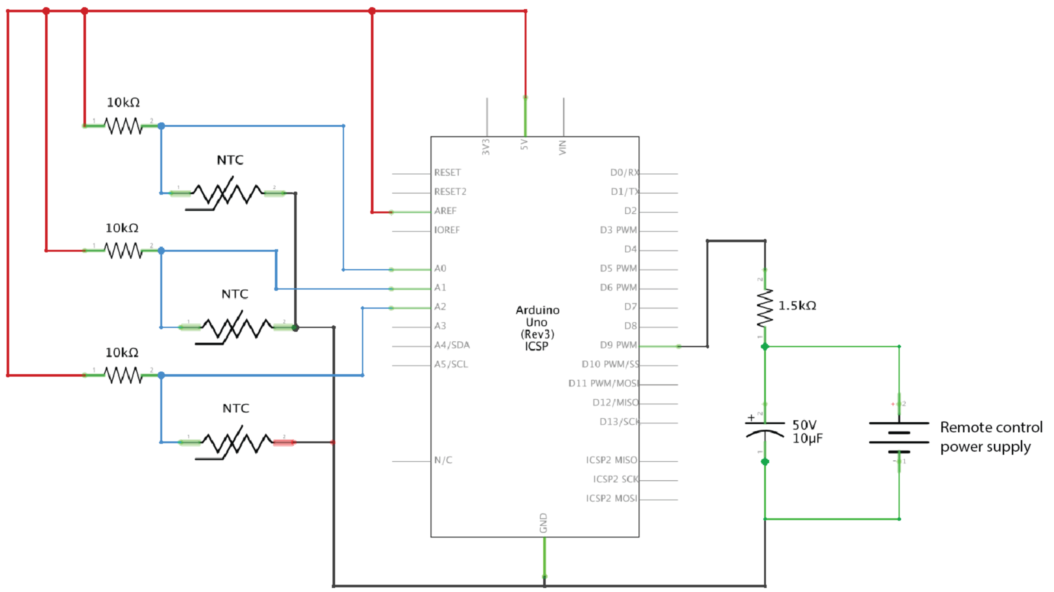

__Figure S15.__ **Electronic circuit diagram of the temperature controller.** This shows the wiring for each thermistor and the connection to the remote power supply (Figure S12). Supplementary Table 2 contains details of all of the parts used.

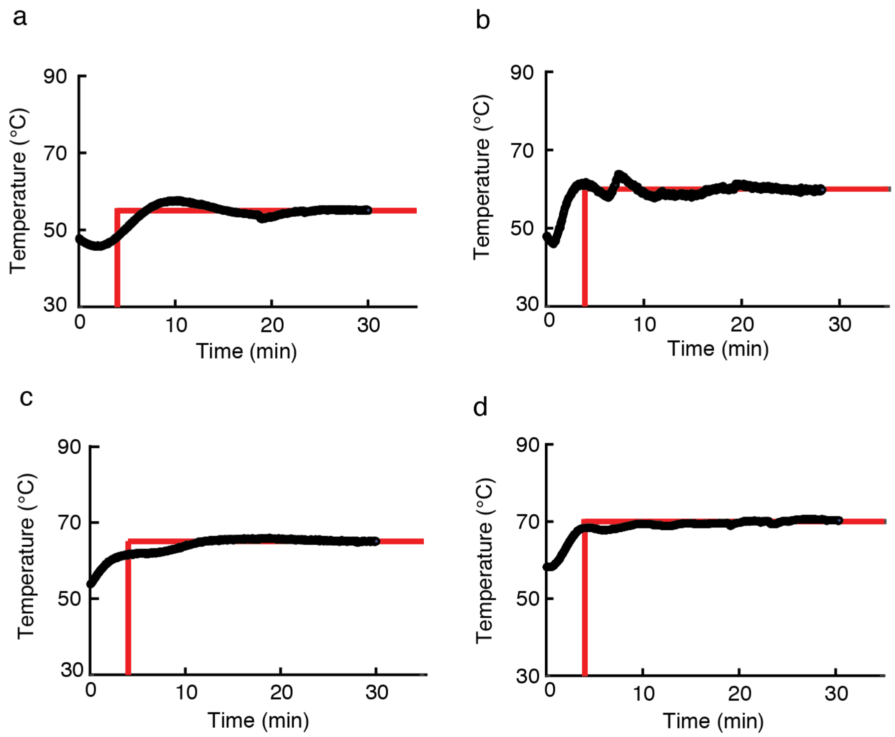

**<u>Figure S16.</u> Testing the PID controller at room temperature (**$T_{ambient}$ = 22°C**).** The setpoint shown in red for each graph are the following: **a,** $T_{chamber}$ = 55°C**; b,** $T_{chamber}$ = 60°C**; c,** $T_{chamber}$ = 65°C; d, $T_{chamber}$ = 70°C. The measure value is shown in black and is takes 5-10 min for it to reach the setpoint value. Figure S12 and S13 show the set-up and circuit diagram used before testing the PID controller.

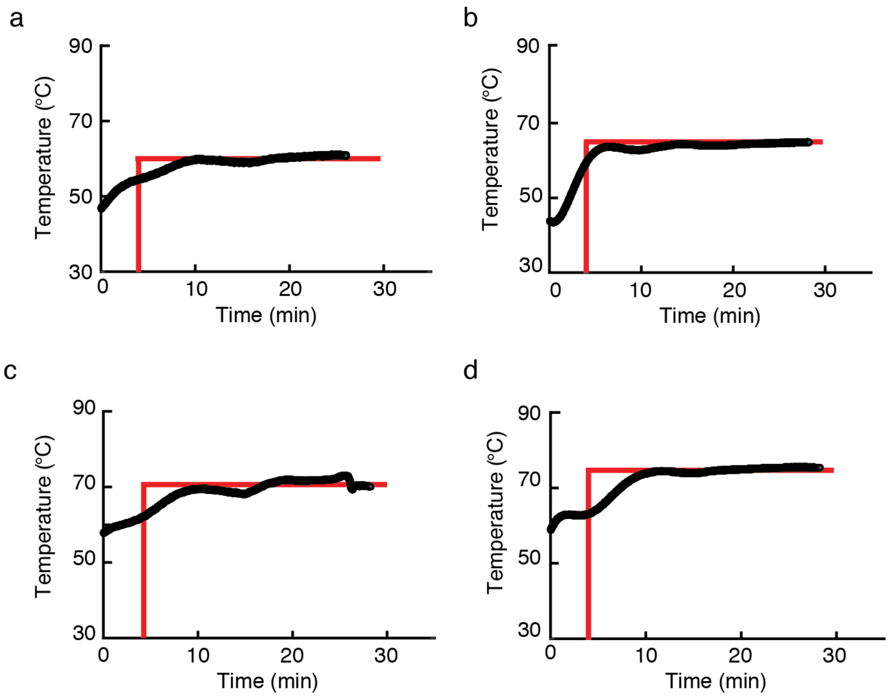

**<u>Figure S17.</u> Testing the PID controller using a cool printing chamber** ($T_{ambient}$ = 16°C)**.** The setpoint shown in red for each graph are the following: **a,** $T_{chamber}$ = 60°C**. b,** $T_{chamber}$ = 65°C**. c,** $T_{chamber}$ = 70°C**. d,** $T_{chamber}$ = 75°C**.** The measure value is shown in black and is takes about 10 min for it to reach the setpoint value. Figure S12 and S13 show the set-up and circuit diagram used before testing the PID controller.

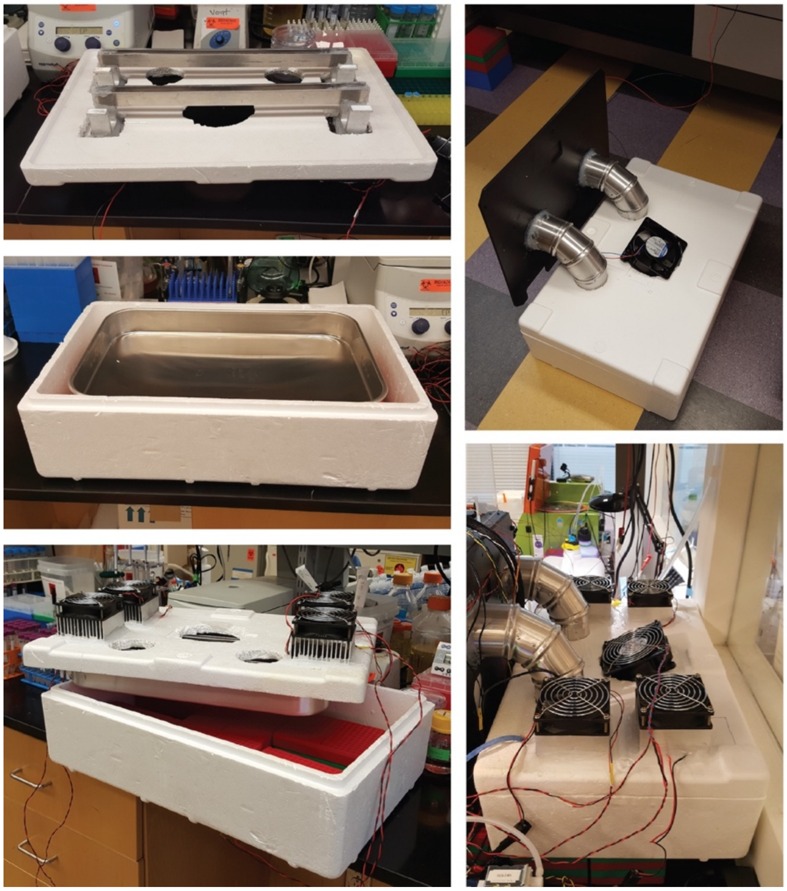

**<u>Figure S18</u>. Construction of the cooling subsystem.** To facilitate rapid gelation of the agarose, the print chamber was maintained between 14-17°C by blowing cold air into it. In the back of the printer, we placed a Styrofoam box with an aluminum pan. Two metal bars that span across this pan are attached to Peltier plates to aid in maintaining the temperature. In addition, metal fins were attached to the metal bars to increase the surface area and to aid cool the air transfer into the chamber. Two aluminum elbow pipes were used to guide the cold air into the printing chamber. A fan position in the middle of the set-up blows the cold air into this chamber.

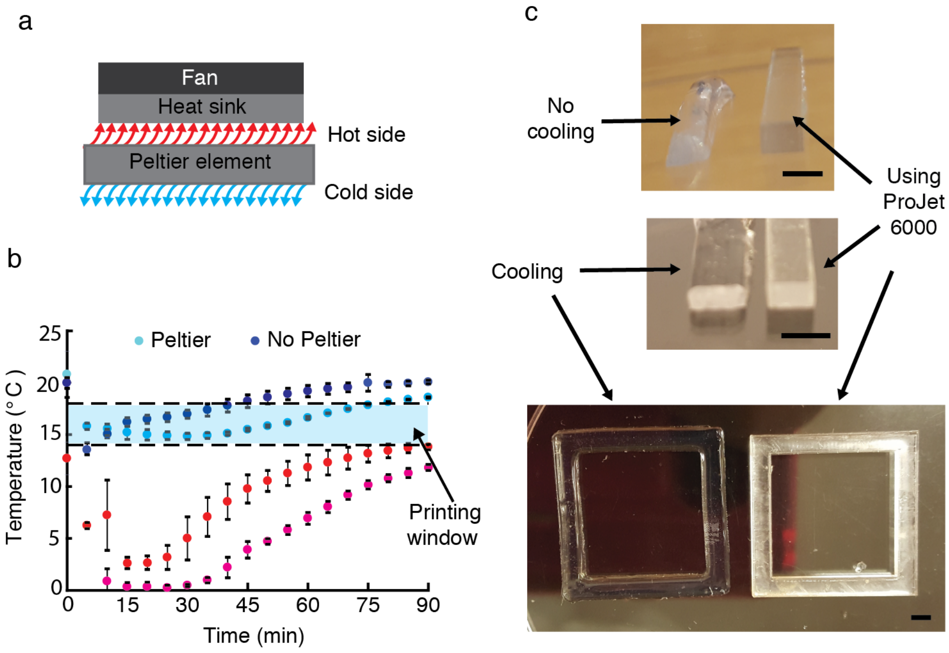

**Figure S19**. **Testing the cooling system.** **a**, Illustration of the Peltier concept. **b**, Temperature in the printing chamber (blue and cyan) and the ice chamber (red, magenta). Printing window is the temperature at which deformation of the printed object is minimized. When the temperature in the printing chamber is greater than 18℃. **c**, Printing a rod and a square bar implementing cooling and comparison with something printed with a high resolution commercially available printer, the ProJet 6000 (3D Systems). Scale bar, 3 mm.

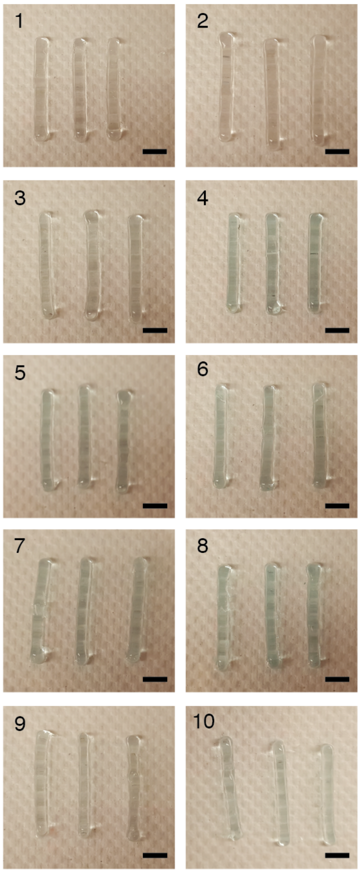

**<u>Figure S20</u>. Improving the quality of the 3D printed parts**. Printed bars (2 x 3 x 25 mm) after cooling and after introducing a 1 mm small orifice at the topmost part of the printhead. This small orifice offers an escape for the bubbles in the agarose material. $T_{ambient}$ was ~16°C for this experiment. The upper left-hand corner number indicate the order at which the bars were printed. Scale bar, 5 mm.

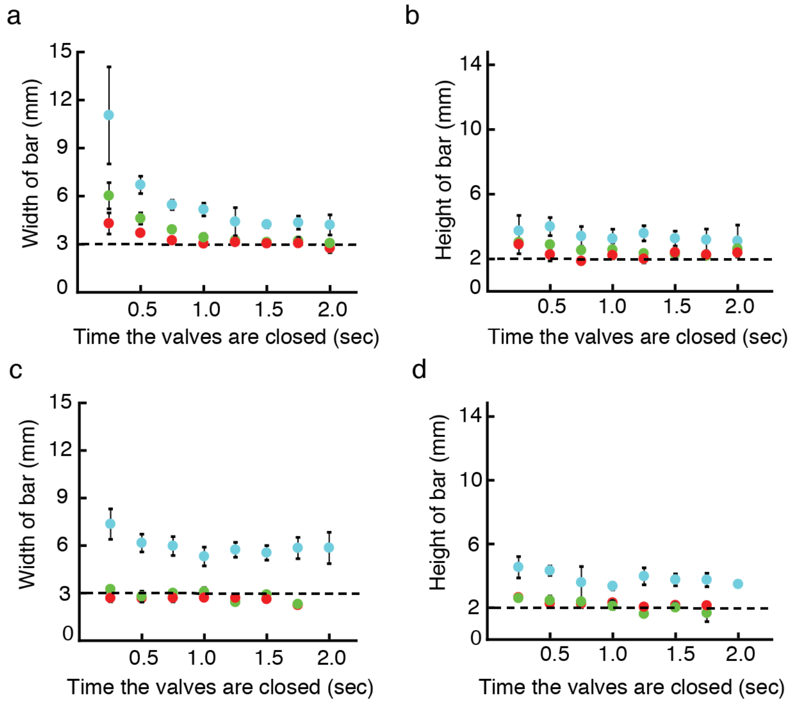

**Figure S21.** **Agarose percentage characterization.** Data are shown without (a and b) and with (c and d) cooling. Data is shown for bars designed to be 2 x 3 x 25 mm (dashed lines). Various % agarose material streams were tested by pulsing the time the solenoid valves are closed (these data were gathered before implementing the relief valve). The agarose percentage has to be between 4-5% for it to work well (3% blue, 4% green, 5% red). The 3% agarose never approaches the expected value. The room temperature was measured to be 22C and the temperature while cooling the chamber between 14 and 17[illegible]C. The datapoints show the mean of eight replicates and the error bars report the standard deviations.

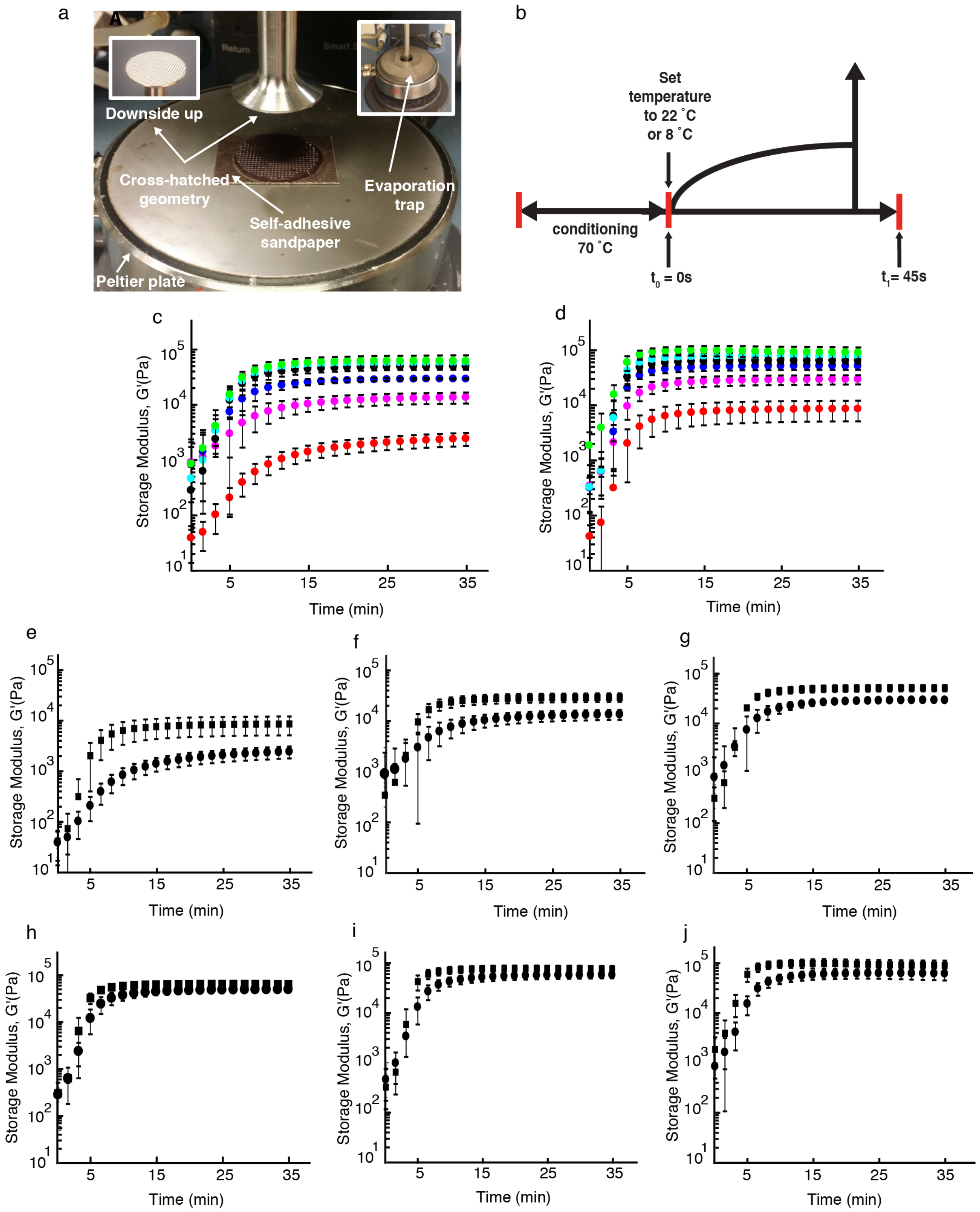

**Figure S22. Gelation time for the different percentage of agarose at 22 and 8°C.** **a**, Rheometer set-up to do this experiment using a Peltier plate. **b**, A schematic showing the conditioning step (temperature at 70°C prior to the start of the data acquisition) and setting the temperature to either 22 or 8°C at the onset of data collection. **c** and **d**, The storage modulus of agarose as a function of time for 22 **(c)** and 8°C **(d)** (red: 1%; magenta: 2%; blue: 3%; black: 4%; cyan: 5%; green: 6%). **e-j**, These plots shows the side to side comparison of the individual percentages at room temperature and at 8°C (e: 1%; f: 2%; g: 3%; h: 4%; i: 5%; j: 6%; squares: 8°C; circles: 22°C). See Embedding Materials Characterization in the Methods section for further details. The error bars represent the standard deviations and these were calculated using three replicates.

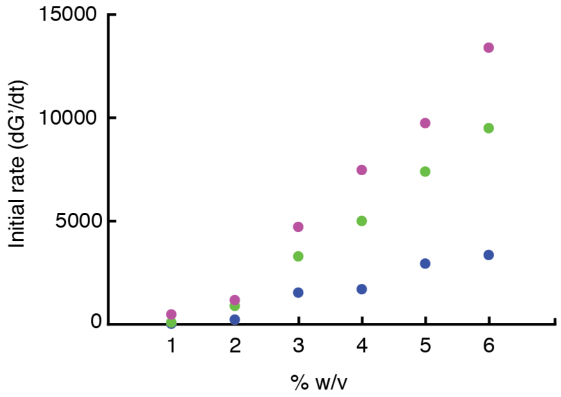

**Figure S23. Initial rate of the gelation for various agarose percentages**. Data are shown for different temperatures: 8°C (magenta), 16°C (green) and 22°C (blue). This shows that the lower the temperature the faster the agarose solidifies. The initial rate was calculated by taking the first derivative of the storage modulus between 0.5 and 5 min. The plot shows the mean using three replicates.

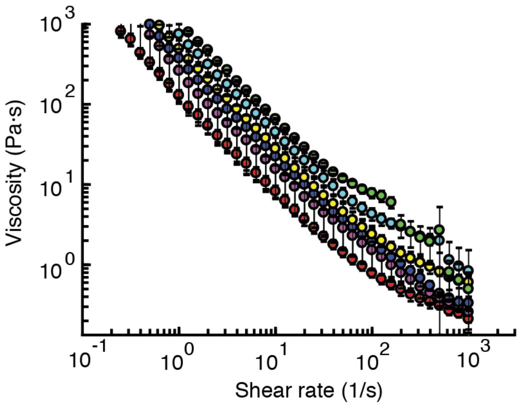

**Figure S24. Shear thinning properties of the Sea Plaque agarose.** The red, magenta, blue, yellow, cyan and green dots represent the agarose at concentration of 1, 2, 3, 4, 5, 6%, respectively. See Methods for experimental details. The error bars represent the standard deviations and these were calculated using three replicates.

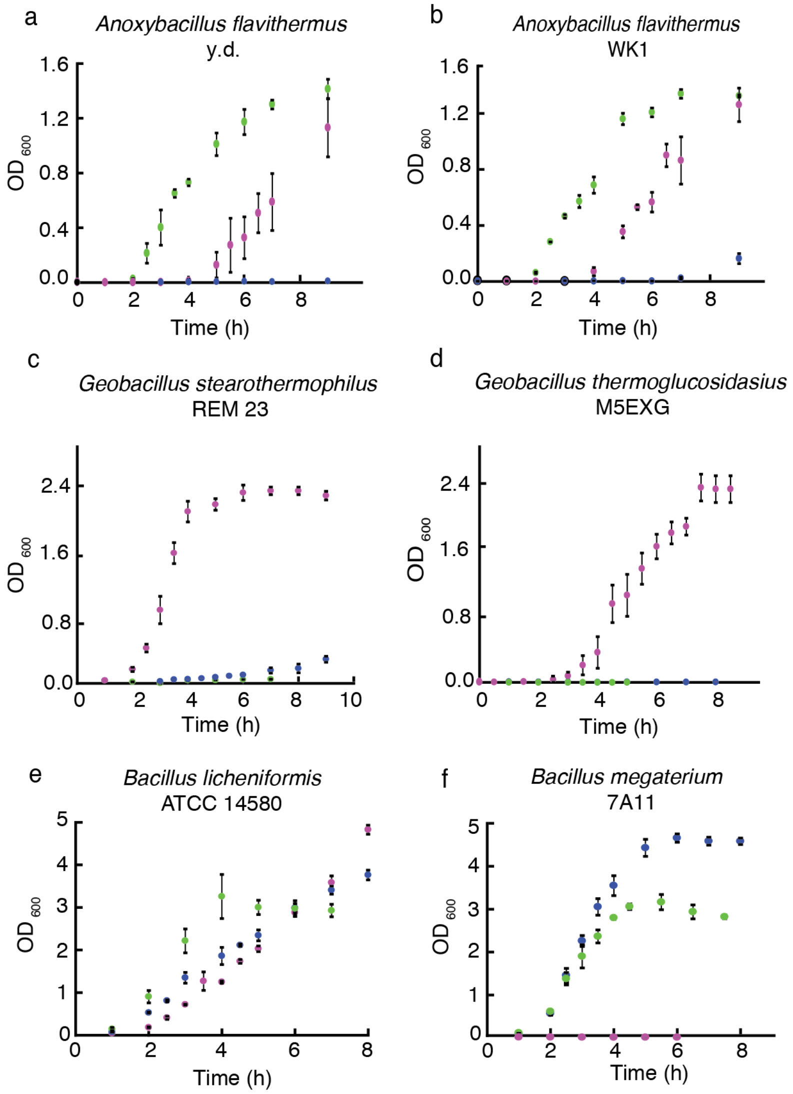

**<u>Figure S25.</u> Comparison between thermophiles and bacilli strains at different temperatures**. **a-f,** Data are shown for the indicated species. The cultures were cultivated at 37°C (blue), 45°C (green) and 55°C (magenta). The error bars represent the standard deviation and these data were calculated from independent experiments collected on three different days, three replicates per day.

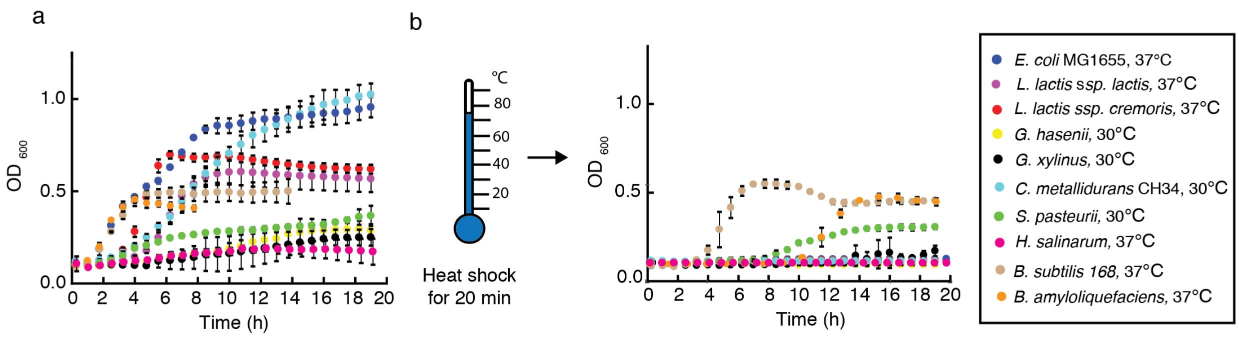

**<u>Figure S26</u>. Screening for species capable of withstanding the high temperatures of the printer**. **a,** Growth curves of a variety of species grown at their optimal temperatures (listed next to the strain name in the legend). **b,** Measurement of growth after heat shock for 20 min at 75°C. In our hands, the cells able to survive this process also are spore-forming species (*B. subtilis*, *S. pasteurii*, *B. amyloliquefaciens*). See Methods for experimental details. The error bars represent the the standard deviation and these data were calculated from independent experiments collected on three different days, four replicates per day.

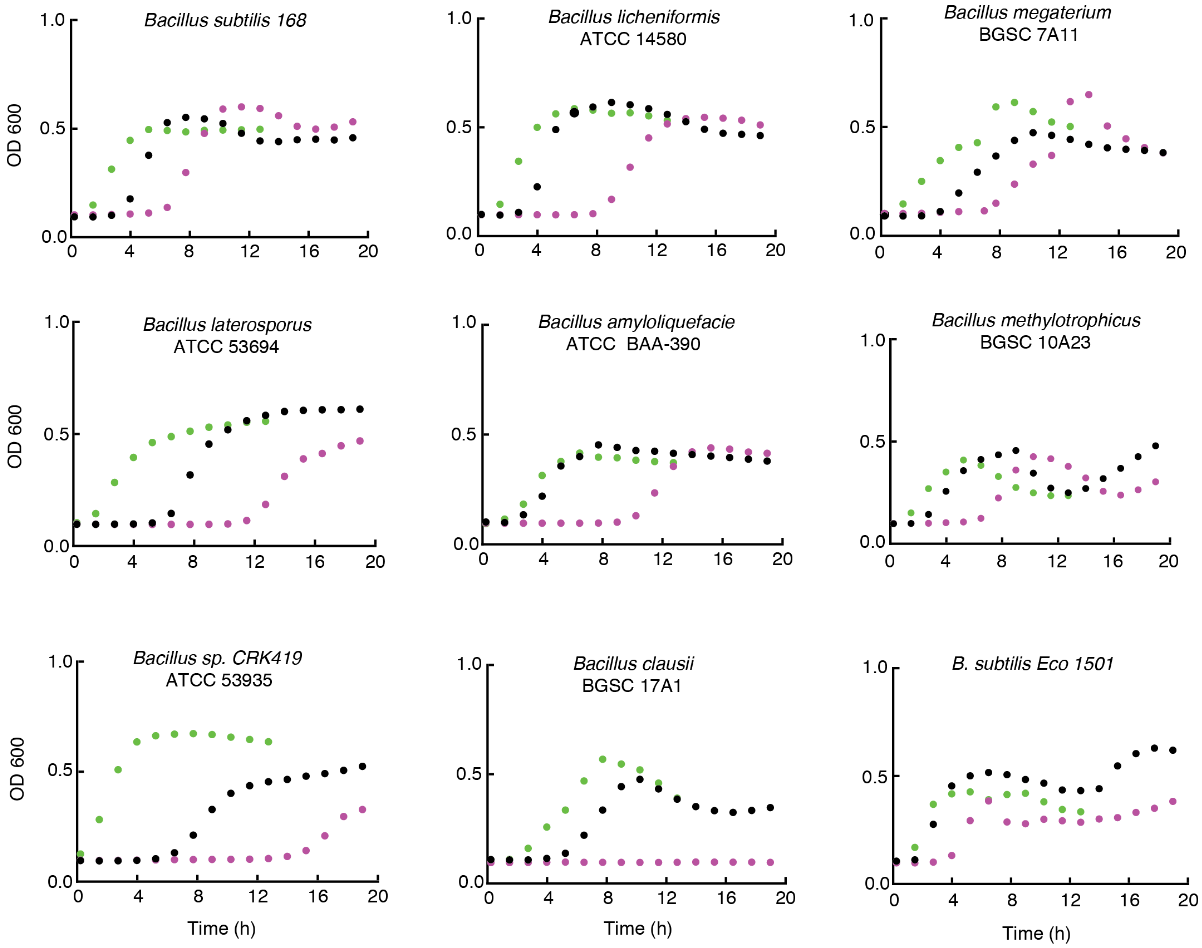

**<u>Figure S27</u>. Comparison of growth curve of different bacilli strains after heat shock**. Cell grown normally without a heat shock (shown in green). Cell grown for two hours in LB media then heat shock at 75°C for 20 min (shown in magenta) and cells induce to sporulate for 2 days in DSM then challenged under a heat shock at 75°C for 20 min (shown in black). The plots show the mean using three replicates performed on the same day.

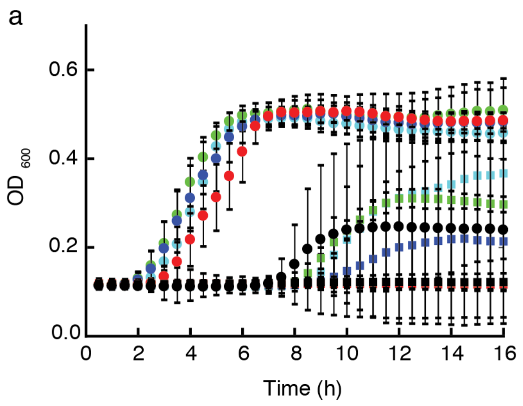

**Figure S28. Exposing spores and vegetative cells to high temperatures.** Cells were exposed to 60°C (cyan), 70°C (green), 80°C (blue), 90°C (red) and 100°C (black) for 20 min using a thermocycler. Circles represent spores and squares represent vegetative cells. For the vegetative group, single colonies were grown for ~3 h in LB media prior to dilutions (Methods). For the spore group, cells were grown in DSM for 2 days prior to dilution. The strain used in this experiment was *B. subtilis* LMG16, containing only a spectinomycin resistance gene. This datapoints are the mean of three experiments performed over different days with four replicates per day.

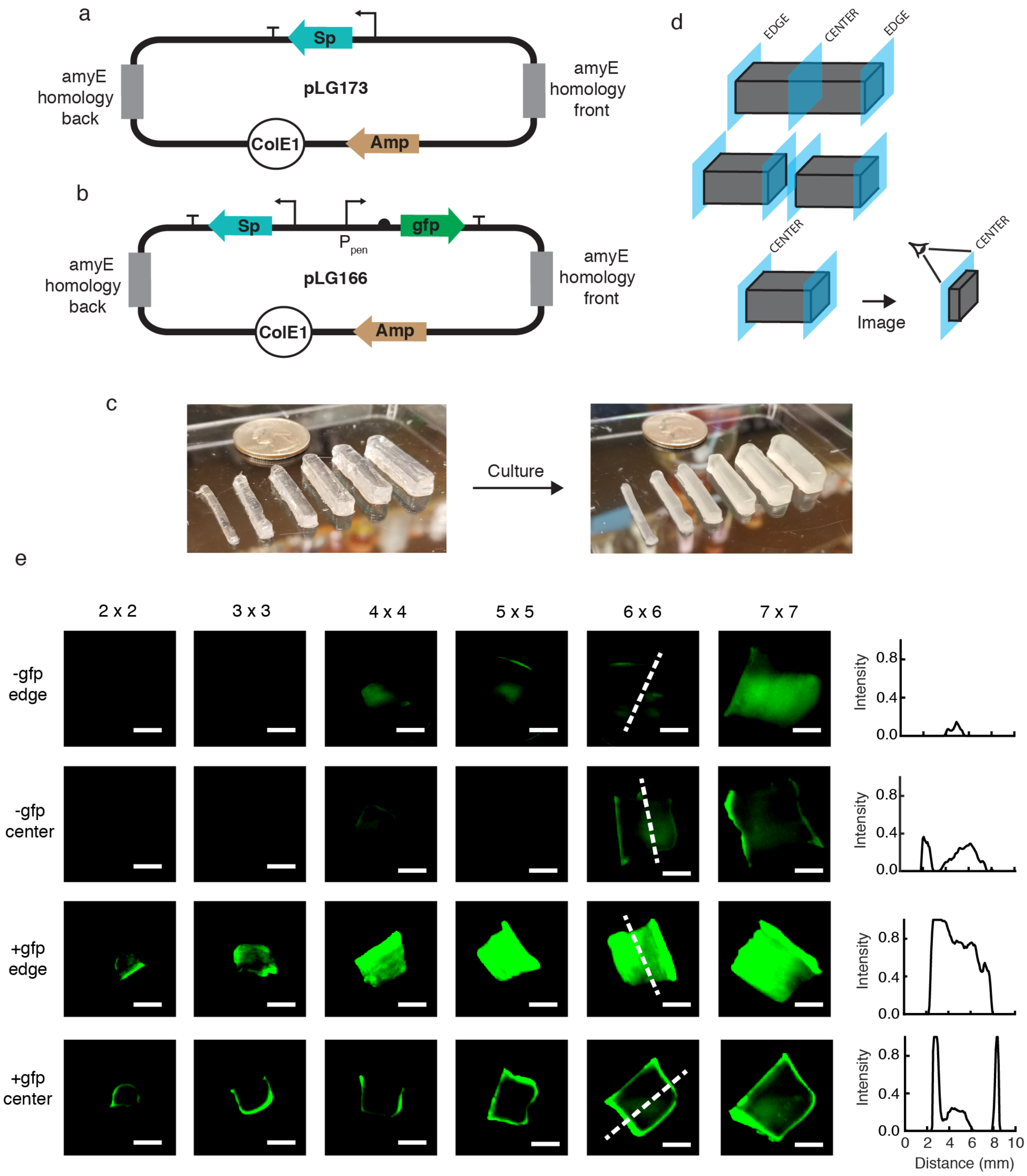

**<u>Figure S29</u>. Growth distribution inside the printed blocks of various sizes**. **a**, The plasmid map for the creation of a *B. subtilis* LMG16 containing only the Sp cassette is shown. **b,** The plasmid map for the *B. subtilis* LMG09 that constitutively expresses GFP is shown. **c,** 3D printed bars containing *B. subtilis* LMG09 before and after culturing in LB media for 10 h. **(d)** Schematic showing how the blocks were sliced. **e,** A thin ~1 mm cross-sectional area was imaged using a Chemidoc Imaging System. +GFP refers to *B. subtilis* LMG09 (constitutive expression) and –GFP refers to *B. subtilis* LMG16 (Sp resistance only). The size of the blocks are a x b x 25 mm where a x b is indicated on the top portion of the image. The graphs to the right of the panel of images indicate the GFP profile across the 6 x 6 x 25 mm blocks. The intensity plots shows the data for a single experiment.

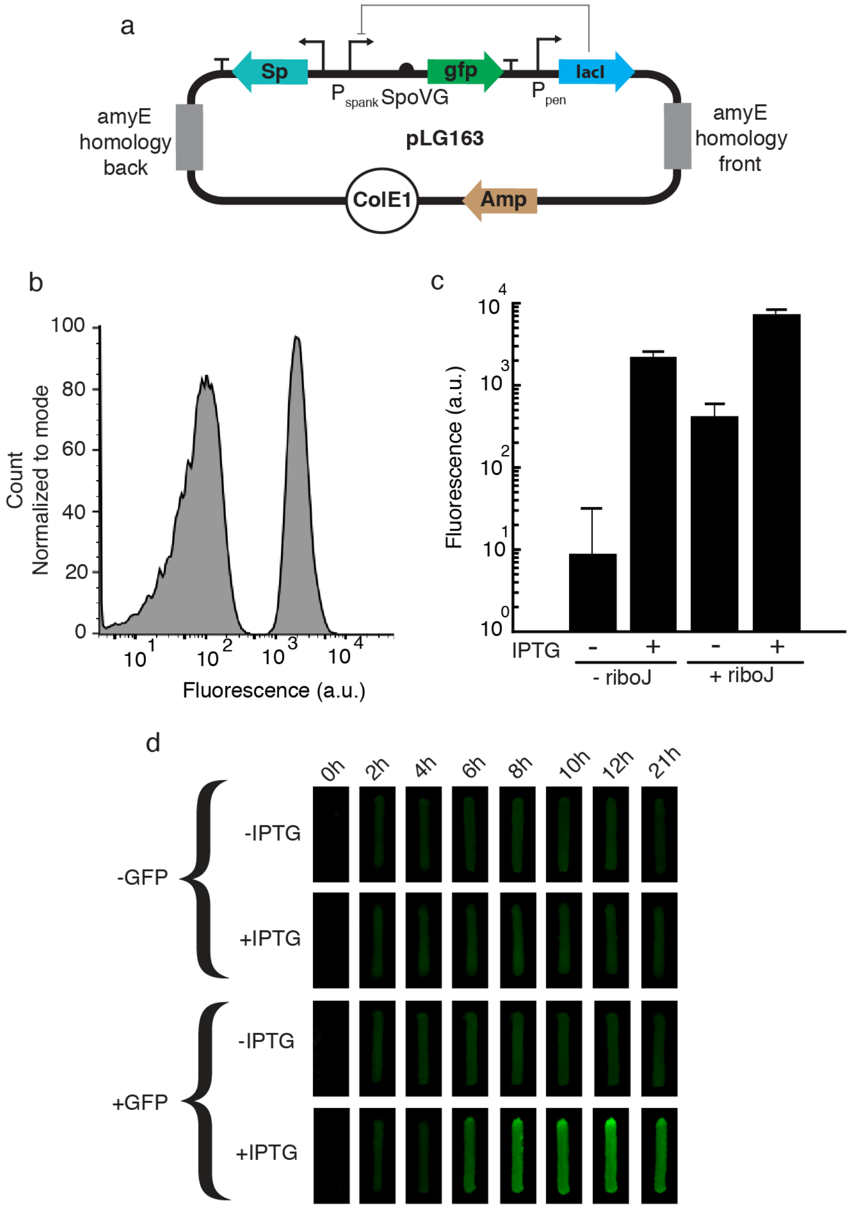

**Figure S30. The IPTG-inducible system in *B. subtilis*.** **a**, Plasmid map, pLG163, with the IPTG sensor used for the construction of the *B. subtilis* LMG04. **b**, Normalized distribution of cells without (left curve) and with 1 mM IPTG (right curve). **c**, Bar graphs of *B. subtilis* LMG04 and LMG59 (made with pLG231) with and without the ribozyme, RiboJ, respectively. There is a 18-fold difference in GFP expression when using riboJ. **d**, Response of embedded cells in agarose to IPTG. Panel of the pictures of all of the time points taken every 2 h and after an overnight growth at 21 h using *B. subtilis* strain LMG04. This panel include the controls (cells harboring the Spectinomycin cassette but no gfp) with and without IPTG. See Viability Assay of Embedded Cells in Methods for further details on data collection for d. The experiment was repeated three times on different days using four replicates and representative images are shown.

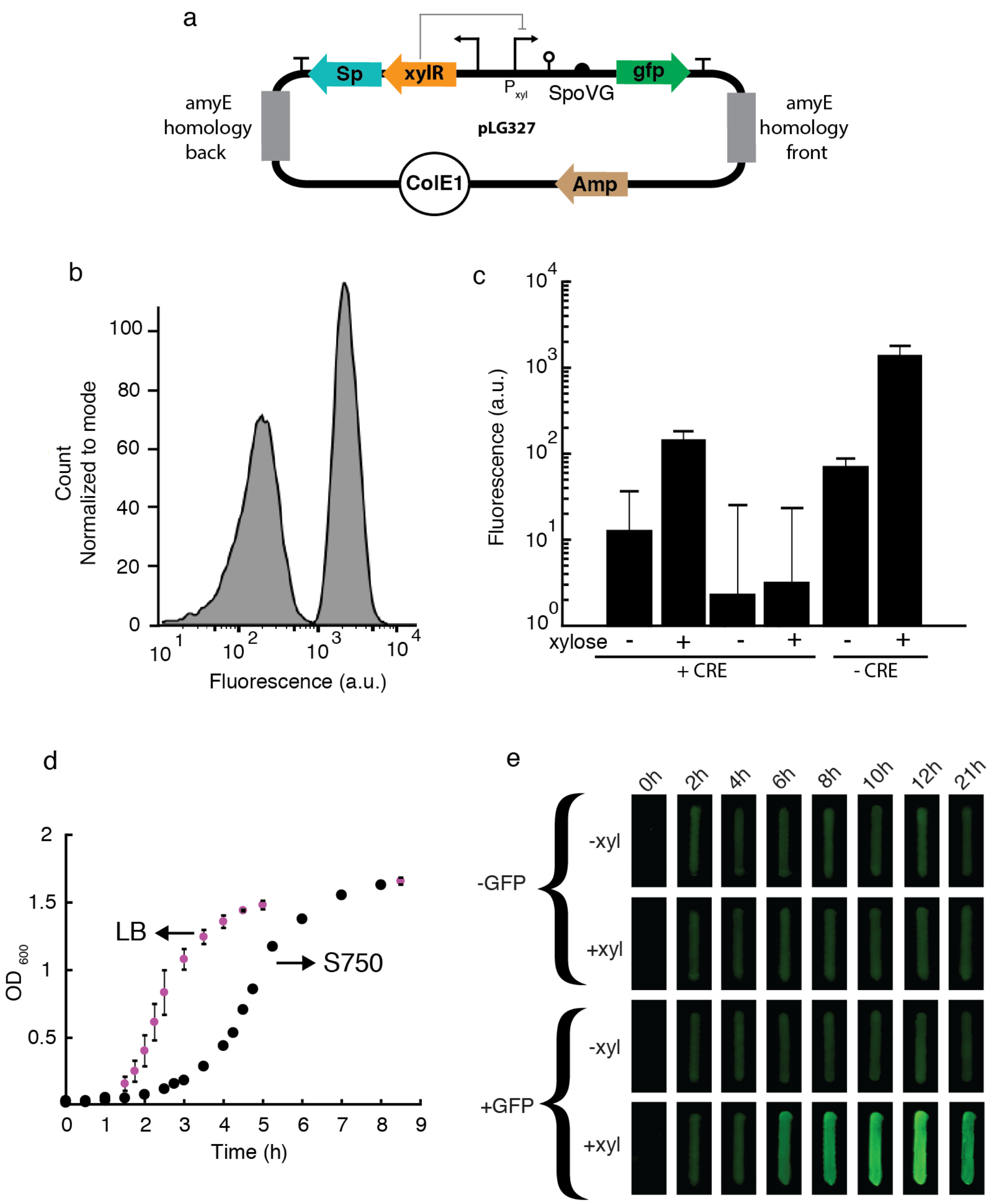

**Figure S31. Improved xylose inducible system in *B. subtilis*.** **a**, Plasmid map, pLG327, with the xylose sensor used for the construction of the *B. subtilis* LMG125. **b**, Normalized distribution of cells without and with 1% xylose. **c**, Bar graphs of *B. subtilis* LMG12 made with pLG169 which harbors the catabolite-responsive element (CRE) in the xylose promoter. *B. subtilis* LMG12 tested in S750 and LB media and comparing the fluorescence activity with and without 1% xylose. The graph also shows the difference in fluorescence activity when the CRE site is removed with and without xylose (*B. subtilis* LMG125). **d**, Growth curve of *B. subtilis* in LB media and the minimal media, S750. **e**, Response of embedded cells in agarose to 1% xylose. Panel of the pictures of all of the time points taken every 2 h, and after an overnight growth at 21 h using *B. subtilis* LMG125. This panel includes the controls (cells harboring the Spectinomycin cassette, but no the gfp one; *B. subtilis* LMG16) with and without xylose. See Methods for details. The experiment was repeated three times on different days using four replicates and representative images are shown.

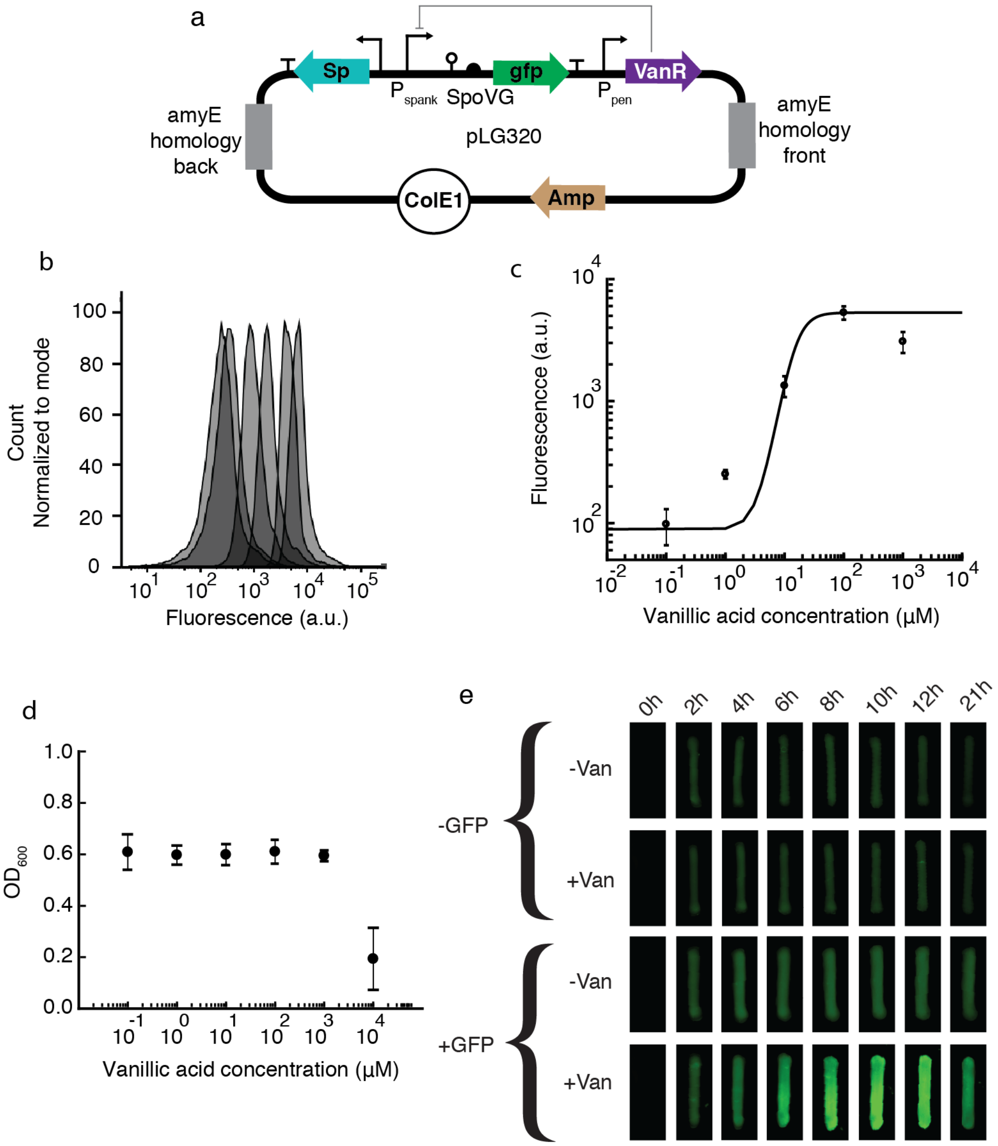

**<u>Figure S32.</u> The vanillic acid inducible system in *B. subtilis*. a,** The plasmid map of pLG320 is shown to build the *B. subtilis* LMG117 with the vanillic acid sensor. **b**, Cytometry data is shown for the induction of *B. subtilis* LMG117 (from left to right: no inducer, 0.1, 1, 10, 100 and 1000µM). **c,** Response function fit to Hill equation, $y_{min} + (y_{max} - y_{min})\frac{x^n}{K^n+x^n}$, where $y_{min} = 88, y_{max} = 5307, K = 15$and $n = 2.9$ The data was plotted in MATLAB using the function lsqcurvefit ( ). **d,** Cell growth is unaffected by concentrations $\leq$ 1mM. **e,** Response of cells in an agarose bar to vanillic acid. Panel of the pictures of all of the time points taken every 2 h, and after an overnight growth at 21 h. This panel includes the controls (cells harboring only a Sp cassette, *B. subtilis* LMG16) with and without 1mM vanillic acid. See Methods for details. The experiment was repeated three times on different days using four replicates and representative images are shown.

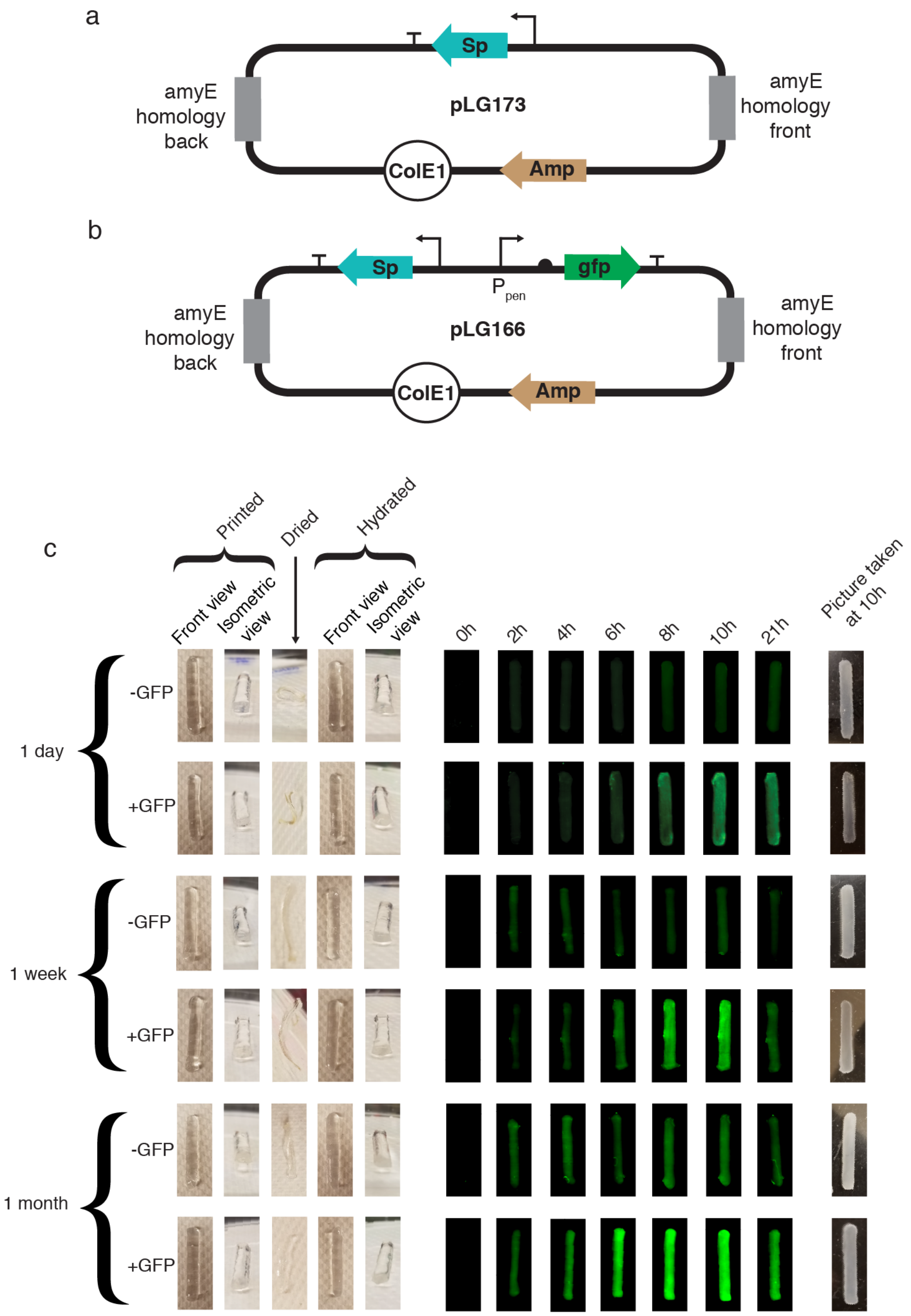

**<u>Figure S33.</u> Exposure of embedded spores to dehydration conditions. a**, The -gfp bars are the ones having cells transformed using pLG173, *B. subtilis* LMG16 (plasmid with only the Sp cassette). **b**, The +gfp bars are the ones having cells transformed with pLG166 (plasmid with the gfp gene expressed constitutively). The transformant cells was name *B. subtilis* LMG09. **c**, Cells were 3D printed, dried overnight and rehydrated after 1 day, 1 week and 1 month. The growth was monitored via fluorescence using gfp and the image were taken with a ChemiDoc Imaging System. A picture was taken at 10h to show the cells emerging from quiescence and show the different between the clear 3D printed part with the dormant spores. See Methods for details. The datapoints are the mean of twelve experiments performed over three days and the error bars are the standard deviation.

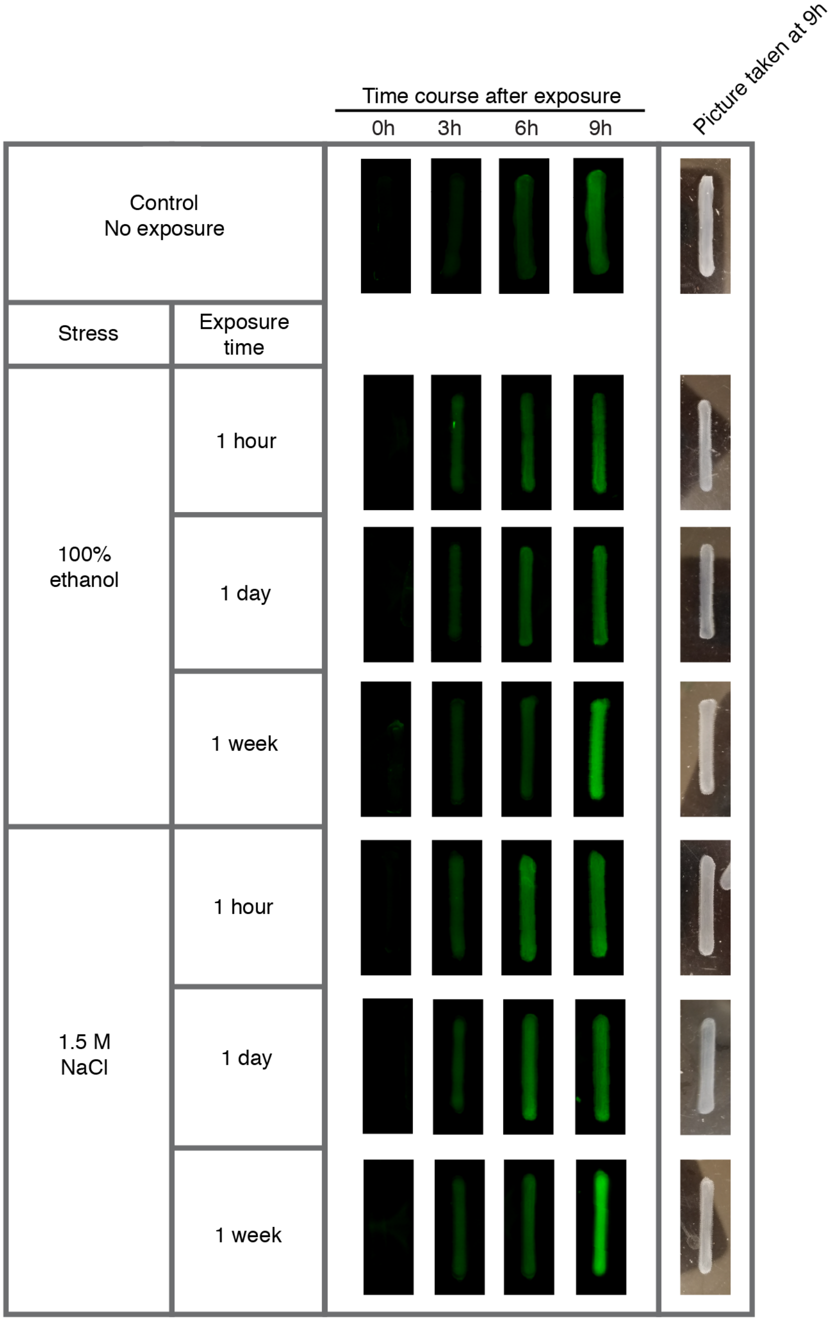

**<u>Figure S34</u>. Challenging spores within the printed agarose bars to ethanol and high osmolarity**. The time course initiates immediately after the exposure period. The strain used for this experiment is *B. subtilis* LMG09. The cell were grown in 5 ml of LB media (with Specinomycin) for 9h. The media was replaced with fresh one every 3 hours. Bars were rinsed with 1X PBS prior to imaging with a ChemiDoc Imaging System. The experiment was repeated three times on different days, with four replicates per day, and representative images are shown.

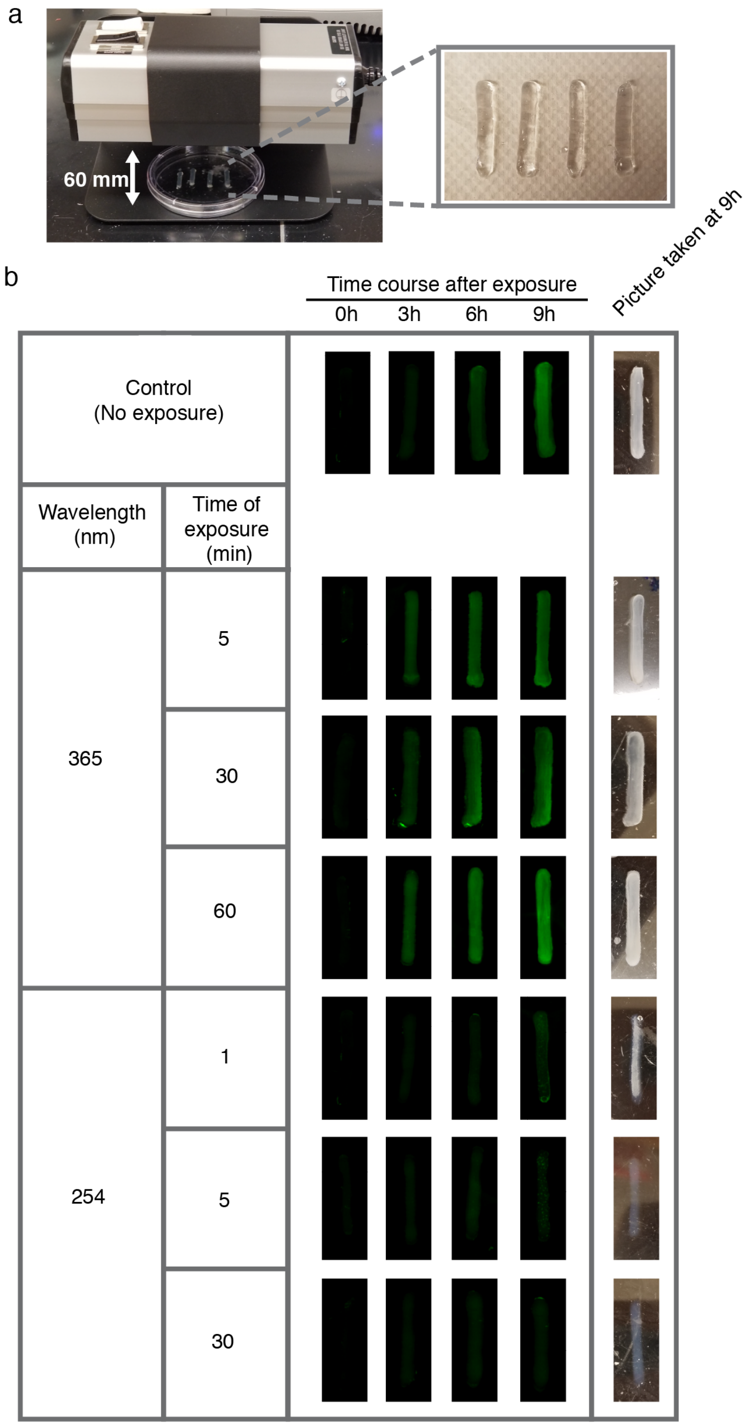

**Figure S35. Challenging spores within the printed agarose bars to Ultraviolet light (UV). a**, Experimental set-up. The UV lamp used in this experiment is shown. The cells were exposed without a petri dish lid and placed in the center (the distance between the lamp and the cells was 6 cm). **b,** The time course was performed immediately following the exposure period. The cells were grown in 5 ml of LB media (with Specinomycin) for 9 h. The media was replaced with fresh media every three hours. The bars were rinsed with 1X PBS prior to imaging with a ChemiDoc Imaging System. Here we show representative bars for each case. The experiment was repeated three times on different days, with four replicates per day, and representative images are shown.

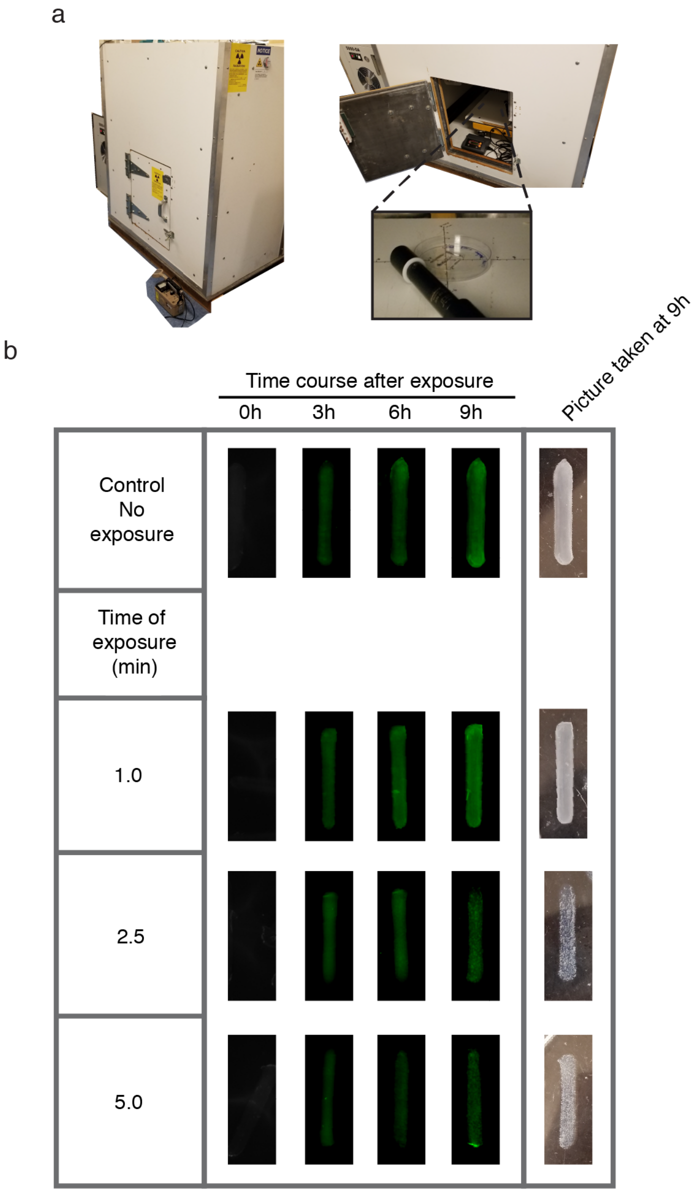

**<u>Figure S36.</u> Challenging spores within the printed agarose bars to X-rays**. **a,** Re-purposed custom X-ray machine used to perform this experiment (Methods). **b,** The measured dose rate was 8.282 R/min. The time course was initiated immediately following the exposure period. The cells were grown in 5 ml of LB media (with Specinomycin) for 9 h. The media was replaced with fresh one every three hours. The bars were rinsed with 1X PBS prior to imaging with a ChemiDoc Imaging System. The experiment was repeated three times on different days, with four replicates per day, and representative images are shown. See Methods for experimental details.

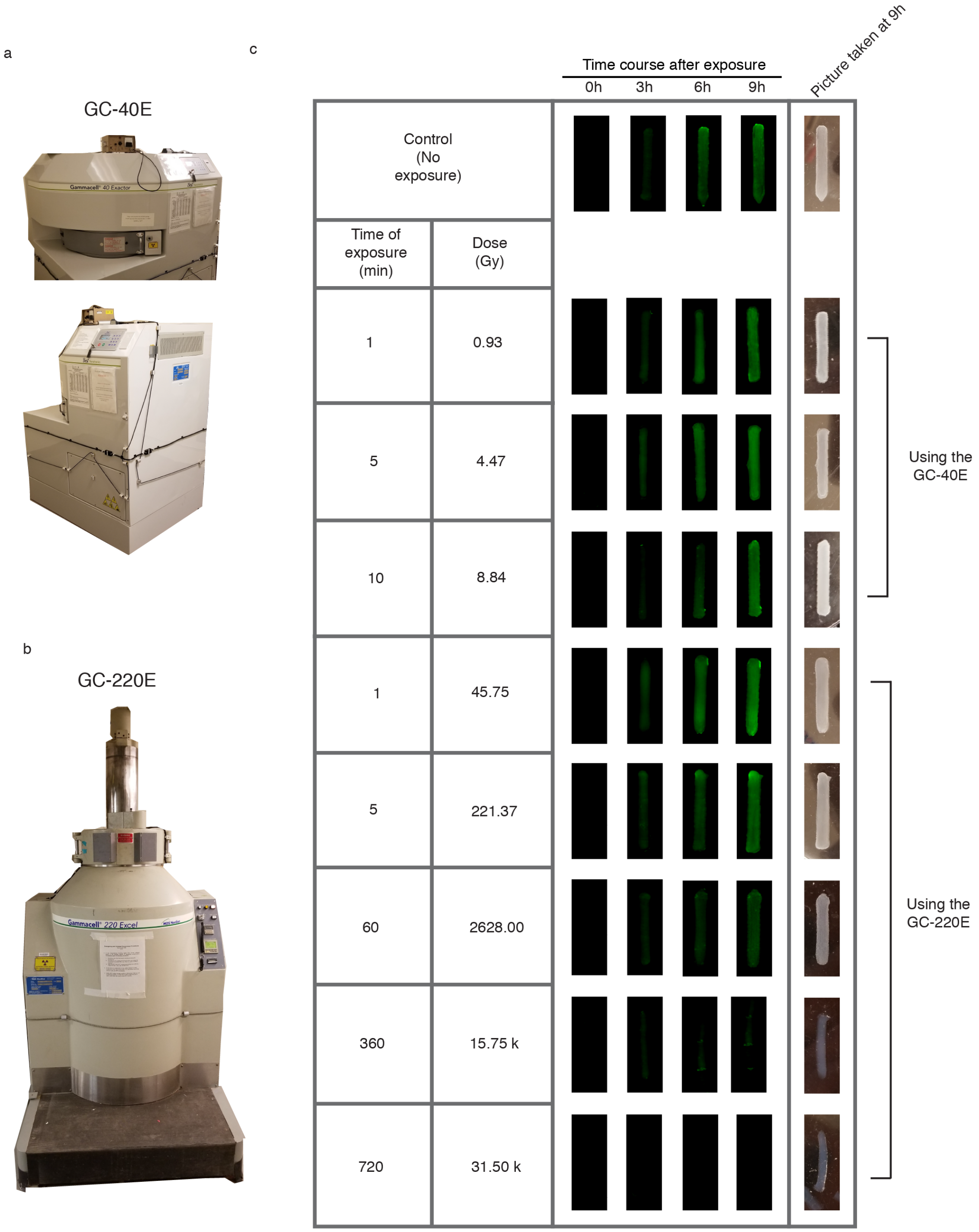

<u>Figure S37</u>. **Challenging spores within the printed agarose bars to γ-radiation**. **a**, Gamma irradiator, GC-40E. **b**, Gamma irradiator, GC-220E. **c**, The dose for this time is note in the second column. The measure dose rate was 93.16 R/min for the GC-40E irradiator. The time course was performed right after exposure. The cells were grown in 5 mL of Lb media (with Specinomycin) for 9h. The media was replaced with fresh one every 3 hours. Bar were rinsed with 1X PBS prior to imaging with a ChemiDoc Imaging System. The experiment was repeated three times on different days, with four replicates per day, and representative images are shown.

### Supplementary Tables

#### Supplementary Table 1: Pin assignment for the 3 Arduino microcontrollers

| **Arduino Mega: to control optical sensors, agarose pumps and cell pumps** | |
|---|---|
| | **Optical sensors** |
| **PIN #** | **Device controlled** |
| 2 | Optical sensor #1 (LEFT extruder) |
| 3 | Optical sensor #2 (LEFT extruder) |
| 18 | Optical sensor #3 (RIGHT extruder) |
| 19 | Optical sensor #4 (RIGHT extruder) |
| 11 | LED#5 indicator for optical sensor#1 (LEFT extruder) |
| 14 | LED#6 indicator for optical sensor#2 (LEFT extruder) |
| 17 | LED#7 indicator for optical sensor#3 (RIGHT extruder) |
| 22 | LED#8 indicator for optical sensor#4 (RIGHT extruder) |
| | **Agarose pump with electronic relief valve** |
| A0 | Thermistor #0 to measure temperature in the printing chamber (ambient) |
| A1 | Electronic relief valve analog pin |
| A2 | Thermistor #5 to measure temperature at the top of the mixing chamber LEFT printhead |
| A3 | Thermistor #6 to measure temperature at the top of the mixing chamber RIGHT printhead |
| 4 | Electronic relief valve digital pin |
| 5 | Solenoid valve #1 controls dispensing of agarose through LEFT extruder |
| 6 | Solenoid valve #2 controls dispensing of agarose through RIGHT extruder |
| 9 | LED#1 indicates when solenoid valve #1 is on |
| 10 | LED#2 indicates when solenoid valve #2 is on |
| | **Cell pumps** |
| 20 | LEFT optical sensor#5 to trigger cell liquid pump |
| 21 | RIGHT optical sensor #6 to trigger cell liquid pump |
| 12 | Liquid pump#1 to pump cells |
| 8 | Liquid pump#2 to pump cells |
| 15 | Solenoid valve #4 for liquid pump #1 |
| 27 | Solenoid valve #5 for liquid pump #2 |
| 16 | LED#4 indicates when either cell liquid pump #1 is used |
| 26 | LED#3 indicates when either cell liquid pump #2 is used |
| **Arduino Uno: to control scaffolding using gelatin** | |
| A0 | Optical IR distance sensor (detect platform z position) |
| 1 | LED#9 indicates when optical IR distance sensor is used |
| 2 | Liquid pump#4 to pump gelatin |
| 3 | Solenoid valve #6 for gelatin liquid pump |
| 4 | Liquid pump#3 to pump gelatin for cooling the platform |
| 5 | LED#10 indicates when cooling is activated |
| 6 | FAN for cooling the printing chamber |
| **Arduino Uno (RFIDUINO) : temperature control** | |
| VREF | REF pin |
| 9 | PWM pin to control the temperature in the chamber through remote power supply |
| A1 | Thermistor #1 to measure temperature in the bottom of the mixing chamber (LEFT printhead) |
| A2 | Thermistor #2 to measure temperature in the middle of the mixing chamber (LEFT printhead) |
| A0 | Thermistor #3 to measure temperature in the middle of the mixing chamber (RIGHT printhead) |
| A3 | Thermistor #4 to measure temperature in the middle of the mixing chamber (RIGHT printhead) |
| --- | Liquid-crystal display (LCD) to monitor temperature in mixing chamber |
| --- | RFID shield to turn on system |

#### Supplementary Table 2: Parts to Build 3D Printer

| Parts | Function | Part No. | Distributer |
|---|---|---|---|
| MarkerBot Replicator 2 |  | ---- | MarkerBot |
| Desiccator cabinet | Agarose pump | Model # 33060 | Fisher Scientific |
| Pressure gauge | Agarose pump | 4000K721 | McMaster-Carr |
| Magnet wire, enameled, AWG 21 | Heating lines in agarose pump | AWG21-200C-11 | Applied Magnets |
| High temperature silicone rubber tubing (OD = ¾", ID = ½") | Agarose pump | 5236K47 | McMaster-Carr |
| High temperature silicone rubber tubing (OD = 5/16", ID = 3/16") | Agarose pump | 5236K841 | McMaster-Carr |
| Straight reducer barbed fitting | Agarose pump | 5463K635 | McMaster-Carr |
| Push-in signal power connector | Agarose pump<br>(allow for easier cleaning of the lines) | 9193T11 | McMaster-Carr |
| High temperature silicone rubber tubing (OD=1/4", ID=1/8") | Agarose pump | 5236K83 | McMaster-Carr |
| Electronic Relief valve (ERV) | Controls agarose flowrate | ERV-1012-0505 | Kelly Pneumatics |
| Silicon heat tape,<br>½" x 24" | Agarose pump (maintain agarose in a molten state inside the chamber) | MFR #: HSTAT051002 | Zero |
| Solenoid valve (5) | Agarose pump (2)<br>DC liquid pump (3) | Item #:<br>HW-WVALVE | Trossen Robotics |
| DC liquid pump (4) | Gelatin pump (1)<br>Cell pumps (2)<br>Water pump (1) | Item # TI-TG-02B-DC12B | Trossen Robotics |
| One-way valves (9) | Cell pumps (8)<br>Agarose pump (1) | | |
| Peltier plates + heat sink assembly (4) | Cooling system | Product ID: 1335 | Adafruit |
| Female DC power adaptor (2.1 mm jack to screw terminal block) | Cooling system | Product ID: 368 | Adafruit |
| 12V 5A Switching power supply | Cooling system | Product ID: 352 | Adafruit |
| Fan (4.69" x 4.69 x 1.5", airflow =205 cfm) | Cooling system | 1939K96 | McMaster-Carr |
| Fan Guards<br>(safety feature) | Cooling system | 19155K125 | McMaster-Carr |
| Bench Power supply- 1901B<br>30A, 32V | Heating lines with constant current<br>(output max current =<br>30 A) | 78Y9404 | Newark |
| Bench Power supply- 1964<br>(30A, 30V) | Heating printhead<br>Using remote control features | 34T4663 | Newark |
| Power supplies<br>3-12V, 2A (6) | To power pump and solenoids | Cat no. 29902PS | Marlin P. Jones & Assoc. Inc. |
| Mini Incubator | Gelatin subsystem<br>(houses the gelatin pump and gelatin itself) | Cat no. LB-BO-MINI-15110A | Lab Planet |
| IR Distance sensor | Gelatin subsystem (to detect the position of the platform | Product ID:<br>GP2Y0A21YK0F | Adafruit |
| Robot Geek Large Workbench | Electronics support board | Item #: ASM-WRKBL | Trossen Robotics |
| Arduino Uno microcontroller (2) | Electronics controller | Part number: 2151486 | Jameco Electronics |
| Arduino Mega microcontroller | Electronics controller | Part No. 1050-1018-ND | Digi-Key |
| RobotGeek Relay board (4) | Use to control the DC liquid pumps | Item #: ASM-RG-Relay | Trossen Robotics |
| Robot Geek Duino Mount (3) | To Mounts Arduino Uno and Arduino Mega | Item #: ASM-RG-DUINOMNT | Trossen Robotics |
| Robot Geek Sensor Shield | Allows to connect to microcontroller without breadboard; it has 3 pin connectors | Item #: RG-SENSHIELSV2 | RobotGeek |

| | | | |
|---|---|---|---|
| TinkerKit Mega Sensor Shield | Allows to connect to microcontroller without breadboard; it has 3 pin connectors | SKU: T020040 | Core Electronics |
| RobotGeek LCD Display with mount | To display the temperature inside Printhead and chamber | Item #: ASM-RG-LCD | Trossen Robotics |
| RFIDuino Shield | To turn on apparatus | Item #: RG-RFIDUINO | Trossen Robotics |
| RFID Key Fob | To turn on apparatus | Item #: RFID-TAG-125-KCBLUE | Trossen Robotics |
| Darlington Transistor<br>NPN Power | Use to control the solenoid valves | Part No. TIP102<br>Mouser Part No: 511-TIP102 | Mouser Electronics |
| Optical sensor (2)<br>MPN: EE-SX1070 | Detects moving motor and when motor reverses direction | Part no. OR571-ND | Digi-Key |
| Optical Sensor (1)<br>MPN: EE-SX1042 | Detect when to input cells/spores | Cat. No. OR518-ND | Digi-Key |
| Thermistor (d) (4) | To measure the temperature in the system<br>Printhead (3)<br>Printing cabinet (1) | Product ID: 372 | Adafruit |
| Capacitors<br>(50V, 10μF)<br>(0.1 μF) | Temperature control and IR sensor | | Digi-Key |
| Diode (general purpose) | Solenoid control with power transistor (TIP102) | 1N4004-TPMSTR-ND | Digi-Key |
| Resistors:<br>220 Ω<br>470 Ω (6)<br>1 kΩ (5)<br>1.5 kΩ (1)<br>10 kΩ (7) | 470 Ω (pull-up resistors for LEDs)<br>1 kΩ (for solenoid control)<br>1.5 kΩ (remote control power supply)<br>10 kΩ (for thermistor control)<br>10 kΩ (for optical sensor control | | Digi-Key |
| Light Emitting Diodes (LEDs) | To indicate what components are active | --- | Digi-Key |
| A male barbed tube fitting tee connector | Agarose line for dual printing<br>Tube ID = 3/16”<br>Nylon Plastic<br>Temperature range = -40 to 170°F | 5463K183 | McMaster-Carr |

#### Supplementary Table 3: Printed Parts Using 3D printer ProJet 6000 (3D Systems)

| Name of part | Function | Picture of the parts | File Names |
|---|---|---|---|
| Mixing chamber | Mixes the agarose and cells/spores before printing | 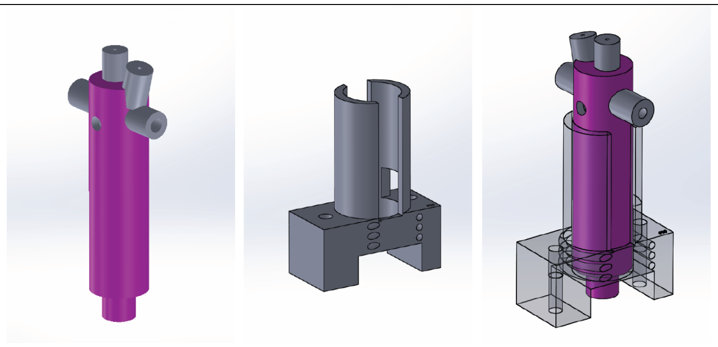 | •Prismatic_slider_side_nozzle_num51_1_17_18_slanted_nozzle. STL<br><br>•Prismatic_slider_holder_num29_7_3_17.STL |
| MS Holder | Holder for motor (s) solenoid valve (s) | 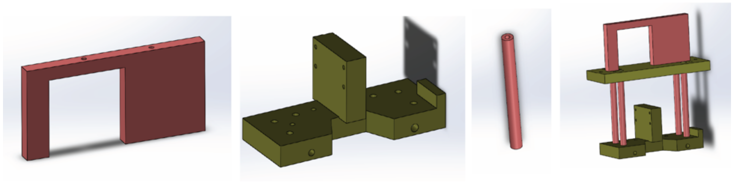 | •Lower_platform_duo_printhead_num3_01_15_18.STL<br>•Upper_platform_num4_11_20_15.STL<br>•Motor_holder_num5_01_25_16.STL<br>•Platform_legs_num3_2_25_17.STL |
| OS Holder (1) | Holder for optical sensors (dual printing and detecting change of direction) | 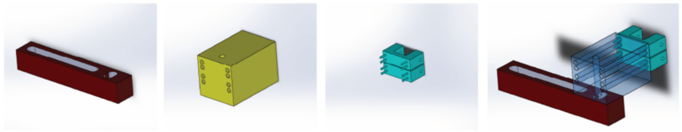 | •Slot_sensor_holder_vertical.STL<br><br>•Slot_sensor_holder_horizontal.STL |
| OS Holder (2) | Holder for optical sensors (cell input) | 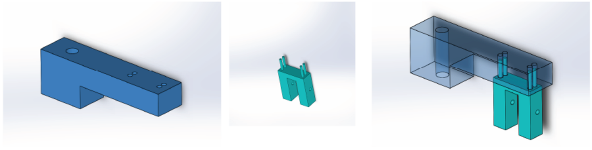 | •Slot_sensor_movement_top_num4_11_04_18.STL<br><br>•Slot_sensor_movement_top_num4_11_04_18.STL |
| Agarose pump connector | Connector to facilitate transfer from agarose pump to printhead | 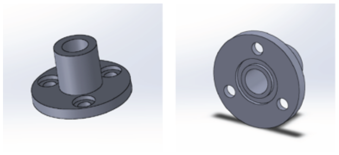 | •Connector for pressurized tank.STL |
| Printer cover | Cover to aid cooling the chamber | 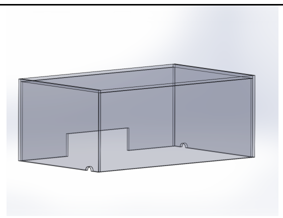 | •cover_side_1.SLDPRT<br>•cover_side_2.SLDPRT<br>•cover_side_3.SLDPRT<br>•cover_top.SLDPRT |

#### Supplementary Table 4: Relief Valve and Pressure Relationship

| Pressure (psi) | Range (analog read) |
|---|---|
| 1 | 150-200 |
| 1.5 | 200-250 |
| 2 | 250-300 |
| 2.5 | 300-350 |
| 3 | 350-400 |

#### Supplementary Table 5: List of Strains Used

| Organism name | ATCC #, BGSC# or DSM# or source |
|---|---|
| *Anoxybacillus flavithermus WK1* | DSM 21510 |
| *Anoxybacillus flavithermus d.y.* | DSM 2641 |
| *Bacillus subtilis* 168 | ATCC 23857 |
| *Bacillus subtilis* PY79 | |
| *Bacillus amyloliquefaciens* | ATCC BAA-390 |
| *Bacillus clausii* | BGSC 17A1 |
| *Bacillus* sp. CRK419 | ATCC 53935 |
| *Bacillus subtilis Eco1501* | Ecovative |
| *Bacillus laterosporus* | ATCC 53694 |
| *Bacillus licheniformis* | ATCC 14580 |
| *Bacillus megaterium* | BGSC 7A16 |
| *Bacillus methylotrophicus* | BGSC 10A23 |
| *Cupriavidus metallidurans* CH34 | ATCC 43123 |
| *Escherichia coli* | MG1655 |
| *Geobacillus stearothermophilus* REM23 | REM 23 |
| *Geobacillus thermoglucosidasius* M5EXG | ATCC BAA-1069 |
| *Halobacterium salinarum strain* NRC-1 | ATCC 700922 |
| *Lactococcus lactis* subsp. cremoris MG1363 | Mobitec |
| *Lactococcus lactis* subsp. lactis | ATCC 11454 |
| *Gluconacetobacter xylinus* | ATCC 700178 |
| *Gluconacetobacter haseniii* | ATCC 23769 |
| *Sporosarcina pasteurii* | ATCC 11859 |

#### Supplementary Table 6: List of Plasmids and Engineered *Bacillus* Strains

| Name | Figure where construct shown | Parent strain name (*Bacillus subtilis* PY79) | Function |
|---|---|---|---|
| pLG163 | Figure 4, Supplementary Fig. 30 | LMG04 | IPTG-inducible system |
| pLG166 | Figure 1, 2, 3, 5, Supplementary Fig. 29, 34-38 | LMG09 | Constitutive GFP expression |
| pLG169 | Supplementary Fig. 32 | LMG12 | Xylose-inducible system *with* CRE site |
| pLG173 | Figure 4, Supplementary Fig. 29, 30-34 | LMG16 | Only the Specinomycin marker (control) |
| pLG213 | Figure 4, Supplementary Fig. 31 | LMG44 | aTc-inducible system |
| pLG231 | Supplementary Fig. 30 | LMG59 | IPTG inducible system |
| pLG295 | Figure 6 | LMG129 | Constitutive melanin expression |
| pLG320 | Figure 4, Supplementary Fig. 33 | LMG117 | Vanillic acid-inducible system |
| pLG327 | Figure 4, Supplementary Fig. 32 | LMG125 | Xylose inducible system *without* CRE site |

**Note**: All of the plasmids listed here were integrated into the amyE locus.

#### Supplementary Table 7: List of Genetic Parts (not to be confused with the mechanical parts)

| Part Name | Type | DNA sequences | Ref. |
|---|---|---|---|
| $P_{spank(hy)}$ | Promoter | **AAATGTGAGCACTCACAATT**CATTTTGCAAAAGTTGTTGACTTTATCTACAAGGTGTGGC<br>ATAATGTGTGTA**AATTGTGAGCGGATAACAATT**AAGCTTTCGGCTG | 1 |
| $P_{spank(V)}$ | Promoter | GGCAAGAACGTTGCTCGAG**ATTGGATCCAAT**CATTTTGCAAAAGTTGTTGACTTTATCTA<br>CAAGGTGTGGCATA**TTGGATCCAAT**AGC | This work |
| $P_{xyl}$ | Promoter | CCTTTATTATATCTAATGTGTTCATGAAAAACTAAAAAAAATATTGAAAATACTGATGAG<br>GTTATTTAAGATTAAAATAA**GTTAGTTTGTTTGGGCAACAAACTAAT**GTGCAACTTACT<br>TACAATATGACATAAAATGCATCTGTATTTGAATTTATTTTAAGGAGGAAATAACATGG<br>CTCAATCTCATTCTAGTTCAGTTAACTATTTTGTAAGCGTTAACAAAGTGGTTTAATTAAG<br>TCGACAGCTAGCCGCACAATTTTATGTAAGG | 2 |
| $P_{xyl(S)}$ | Promoter | ATATCTAATGTGTTCATGAAAAACTAAAAAAAATATTGAAAATACTGATGAGGTTATTTA<br>AGATTAAAATAA**GTTAGTTTGTTTGGGCAACAAACTAAT**GTGCAACTTACTTACAATAT<br>TTACTGTTTGAGTCGGCTG | This work |
| $P_{tetR2}$ | Promoter | GGATCACCGGTACCTTGACATAACTG**TTTATCAGTGATAAA**GTCTGTTGCAGATCTTATT<br>ATTCACACTTTCTAGAAATAATTTTCTTAAGTGTTTGAGTCGGCTG | 3 |
| $P_{ptsG}$ | Promoter | TTAATGTTGTTATTGAAAAATGAATATCCGCTATGCTACAATACAGCTTGGAAATTGATTA<br>AAATCTTGCAGCAAACATTGAAGAAACCAGTTCATGAGGCGGAAGCGGTTTATCTGACGC<br>TTCATCTGTACCGATTAACCAATAAAATTTCATAAATTCAGTTTATCCTTATAACGTGTTA<br>CTGATTCGATCAGGCATGAGTGATTGAGGGAAAAAAACGGGAAGTTCATTCTCGTTCTTT<br>TGCGCACCCAATTTTGCTCATGCCTTTTTGTTGTGTAAAAGGGCAAATGTAAACGGTTAAA<br>CTGGAAGACTTACGCCTGTGAATTCGTTGTCATGATTTTTAGCTGTAAGGTCAGACTAGTA<br>AAAAGAGGAGGTCAATT | 3 |
| $P_{pen}$ | Promoter | CGGTGGAAACGAGGTCATCATTTCCTTCCGAAAAAACGGTTGCATTTAAATCTTACATAT<br>GTAATACTTTCAAAGACTACATTTGTAAGATTTGA | 4 |
| riboJ | Insulator | AGCTGTCACCGGATGTGCTTTCCGGTCTGATGAGTCCGTGAGGACGAAACAGCCTCTACA<br>AATAATTTTGTTTAA | 5 |
| spoVG | RBS | AAAGGTGGTGAA | 6 |
| GsiB | RBS | TAAAGGAGGAA | |
| CRE site | Operator | TTGTAAGCGTTAACA | 7 |
| *vanR*$^{AM}$ | Repressor | ATGGACATGCCTCGTATTAAACCGGGTCAGCGTGTTATGATGGCACTGCGTAAAATGATT<br>GCAAGCGGTGAAATCAAAAGTGGTGAACGTATTGCAGAAATTCCGACCGCAGCAGCACT<br>GGGTGTTAGCCGTATGCCGGTTCGTATCGCACTGCGTTCACTGGAACAAGAAGGTCTGGT<br>TGTTCGTCTGGGTGCACGTGGTTATGCAGCCCGTGGTGTTAGCAGCGATCAGATTCGTGAT<br>GCAATTGAAGTTCGTGGTGTTCTGGAAGGTTTTGCAGCACGTCGTCTGGCAGAACGTGGT<br>ATGACCGCAGAAACCCATGCACGTTTTGTTGTACTGATTGCAGAAGGTGAAGCACTGTTT<br>GCAGCCGGTCGCCTGAATGGTGAAGATCTGGATCGTTATGCCGCATATAATCAGGCATTT<br>CATGATACCCTGGTTAGCGCAGCAGGTAATGGTGCAGTTGAAAGCGCACTGGCACGTAAT<br>GGTTTTGAACCGTTTGCAGCAGCCGGTGCACTGGCCCTGGATCTGATGGACCTGTCTGCC<br>GAATATGAACATCTGCTGGCAGCACATCGTCAGCATCAGGCAGTTCTGGATGCAGTTAGC<br>TGTGGTGATGCCGAAGGTGCAGAACGTATTATGCGTGATCATGCACTGGCAGCAATTCGT<br>AATGCAAAAGTTTTTGAAGCAGCAGCAAGCGCAGGCGCACCGCTGGGTGCAGCATGGTC<br>AATTCGTGCAGATTGATAA | 8, 9 |
| *xylR* | Repressor | ATGACTGGATTAAATAAATCAACTGTTTCATCACAGGTAAACACGCTGATGAAAGAAAAT<br>CTTGTATTTGAAATAGGTCAAGGACAATCAAGTGGCGGAAGAAGACCTGTGATGCTTGTT<br>TTTAATAAGAAGGCAGGATACTCCATAGGCATAGATGTTGGTGTGGATTATATTAGTGGG<br>ATTTTAACAGACCTTGAAGGGACGATCATTCTTGATCAGCAAAGTGGCGGAAGAAGACCT<br>GTCATGCTTGTTTTTAATAAAAAGGCAGGATACTCCATAGGAATAGATGTTGGTGTGGAT<br>TATATTAGTGGCATTTTAACAGACCTTGAAGGAACGATCATTCTTGATCAACATCACCATT<br>TAGAATCGAATTCTCCAGAAATAACTAAAGACATTTTAATTGATATGATTCATCACTTTAT<br>TACGCGTATGCCACAATCTCCGTACGGGCTTATCGGTATAGGCATATGCGTGCCTGGACT | 2 |

| | | | |
|---|---|---|---|
| | | AATTGATAAAAATCAAAAAATTGTTTTCACTCCGAACTCCAACTGGAGAGATATTGACTT<br>AAAATCTTTCATACAAGAGAAGTTCAATGTGCCTGTTTTTATTGAAAATGAGGCAAATGC<br>TGGCGCGTATGGAGAAAAAGTATTTGGTGCTGCAAAAAATCACAATAACATTATTTATGC<br>TAGTATCAGTACAGGTATAGGGATCGGTGTTATTATCAACAATCATTTATATAGAGGGGT<br>AAGCGGATTCTCTGGAGAAATGGGACATATGACAATAGACTTTAATGGTCCTAAATGCAG<br>TTGCGGAAACCGAGGTTGCTGGGAATTGTATGCTTCAGAGAAGGCTTTATTAAAATCTCT<br>TCAGACTAAAGAGAAAAAAGTGTCCTATCAAGATATCATAGACCTCGCCCATCTGAATGA<br>TATCGGCACCTTAAATGCATTACAGAATTTCGGATTCTATTTAGGAATTGGCCTTACTAAT<br>ATTCTAAATACATTCAATCCACAAGCCATCATTTTAAGAAACAGCATAATTGAATCACAT<br>CCTATGGTTTTAAATTCAATTAGAAGTGAAGTGTCGTCCAGGGTTTATCCCCAATTAGGCA<br>ATAGCTATGAATTATTACCATCTTCCTTAGGAAAGAATGCACCGGCATTAGGAATGTCTTC<br>CATTGTTATTGAACATTTTCTAGATATCGTTAAAATGTAA | |
| *tetR* | Represor | ATGACAAAGTTGCAGCCGAATACAGTGATCCGTGCCGCCCTGGACCTGTTGAACGAGGTC<br>GGCGTAGACGGTCTGACGACACGCAAACTGGCGGAACGGTTGGGGGTTCAGCAGCCGGC<br>GCTTTACTGGCACTTCAGGAACAAGCGGGCGCTGCTCGACGCACTGGCCGAAGCCATGCT<br>GGCGGAGAATCATACGCATTCGGTGCCGAGAGCCGACGACGACTGGCGCTCATTTCTGAT<br>CGGGAATGCCCGCAGCTTCAGGCAGGCGCTGCTCGCCTACCGCGATGGCGCGCGCATCCA<br>TGCCGGCACGCGACCGGGCGCACCGCAGATGGAAACGGCCGACGCGCAGCTTCGCTTCCT<br>CTGCGAGGCGGGTTTTTCGGCCGGGGACGCCGTCAATGCGCTGATGACAATCAGCTACTT<br>CACTGTTGGGGCCGTGCTTGAGGAGCAGGCCGGCGACAGCGATGCCGGCGAGCGCGGCG<br>GCACCGTTGAACAGGCTCCGCTCTCGCCGCTGTTGCGGGCCGCGATAGACGCCTTCGACG<br>AAGCCGGTCCGGACGCAGCGTTCGAGCAGGGACTCGCGGTGATTGTCGATGGATTGGCG<br>AAAAGGAGGCTCGTTGTCAGGAACGTTGAAGGACCGAGAAAGGGTGACGATTGA | 3 |
| *lacI* | Represor | ATGAAACCAGTAACGTTATACGATGTCGCAGAGTATGCCGGTGTCTCTTATCAGACCGTT<br>TCCCGCGTGGTGAACCAGGCCAGCCACGTTTCTGCGAAAACGCGGGAAAAAGTGGAAGC<br>GGCGATGGCGGAGCTGAATTACATTCCCAACCGCGTGGCACAACAACTGGCGGGCAAAC<br>AGTCGTTGCTGATTGGCGTTGCCACCTCCAGTCTGGCCCTGCACGCGCCGTCGCAAATTGT<br>CGCGGCGATTAAATCTCGCGCCGATCAACTGGGTGCCAGCGTGGTGGTGTCGATGGTAGA<br>ACGAAGCGGCGTCGAAGCCTGTAAAGCGGCGGTGCACAATCTTCTCGCGCAACGCGTCA<br>GTGGGCTGATCATTAACTATCCGCTGGATGACCAGGATGCCATTGCTGTGGAAGCTGCCT<br>GCACTAATGTTCCGGCGTTATTTCTTGATGTCTCTGACCAGACACCCATCAACAGTATTAT<br>TTTCTCCCATGAAGACGGTACGCGACTGGGCGTGGAGCATCTGGTCGCATTGGGTCACCA<br>GCAAATCGCGCTGTTAGCGGGCCCATTAAGTTCTGTCTCGGCGCGTCTGCGTCTGGCTGGC<br>TGGCATAAATATCTCACTCGCAATCAAATTCAGCCGATAGCGGAACGGGAAGGCGACTG<br>GAGTGCCATGTCCGGTTTTCAACAAACCATGCAAATGCTGAATGAGGGCATCGTTCCCAC<br>TGCGATGCTGGTTGCCAACGATCAGATGGCGCTGGGCGCAATGCGCGCCATTACCGAGTC<br>CGGGCTGCGCGTTGGTGCGGATATCTCGGTAGTGGGATACGACGATACCGAAGACAGCTC<br>ATGTTATATCCCGCCGTCAACCACCATCAAACAGGATTTTCGCCTGCTGGGGCAAACCAG<br>CGTGGACCGCTTGCTGCAACTCTCTCAGGGCCAGGCGGTGAAGGGCAATCAGCTGTTGCC<br>CGTCTCACTGGTGAAAAGAAAAACCACCCTGGCGCCCAATACGCAAACCGCCTCTCCCCG<br>CGCGTTGGCCGATTCATTAATGCAGCTGGCACGACAGGTTTCCCGACTGGAAAGCGGGCA<br>GTGA | 10 |
| *gfpmut2* | CDS | ATGAAAGGAGAAGAACTTTTCACTGGAGTTGTCCCAATTCTTGTTGAATTAGATGGTGAT<br>GTTAATGGGCACAAATTTTCTGTCAGTGGAGAGGGTGAAGGTGATGCAACATACGGAAA<br>ACTTACCCTTAAATTTATTTGCACTACTGGAAAACTACCTGTTCCATGGCCAACACTTGTC<br>ACTACTTTCGCGTATGGTCTTCAATGCTTTGCGAGATACCCAGATCATATGAAACAGCATG<br>ACTTTTTCAAGAGTGCCATGCCCGAAGGTTATGTACAGGAAAGAACTATATTTTTCAAAG<br>ATGACGGGAACTACAAGACACGTGCTGAAGTCAAGTTTGAAGGTGATACCCTTGTTAATA<br>GAATCGAGTTAAAAGGTATTGATTTTAAAGAAGATGGAAACATTCTTGGACACAAATTGG<br>AATACAACTATAACTCACACAATGTATACATCATGGCAGACAAACAAAAGAATGGAATC<br>AAAGTTAACTTCAAAATTAGACACAACATTGAAGATGGAAGCGTTCAACTAGCAGACCAT<br>TATCAACAAAATACTCCAATTGGCGATGGCCCTGTCCTTTTACCAGACAACCATTACCTGT<br>CCACACAATCTAAGCTTTCGAAAGATCCCAACGAAAAGAGAGACCACATGGTCCTTCTTG<br>AGTTTGTAACAGCTGCTGGGATTACACATGGCATGGATGAACTATACAAA | 11 |
| *gfpmut2x* | CDS | ATGAAAGGAGAAGAACTTTTCACTGGAGTTGTCCCAATTCTTGTTGAATTAGATGGTGAT<br>GTTAATGGGCACAAATTTTCTGTCAGTGGAGAGGGTGAAGGTGATGCAACATACGGAAA<br>ACTTACCCTTAAATTTATTTGCACTACTGGAAAACTACCTGTTCCATGGCCAACACTTGTC<br>ACTACTTTCGCGTATGGTCTTCAATGCTTTGCGAGATACCCAGATCATATGAAACAGCATG<br>ACTTTTTCAAGAGTGCCATGCCCGAAGGTTATGTACAGGAAAGAACTATATTTTTCAAAG<br>ATGACGGGAACTACAAGACACGTGCTGAAGTCAAGTTTGAAGGTGATACCCTTGTTAATA<br>GAATCGAGTTAAAAGGTATTGATTTTAAAGAAGATGGAAACATTCTTGGACACAAATTGG<br>AATACAACTATAACTCACACAATGTATACATCATGGCAGACAAACAAAAGAATGGAATC<br>AAAGTTAACTTCAAAATTAGACACAACATTGAAGATGGAAGCGTTCAACTAGCAGACCAT<br>TATCAACAAAATACTCCAATTGGCGATGGCCCTGTCCTTTTACCAGACAACCATTACCTGT<br>CCACACAATCTAAGCTTTCGAAAGATCCCAACGAAAAGAG**G**GACCACATGGTCCTTCTTG<br>AGTTTGTAACAGCTGCTGGGATTACACATGGCATGGATGAACTATACAAATACTAG | This work |
| BBa_B0010 | Terminator | TGAGGCATCAAATAAAACGAAAGGCTCAGTCGAAAGACTGGGCCTTTCGTTTTATCTGTT<br>GTTTGTCGGTGAACGCTCTC | 12 |

| | | | |
|---|---|---|---|
| T0 | Terminator | CTTGGACTCCTGTTGATAGATCCAGTAATGACCTCAGAACTCCATCTGGATTTGTTCAGAA CGCTCGGTTGCCGCCGGGCGTTTTTTATTGGTGAGAATCCAG | |
| lacI T | Terminator | TAACCGGGCAGGCCATGTCTGCCCGTATTTCG | |
| short T | Terminator | CGTCGAGACCCCTGTGGGTCTCGTTTTTT | 3 |
| T7 | Terminator | CTAGCATAACCCCTTGGGGCCTCTAAACGGGTCTTGAGGGGTTTTTTG | |

**Notes:** The -35 box, -10 box and the +1 start site are underlined. Operators are shown in bold. Annotations of the -35 and -10 region are based on the consensus hexanucleotides sequences reported by Moran et al.[13] Mutations are included in blue.
